# MEIsensor: a deep-learning method for mobile element insertion discovery

**DOI:** 10.64898/2026.03.25.714113

**Authors:** Yudong Wang, Pengyu Zhang, Shijie Wan, Zexin Zhang, Peisen Sun, Tun Xu, Peng Jia, Kai Ye, Xiaofei Yang

## Abstract

Mobile element insertions (MEIs) are a critical source of structural variation in the human genome, yet their accurate detection remains challenging, particularly within repetitive regions and for structurally complex insertions. While long-read sequencing enables the direct recovery of inserted sequences, current analytical pipelines rely heavily on repeat-library alignment, which can limit both computational efficiency and accuracy. Here, we present MEIsensor, a deep learning framework designed to detect and classify MEIs directly from long-read sequencing data. Unlike conventional methodologies, MEIsensor performs direct, sequence-based annotation of Alu, LINE1, and SVA insertions. Evaluated against HGSVC benchmarks using HiFi long-read datasets, MEIsensor consistently outperformed existing tools across major MEI classes while substantially improving computational efficiency, demonstrating particularly pronounced gains for structurally complex SVA insertions. Furthermore, manual curation confirmed that MEIsensor successfully identifies MEIs hidden in highly repetitive genomic regions, including centromeric higher-order repeats, that were previously absent from benchmark datasets. Together, these advancements establish MEIsensor as an efficient framework for MEI analysis that promises to significantly advance our understanding of human genomic architecture and evolution.

## Introduction

Mobile elements account for approximately 45% of the human genome^1^. Among them, retrotransposons including Alu, LINE1, and SVA remain active and generate novel insertions^2^. These mobile element insertions (MEIs) are a major class of structural variation (SV) and contribute to genome evolution and local regulatory variation^2^. Increasing evidence indicates that mobile elements harbor regulatory elements, such as promoters, enhancers, and transcription factor binding sites, which can influence gene expression, thereby affecting genome stability and disease susceptibility^3–5^, highlighting the need for accurate MEI detection and classification, particularly in repetitive and structurally complex regions such as centromeres.

Early MEI detection methods were largely developed for short-read data^6^, which often provide limited accuracy in repetitive regions. Recent advances in long-read sequencing (LRS) technologies, such as PacBio HiFi^7^ and Oxford Nanopore Technology sequencing^8^, have alleviated these limitations by spanning insertions and flanking regions, improving breakpoint resolution and sequence recovery. Despite these advances, accurate MEI type classification remains difficult because closely related elements often share substantial sequence similarity. Existing long-read pipelines, such as xTea^9^ and TLDR^10^, annotate candidate insertions by aligning sequences to reference repeat libraries, typically using tools such as RepeatMasker^11^. While effective in many scenarios, these approaches depend on predefined repeat libraries and sequence similarity for assignment. Because closely related MEI types share conserved domains and terminal motifs, library-based annotation can be ambiguous, especially in repetitive contexts.

This problem is further exacerbated by truncated or rearranged insertions and local repetitive motifs, which can obscure type-specific sequence signals. More recently, alignment-free strategies have been explored, exemplified by MEHunter^12^, which applies pre-trained large language models^13^ to annotate insertion sequences without explicit reference libraries. However, such large pre-trained models may incur higher false-positive rates and greater computational cost. Due to MEI type classification is a relatively focused sequence discrimination task, lighter task-specific models may be better suited to capture localized discriminative signals.

To address this challenge, we developed MEIsensor, a deep learning framework for MEI detection and classification from long-read sequencing (LRS) data. MEIsensor integrates candidate discovery with a lightweight convolutional neural network (CNN) equipped with residual connections^14, 15^, enabling direct sequence-based annotation without reliance on large repeat libraries. By tailoring the model architecture to the intrinsic sequence characteristics of LINE1, SVA, and Alu elements, MEIsensor achieves accurate and robust MEI classification across diverse insertion contexts while avoiding the limitations inherent to repeat-library based annotation pipelines.

We demonstrate that MEIsensor outperforms existing methods, *e.g.* xTea, MEHunter, and TLDR^9, 10, 12^ , in detection accuracy across multiple MEI types, with particularly strong performance on structurally complex insertions such as SVA elements that remain challenging for repeat-library dependent annotation. MEIsensor also maintains reliable performance in trio-based analyses and enables accurate detection of MEIs embedded within highly repetitive regions, including centromeric higher-order repeat (HOR) arrays.

## Results

### Overview of MEIsensor

MEIsensor operates in three steps. (1) We first identify candidate mobile element insertion (MEI) sites from long-read alignments against the reference genome by extracting insertion signals, *e.g.* soft-clipped reads and insertion operations in CIGAR string. (2) For each candidate locus, the inserted segment, such as clipped regions or insertion-containing read segments, are collected and summarized as candidate insertion sequences. (3) Each candidate insertion is classified into Alu, LINE1, or SVA using a trained deep learning model, and genotyped based on the relative supported reads at the locus.

To detect candidate MEI sites, we parse the alignments of long read and extract insertion signals, including soft-clipped reads, split alignments, and large insertion operations in CIGAR string (**Fig. 1a**). For each candidate locus, we extract the inserted sequence from the long reads. Specifically, the soft-clipped segments and insertion-containing read fragments are collected and summarized to generate candidate insertion sequences, which serve as the inputs for downstream feature extraction and MEI type classification.

**Fig. 1.**
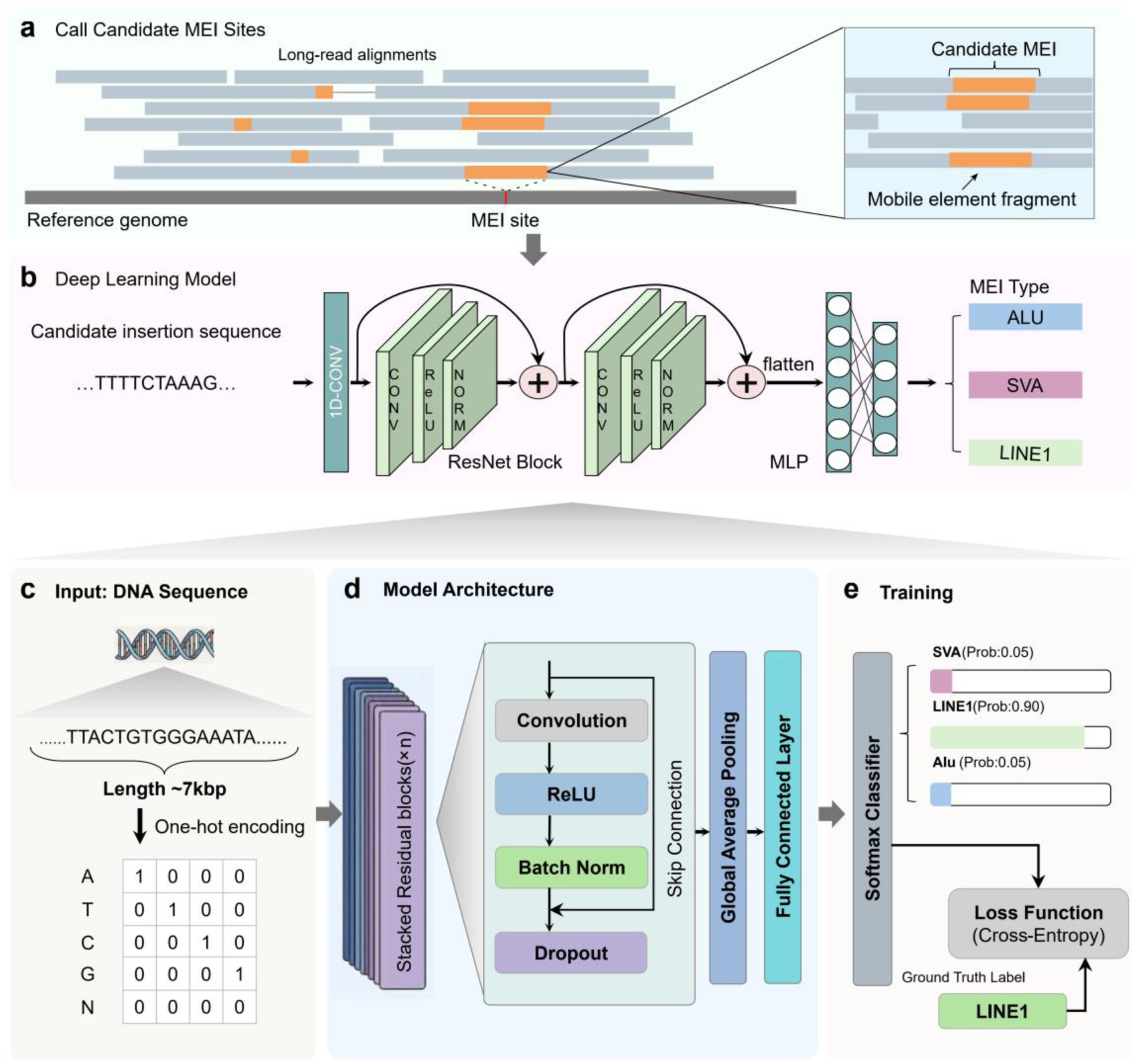
Overview of the MEIsensor framework. **a.** Candidate mobile element insertion (MEI) site identification. Candidate sites are identified using extracted insertion signals from long-read alignments, including soft-clipped reads, split alignments, and large insertion operations in CIGAR strings. **b.** Overview of the developed ResNet-based model. The local insertion sequence for each candidate site is extracted and analyzed by the model to categorize the insertion as an Alu, LINE1, or SVA element. **c.** Input sequence processing. A ∼7-kb genomic window centered on each candidate site is extracted, and its nucleotide sequence (A/C/G/T/N) is converted into a one-hot encoded matrix to serve as the model input. **d.** ResNet model architecture. The network comprises stacked one-dimensional convolutional residual blocks, followed by global average pooling and a fully connected layer for sequence feature extraction and classification. **e.** Model training. The framework is trained on known MEI types using a SoftMax classifier and cross-entropy loss, enabling the supervised learning of sequence patterns that distinguish between different MEI classes.

Deep learning models, especially convolutional neural network (CNN)^14^ and ResNet^15^, are well suited to address these challenges, arising from the intrinsic heterogeneity, structural variability, and frequent truncation of MEI insertion sequences that complicate type discrimination. MEI type classification is driven by multi-scale sequence signatures. At the local scale, MEI types exhibit characteristic motifs, such as poly(A) tails, target-site duplication (TSD) boundaries, and type-specific k-mer patterns. At broader scales, they differ in the organization and composition of sequence segments, including internal domains, truncated regions, and complex junction structures that extend beyond individual motifs^16^. CNNs naturally capture such hierarchical information by first learning short-range motif detectors and progressively integrating them into higher-order representations across increasing receptive fields, thereby modeling multi-scale sequence dependencies. Residual architectures further enhance this process by stabilizing optimization and preserving information flow, enabling deeper networks to model complex structural patterns without degradation. Motivated by these considerations, we developed a ResNet-based framework to extract informative features directly from candidate insertion sequences and perform MEI classification (**Fig. 1b**). This framework eliminates manual feature engineering and enables accurate and robust classification across diverse insertion contexts.

Specifically, we first padded the candidate insertion sequence to a fixed length, *e.g.* 7 Kb, and encoded as a one-hot A/C/G/T/N matrix to generate a standardized representation for downstream modeling (**Fig. 1c**). These representations are processed by an end-to-end deep learning architecture composed of one-dimensional convolutional layers with residual connections^15^ (**Fig. 1d**, **Supplementary Fig. 1**). Stacked residual blocks enable hierarchical feature extraction from raw sequence input, allowing the model to capture both local sequence motifs and longer-range contextual patterns. At the classification stage, MEIsensor applies a SoftMax classifier to assign each candidate insertion to a specific MEI type-Alu, LINE1, or SVA-yielding probabilistic predictions (**Fig. 1e**). The network is trained in a supervised manner using annotated insertion sequences derived from high-confidence MEI calls of 62 samples from Human Genome Structural Variation Consortium (HGSVC) v3 release^17^, with known type labels serving as ground truth (**Supplementary Table 1**). Encoded insertion sequences are provided as input, and model parameters are optimized by minimizing cross-entropy loss through iterative backpropagation. Model convergence and generalization are assessed on held-out validation data, ensuring robust type discrimination across diverse insertion contexts (**Supplementary Figure 2**).

By combining candidate discovery with deep learning-based sequence classification, MEIsensor provides a unified framework for detecting and classifying MEIs from long-read sequencing data. This design supports robust analysis across pedigrees, highly repetitive genomic regions, and large-scale population datasets.

### Performance evaluation for MEI detection

We evaluated MEIsensor using PacBio HiFi long-read sequencing data of HG00512 trio and CEPH1463 pedigrees generated by the HGSVC^17, 18^. We compared MEIsensor against three widely used state-of-the-art long-read MEI analysis tools, including xTea^9^, TLDR^10^, and MEHunter^12^. xTea and TLDR showed that long-read sequencing can recover full-length MEIs and complex insertion structures that are difficult to resolve with short reads, and both have been widely adopted in population-scale studies^17, 19^. MEHunter introduced a transformer-based approach for sequence-level MEI classification from long reads. Together, these methods represent distinct and influential strategies for long-read MEI analysis and serve as strong baselines for benchmarking MEIsensor in precision, recall, genotype inconsistency, and computational efficiency. (**Fig. 2**).

**Fig. 2.**
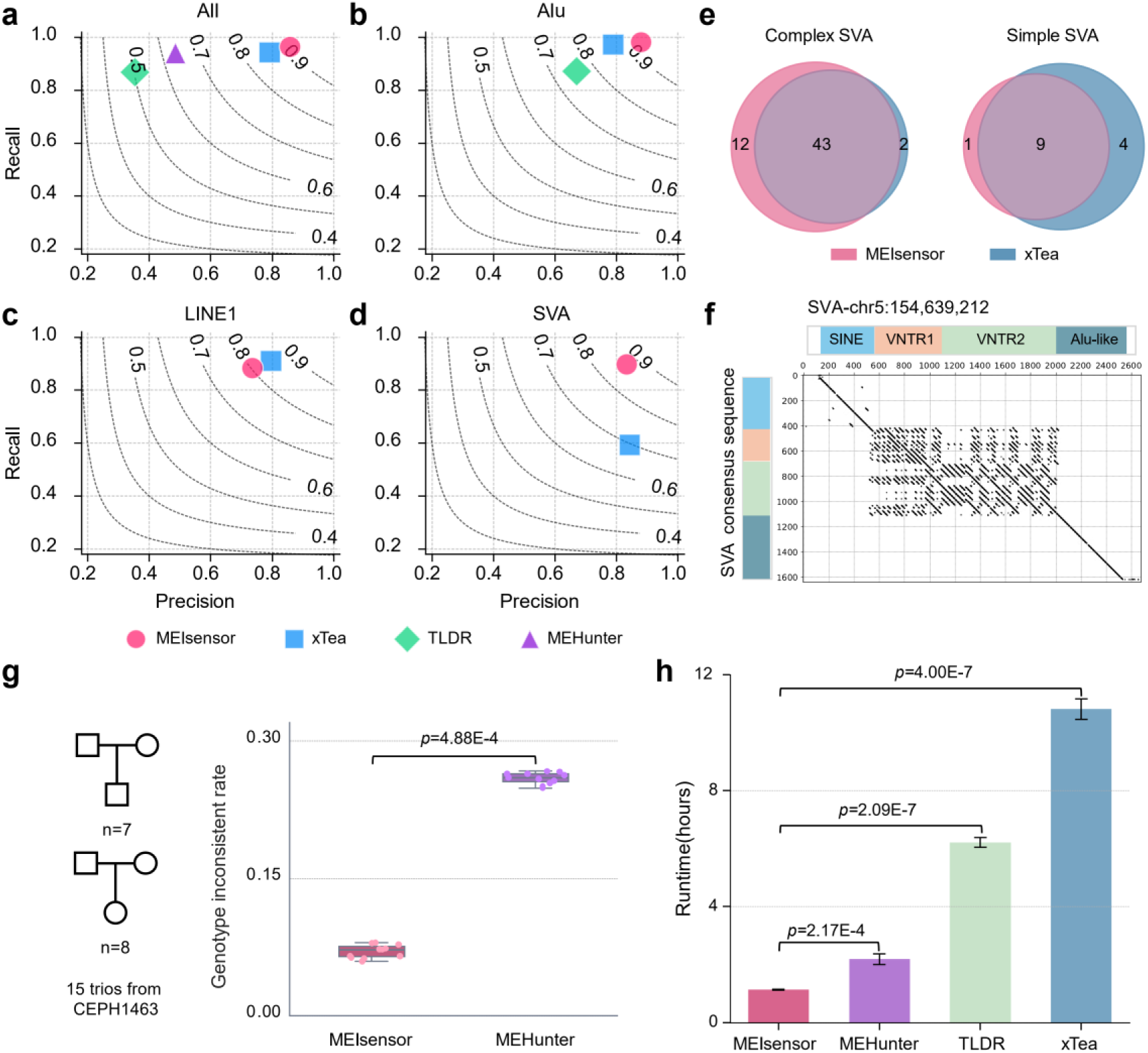
Performance evaluation of MEIsensor. a-d. Average detection performance of MEIsensor, xTea, TLDR, and MEHunter using HiFi long-read sequencing data from the HG00512, HG00513, and HG00514 trio. Panels display results for all MEIs (**a**), as well as specific Alu (**b**), LINE1 (**c**), and SVA (**d**) insertions. Each point represents the overall precision and recall under a unified filtering scheme, with dashed curves indicating iso-F1 contours. MEIsensor achieves a superior precision–recall balance for all MEIs combined, as well as for Alu and SVA insertions. For LINE1 insertions, xTea attains a slightly higher recall, whereas MEIsensor maintains higher precision. Note that MEHunter does not natively report MEI type annotations, and TLDR annotates only Alu insertions by default. **e.** Comparison of SVA insertions detected by MEIsensor and xTea in the HG00512 sample. Venn diagrams illustrate the overlap for complex (left) and simple (right) SVA insertions. The numbers indicate unique and shared events, demonstrating that MEIsensor detects numerous SVA insertions missed by xTea, particularly complex events. **f.** Dot plot visualization of a representative complex SVA insertion detected by MEIsensor but missed by xTea. The complex internal structure of the insertion highlights MEIsensor’s ability to resolve intact and structurally intricate SVA retrotransposition events. **g.** Genotype inconsistency assessment across 15 trios from the CEPH1463 cohort. Boxplots display Mendelian inconsistency rates at MEI sites for MEIsensor and MEHunter. MEIsensor exhibits significantly lower inconsistency rates (*p*-value = 4.88E-4, Wilcoxon rank-sum test), indicating more reliable genotyping. **h.** Computational efficiency comparison on the HG00512 HiFi dataset. Each tool was executed five independent times to assess runtime reproducibility.

We first evaluated MEIsensor on HG00512, HG00513, and HG00514 trio HiFi reads from HGSVC^18^. Benchmark datasets were constructed using curated non-reference MEI calls provided by HGSVC, which integrate multiple sequencing technologies and manual curation to serve as high-confidence ground truth (**Supplementary Tables 2-4**).

Detection performance was first assessed across different types of mobile elements. Averaged across the three benchmark samples, MEIsensor showed robust strong performance for both overall MEIs and specific MEI type (**Fig. 2a-d**, **Supplementary Fig. 3, Supplementary Table 5**). Overall, MEIsensor achieved a mean F1 score of approximately 0.91, with the highest recall (0.96) and precision (0.86) among four methods (**Fig. 2a**). In comparison, xTea reached a lower average F1 score (0.86), while TLDR and MEHunter showed substantially reduced F1 score of 0.50 and 0.64, respectively (**Fig. 2a**). Specifically, for Alu insertion, MEIsensor achieved the highest mean F1 score of 0.93, outperforming xTea (0.87) and TLDR (0.76) (**Fig. 2b**). For LINE1 insertions, MEIsensor achieved a mean F1 score of 0.80, slight lower than that of 0.85 for xTea (**Fig. 2c**). We have manually checked all putative false-positive detected LINE1 insertions by MEIsensor in HG00512 and found that more than half of them (18/33, 54.54%) are actual LINE1 insertion but absenting in the benchmark (**Supplementary Table 6, Supplementary Fig. 4**). In addition, four of them represented insertion events containing partial LINE1 sequence, while three events were annotated as LINE1 insertions with a few reads support and the remaining eight were primarily attributable to model misclassification (**Supplementary Table 7, Supplementary Data 1**). For SVA insertions, MEIsensor showed a clearer performance advantages achieving an average F1 score of 0.86, largely higher than that of 0.70 for xTea (**Fig. 2d**), reflecting improved sensitivity to this structurally complex MEI class and indicating that MEIsensor is able to learn and leverage sequence features associated with complex internal organization and structural variability of SVA insertions. All of these evaluation results indicate that MEIsensor provides more accurate and robust detection across different MEI categories, especially for the structurally complex SVAs.

Given the pronounced performance advantage observed for SVA detection, we performed a focused comparison of SVA insertions identified by MEIsensor and xTea in the HG00512 sample. We found that MEIsensor recovered 13 SVA insertion events that were missed by xTea, including 1 simple SVAs and a notably larger fraction (92.3%, 12/13) of complex SVA insertions (**Fig. 2e, Supplementary Table 8**). We found that these events are characterized by highly complex internal variable number tandem repeat (VNTR) structures (**Supplementary Fig. 5**), which pose challenges for repeat library alignment-based detection and classification. A representative example illustrates that MEIsensor can fully resolve a structurally complex and intact SVA insertion (chr5:154, 639, 212) that evade detection by xTea **(Fig. 2f**). In this case, the insertion exhibits a full-length SVA structure with characteristic internal components, including VNTR regions, embedded within a highly repetitive genomic context that poses substantial challenges for accurate detection. The identification of these complex SVAs highlight the sensitivity of MEIsensor to complete retrotransposition events even in structurally complex and repetitive regions.

Because MEIs frequently occur in repetitive and complex genomic regions, accurate genotyping remains intrinsically challenging and high-confidence ground truth genotypes are often unavailable. Trio-based inheritance patterns therefore provide an independent and biologically grounded criterion for assessing genotyping reliability. Accordingly, we evaluated genotyping consistency using pedigrees samples (including 15 trios) from the CEPH1463 cohort^20^ by assessing Mendelian inconsistency at detected insertion sites (**Fig. 2g, Supplementary Table 9**). MEIsensor exhibited significant lower genotype inconsistency rates than MEHunter across all pedigrees (mean inconsistency 0.07 vs. 0.26, *p*-value=4.88E-4, Wilcoxon signed-rank test), indicating reliable genotyping of MEIsensor.

Finally, we evaluated computational efficiency by executing each method on five independent runs on the HG00512 HiFi dataset (**Fig. 2h**). The tools differed substantially in resource usage. TLDR^10^ was run with four CPU cores; MEIsensor and MEHunter^12^ were executed using four CPU cores together with a single NVIDIA RTX 2080 Ti GPU; and xTea^9^ was configured to use 48 CPU cores. Despite this substantially higher CPU allocation, xTea required markedly longer runtimes (10.8h), followed by TLDR (6.2 h) and MEHunter (2.2 h). MEIsensor consistently achieved the shortest and most stable runtime (1.1h) under a more modest resource configuration, with significantly shorter runtimes than MEHunter (*p* = 2.17E-4), TLDR (*p* = 2.09E-7), and xTea (*p* = 4E-7) by Welch’s t-test.

### MEIsensor enables MEI detection in highly repetitive genomic region

To investigate the origin of apparent false-positive calls produced by MEIsensor, we systematically examined all putative false-positive MEIs detected in HG00512 through manual inspection using IGV^21^ and RepeatMasker annotation^11^. This rigorous manual curation revealed that a substantial number of these putative false positives (*n* = 34), comprising 18 LINE1s, 13 Alus, and 2 SVAs, corresponded to *bona fide* mobile element insertions. These insertions were embedded within highly repetitive and structurally complex genomic regions (**Fig. 3a, Supplementary Figs. 4, 6, and 7, Supplementary Tables 6 and 7**) and were missing from the original benchmark due to limitations in its construction^17^. A representative example is an insertion (chr1:13, 161, 043) contains two fragmented LINE1 with length of 2, 949 bp and 378 bp, respectively (**Fig. 3b**). Because the insertion locus resides within a highly repetitive genomic context, the mapping quality is significantly reduced, causing a substantial fraction of reads to fail contiguous mapping across the region. Additionally, we manually curated an ∼300 bp Alu insertion in 3q22.1 (chr3:130, 090, 881), which was surrounded by highly diverse sequences (**Supplementary Fig. 8**).

**Fig. 3.**
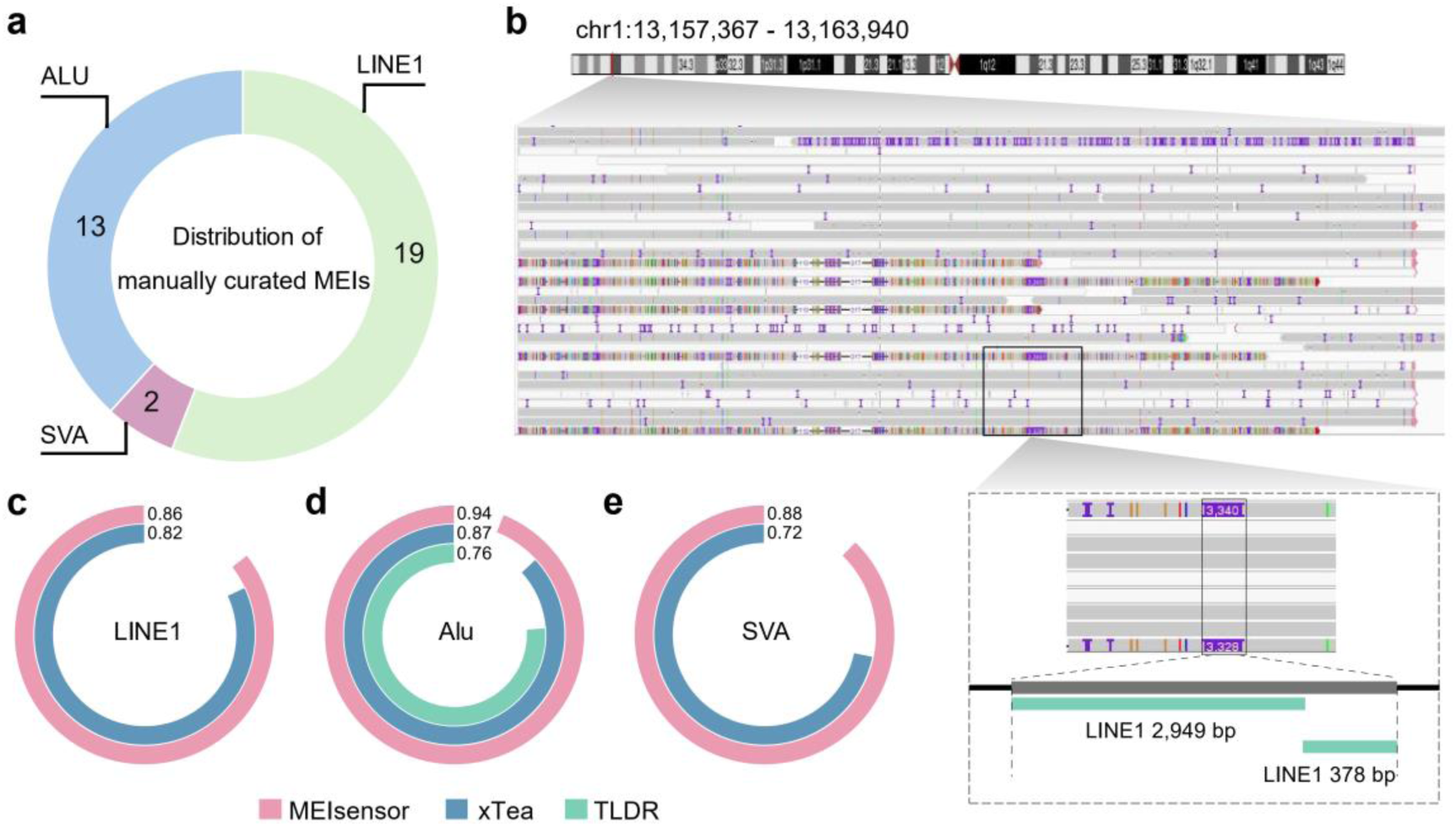
MEIsensor identifies *bona fide* MEIs in complex genomic regions missed by current benchmarks. **a.** Distribution of manually curated MEIs. The distribution of LINE1, Alu, and SVA insertions among the putative false-positive events in HG00512. Manual curation via IGV visualization and RepeatMasker annotation confirmed these as bona fide MEIs that were absent from the original benchmark due to their complex genomic contexts. **b.** Example of a *bona fide* LINE1 insertion. A genuine LINE1 insertion present in HG00512 but missing from the original benchmark dataset. The insertion resides in a structurally complex and highly repetitive region, which likely resulted in its prior exclusion. RepeatMasker annotations of this locus reveal two fragmented LINE1 elements (2, 949 bp and 378 bp in length), further validating the authenticity of the insertion event. **c-e.** Performance comparison of MEIsensor, xTea, and TLDR on the HG00512 sample using the updated benchmark that incorporates the 34 manually curated events. Results are shown for LINE1 (**c**), Alu (**d**), and SVA (**e**) insertions.

Based on these manual curation results, we constructed a high-confidence benchmark extension comprising these 33 events for HG00512. Re-evaluating detection performance against this updated benchmark led to a substantial improvement in performance metrics. Specifically, MEIsensor exhibited higher F1-scores for LINE1 (0.86), Alu (0.94), and SVA (0.88) insertions, outperforming xTea (0.82 for LINE1, 0.87 for Alu, and 0.72 for SVA) and TLDR (0.76 for Alu) (**Fig. 3c-e**). These findings indicate that the previously observed performance deficit for LINE1 insertions was primarily attributable to incomplete benchmark annotations rather than intrinsic limitations of MEIsensor. Collectively, these results demonstrate that MEIsensor not only provides robust and balanced detection across all major MEI types, but also offers a more faithful representation of the true insertion landscape by capturing events that are often omitted from incomplete benchmark sets due to complex genomic contexts.

### Detection of centromeric mobile element insertions by MEIsensor

Centromeric regions are enriched for repetitive sequences and complex SV^22, 23^ and are poorly represented in existing euchromatic region dominated MEI benchmarks. As long-read sequencing expands to trio-based and population-scale studies, comprehensive genome-wide MEI detection becomes essential. Evaluating MEI discovery within centromeric higher-order repeat (HOR) arrays therefore provides a stringent test of whether detection performance generalizes beyond well-curated regions to the most challenging parts of the genome.

Using long-read sequencing data from the CEPH1463 pedigree^20^, we examined MEIs within centromeric regions to assess the applicability and robustness of MEIsensor in highly repetitive genomic contexts. Across G3 and G4 individuals (*n* = 15), we totally detected 270 centromeric MEIs, including 167 ALU, 99 LINE1, and 4 SVA, and found most of them 94.8% (256/270) were located within α-satellite arrays (**Supplementary Fig. 9** and **Supplementary Table 10**). Specifically, for each sample, we found that an average of 15 MEIs located in α-satellite regions, whereas only 0 or 1 event was observed in β-satellite and HSAT regions, indicating α-satellite arrays being more active than β-satellite and HSAT arrays (**Fig. 4a**).

**Fig. 4.**
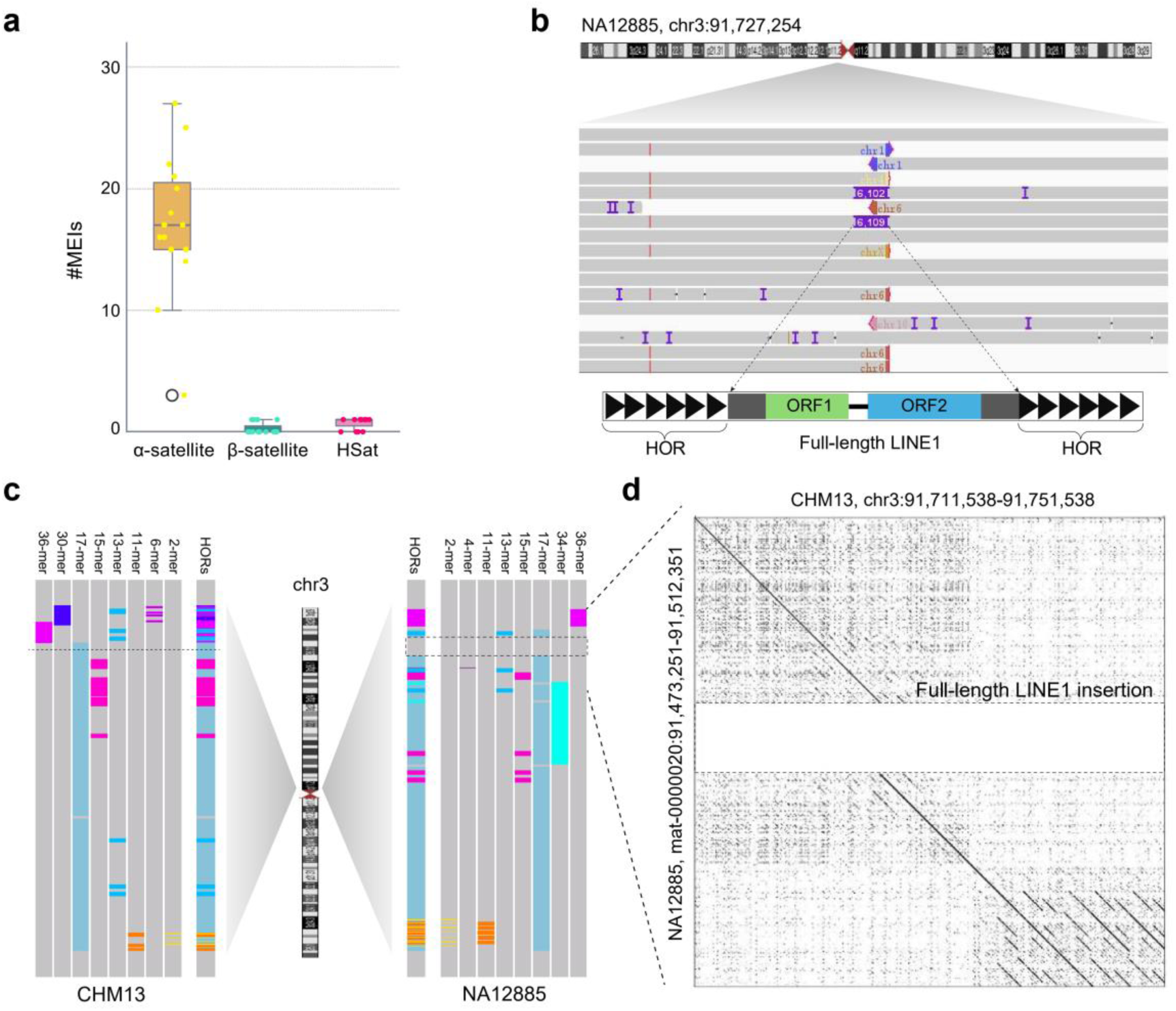
MEIsensor enables profiling of mobile element insertions within centromeric regions. **a.** Distribution of centromeric MEIs detected by MEIsensor using CHM13 as the reference genome. The number of MEIs detected across different centromere satellite classes (α-sat, β-sat, and HSAT) in G3 and G4 individuals from the CEPH1463 pedigree. **b.** Example of a centromeric LINE1 insertion. An ∼6.1 kb full-length LINE1 insertion identified by MEIsensor within the chromosome 3 centromeric region of sample NA12885. IGV visualization and centromere annotations demonstrate that the insertion resides deep within a higher-order repeat (HOR) array. **c.** A comparison of HiCAT-annotated HOR structures between NA12885 and the CHM13 reference at the chromosome 3 centromeric region. The alignment reveals that the LINE1 insertion occurs directly within the HOR array, disrupting a 17-mer HOR structure. **d.** A dot plot comparison between the CHM13 reference (*x*-axis) and the corresponding contig-level assembly of NA12885 (*y*-axis), providing orthogonal validation of a full-length LINE1 insertion at this highly repetitive locus.

In sample NA12885, MEIsensor identified a representative full-length LINE1 insertion of 6, 102 bp located at chr3:91, 727, 254, a locus within the centromere, together with centromere annotations, showed that the insertion is embedded within a HOR array, a region characterized by extreme sequence repetitiveness and structural complexity (**Fig. 4b**). Comparison of HOR organization between NA12885 and the CHM13 reference^24^ using HiCAT^25^ revealed that the LINE1 insertion disrupts a canonical 17-mer HOR unit in this region (**Fig. 4c**). Consistent with this observation, dotplot analysis between the corresponding CHM13 and NA12885 sequences revealed a marked interruption in sequence collinearity at the insertion locus, further demonstrating that MEIsensor resolves insertion-induced perturbations within centromeric HOR architecture (**Fig. 4d**).

Collectively, these analyses demonstrate that MEIsensor not only detects MEIs embedded within centromeric HOR arrays but also resolves their structural consequences on HOR organization. By extending MEI discovery into highly repetitive and structurally complex centromeric regions, MEIsensor broadens the applicability of genome-wide MEI detection beyond benchmarked euchromatic regions and supports its use in comprehensive long-read based population and pedigree analyses.

## Discussion

In this study, we show that MEIsensor is an effective long-read based framework for the discovery and genotype of MEIs, combining candidate discovery with deep-learning based sequence annotation. Our results suggest that MEIsensor effectively addresses two persistent challenges in MEI analysis, the computational burden of repeat library dependent annotation pipelines and the limited classification accuracy of existing approaches in complex genomic contexts. By replacing repeat-library matching^26, 27^ with a lightweight CNN for sequence annotation, MEIsensor achieves clear advantages in terms of detection accuracy, scalability, and computational efficiency across three major MEI classes, especially for the MEIs located in complex genomic regions and structurally complex MEIs

Across real HiFi long-read datasets, MEIsensor showed robust and well-balanced performance across all three major MEI classes while also recovering *bona fide* insertions that were absent from existing benchmark due to the highly repetitive and structurally complex genomic context. Together, these findings highlight the advantage of repeat-library-free, task-specific deep learning for MEI analysis. This advantage was particularly evident for SVA insertions, which showed the most pronounced improvement and remain especially challenging for conventional annotation strategies because of their composite structure and high internal complexity. The strong performance of MEIsensor on SVA is of particular importance given the established role of disease-associated SVA insertions in human disorders, including X-linked dystonia-parkinsonism and Fukuyama congenital muscular dystrophy^28–30^. These observations suggest that MEIsensor may provide a useful framework for future studies aimed at identifying and characterizing pathogenic SVA insertions in neurologic and other mobile-element–associated diseases.

A major strength of MEIsensor is its broad applicability across emerging long-read study designs, from pedigree-based analyses to population-scale cohorts. Its low Mendelian inconsistency rate supports reliable use in trio-based studies, whereas its favorable combination of speed and accuracy makes it well suited for large-scale population analyses. Beyond these conventional applications, MEIsensor also extends MEI analysis into structurally complex genomic regions such as centromeres, where conventional short-read approaches and repeat library dependent pipelines remain limited. As long-read pedigree and population datasets continue to expand, this capability may open new opportunities to investigate the evolutionary dynamics of human centromeres, including higher-order repeat (HOR) birth-and-death processes, as well as the inheritance and diversification of transposon insertions within these highly repetitive regions.

Despite these advances, several challenges remain. MEIsensor currently focuses on sequence-based classification of candidate insertions and does not explicitly model multi-breakpoint or nested insertion events. Moreover, as with other MEI detection methods, its performance ultimately depends on the quality of read alignment and candidate site discovery. Future work will focus on extending the framework to capture more complex insertion architectures, improving its integration with phased assemblies, and expanding training datasets to encompass a broader spectrum of mobile element diversity.

In summary, MEIsensor provides a practical and extensible framework for MEI detection and classification in long-read sequencing data. By enabling reliable analysis in repetitive and structurally complex regions, MEIsensor expands the scope of mobile element analysis and supports large-scale studies of genome evolution, inheritance, and disease.

## Methods

### Data preparation

All data used in this study were obtained from the MEI dataset provided by HGSVC^17, 18^, which is based on the human reference genome GRCh38. To avoid overfitting, MEI sequences from HG00512, HG00513, and HG00514 were held out as an independent benchmark set for performance evaluation, whereas the remaining samples were used for model development and training.

Raw HiFi reads from the HG00512 trio were aligned to the reference genome (GRCh38) using minimap2^31^, and the resulting alignment files were used as input for MEIsensor for candidate detection and MEI classification.

### MEI detection

MEIsensor processed insertion signals through a three-stage merging strategy following principles commonly used in long-read structural variant detection ^32^. MEIsensor first filtered the alignments using stringent quality criteria, including a minimum alignment length of 1 kb and a maximum number of split alignments of (3 + 0.1 \times) read length (in kb). The retained signals were then partitioned into 100-bp genomic bins, and nearby bins were iteratively merged into candidate insertion regions based on genomic proximity. Signals derived from the same read and from neighboring reads were subsequently consolidated to refine breakpoint localization, while candidate clusters with discordant insertion lengths were excluded using a maximum relative length difference threshold of 33%. For each resulting candidate site, the insertion-supporting sequence was extracted from the corresponding unaligned or soft-clipped read segment and used for downstream MEI annotation. This multi-step signal clustering and sequence extraction procedure generated high-confidence candidate insertions for classification.

### MEI classification

#### Model structure

We developed a convolutional neural network (CNN) to annotate mobile element (ME) insertions by analyzing insertion sequences encoded as one-hot vectors, where nucleotides (A, C, G, T/U, N) are represented as [1, 0, 0, 0], [0, 1, 0, 0], [0, 0, 1, 0], [0, 0, 0, 1], and [0, 0, 0, 0], respectively. The model generated four normalized probability scores (summing to 1) corresponding to Alu, SVA, LINE1, and non-MEI insertions. The architecture integrated three core components, (1) residual units combining batch normalization^33^, ReLU activation^34^, and convolutional layers to mitigate gradient vanishing/exploding issues; (2) adaptive pooling layers employing a dual-phase strategy—max pooling to amplify localized sequence motifs, followed by average pooling to integrate global contextual features; and (3) dense classifiers with linear and feed-forward networks for final prediction (**Supplementary Fig. 1**).

Residual connections were used to facilitate gradient propagation and stabilize model training. This hierarchical design resolves spatial patterns in MEI through biologically informed operations, that is max pooling prioritizes detection of critical motifs while average pooling captures broader sequence context. The architecture balances computational efficiency with interpretability, as feature extraction aligns with known ME insertion mechanisms.

#### Model training

We retrieved the Mobile Element Insertion (MEI) v1.0 dataset from HGSVC3, consisting of 62 human and cell line genomic samples. MEI sequences from 60 samples (excluding HG00512, HG00513, and HG00514) were allocated for model development, with the original distribution containing 8, 781 Alu, 1, 412 LINE1, 664 SVA, and 8, 000 non-MEI insertions. To mitigate class imbalance, we implemented resampling for LINE1 (1, 412→8, 472) and SVA (664→8, 632) insertions, resulting in a balanced training set of 8, 781 Alu, 8, 472 LINE1, 8, 632 SVA, and 8, 000 non-MEI insertions (**Supplementary Fig. 6**). MEI sequences from 60 samples were used for model development and were randomly split into training and test sets at an 8:2 ratio, whereas HG00512, HG00513, and HG00514 were held out as an independent validation cohort.

Our preprocessing approach involved three key steps to process MEI sequences. First, nucleotide sequences were converted into one-hot vectors, where each base (A, C, G, T, N) was encoded as [1, 0, 0, 0], [0, 1, 0, 0], [0, 0, 1, 0], [0, 0, 0, 1] and [0, 0, 0, 0], respectively, preserving base-level specificity. Next, sequences were uniformly padded with zeros to a fixed length of 7, 000 base pairs (bp), covering the full-size range of human MEIs according to HGSVC3 data. Terminal padding maintained original sequence structure while creating dimensionally consistent inputs, with flanking zeros preserving boundary context for positional feature extraction. Finally, categorical labels underwent one-hot encoding to represent MEI types, Alu ([1, 0, 0, 0]), LINE1 ([0, 1, 0, 0]), SVA ([0, 0, 1, 0]), and non-ME insertions ([0, 0, 0, 1]).

The processed one-hot sequences served as model inputs, containing detailed representations of MEI sequence features including local nucleotide patterns and global structural attributes. Corresponding one-hot labels functioned as training targets, enabling the model to establish precise sequence-to-class mappings for accurate MEI annotation.

The model was trained on an NVIDIA GeForce GTX 2080 Ti GPU for 40 epochs with a batch size of 64. We employed the Adam optimizer to minimize the categorical cross-entropy loss between predicted and ground-truth labels, using a learning rate of 0.000001 to ensure stable convergence. The model trained for the full 40 epochs was selected as the final version and evaluated on the test set. For comprehensive validation, we further applied this model to an independent cohort comprising samples HG00512, HG00513, and HG00514, which served as the basis for generating the results in Figure 2.

### MEIsensor evaluation

To comprehensively assess classification performance, we reported both class-specific and aggregate metrics. For each category (Alu, LINE1, SVA, and non-MEI), we calculated precision, recall, and F1-score based on standard definitions of true positives (correct predictions for the target class), false positives (samples incorrectly assigned to the target class), and false negatives (samples from the target class predicted as other classes).

To summarize overall model performance across classes, we used Micro-F1, which is computed from pooled true-positive, false-positive, and false-negative counts over all evaluated classes and reflects global predictive performance.

Class-specific precision, recall, and F1-score are shown in Fig. 2a-d, providing a detailed view of performance across MEI types, including rare classes such as SVA.

### Maximum-likelihood based genotyping

Genotypes were determined through a maximum-likelihood framework, which evaluates the probability of observing the sequencing data under each possible genotype hypothesis (0/0: homozygous reference; 0/1: heterozygous; 1/1: homozygous alternate). The genotype quality score (*GQ*) quantifies confidence in the called genotype and is derived from the likelihood ratio of the second most probable genotype to the selected genotype:

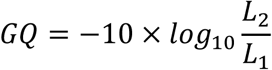

where *L1* and *L2* represent the likelihoods of the most likely (selected) and second most likely genotypes, respectively. This transformation converts likelihood ratios into Phred-scaled quality scores, where a *GQ* of 30 corresponds to a 1 in 1000 probability of genotyping error.

Genotype Likelihood Computation: Likelihoods for each genotype were modeled under a binomial distribution, considering the observed counts of variant (V) and reference (R) reads. For a diploid organism, the expected allele fractions are:

0/0 (homozygous reference): All reads expected to be reference (*p* = 0)

0/1 (heterozygous): Equal mix of reference and variant (*p* = 0.5)

1/1 (homozygous alternate): All reads expected to be variant (*p* = 1)

The binomial likelihood for genotype *g* is calculated as:

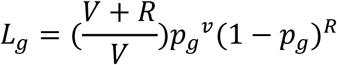

Error Modeling and Thresholds: To account for sequencing and alignment artifacts, we incorporated a genotype error rate (β) into the likelihood thresholds:

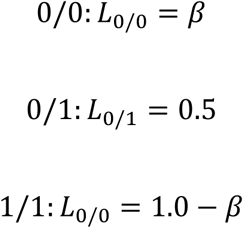

The default value *β*=0.05 reflects empirical observations of technical error rates in high-coverage WGS data (>30×), where 95% of reads at homozygous sites are expected to support the true allele. This conservative threshold minimizes false positives while accommodating occasional sequencing noise.

### Manually curated putative false positive MEI calls

Manual curation was based on the following two lines of evidence

1. Alignment-level evidence from IGV. For each candidate locus, long-read alignments were manually inspected in IGV^21^ to determine whether the observed pattern was consistent with a genuine insertion event. Specifically, we evaluated whether, (a) multiple reads showed concordant alignment signals near the same genomic position; (b) these breakpoint-associated signals were clustered consistently rather than being randomly distributed; (c) reads spanning the flanking regions on both sides of the candidate site supported the presence of an insertion; (d) despite local repetitiveness or reduced mappability, the overall read evidence remained reproducible, directionally consistent, and concentrated at the same breakpoint region. Candidate events showing these features were considered to have primary alignment support for a true insertion **(Supplementary Figs. 4, 6 and 7, Supplementary Data 1)**.
2. RepeatMasker annotation of the inserted sequence. For each candidate event, the extracted insertion sequence was further annotated with RepeatMasker^11^ to assess whether it contained sequence features consistent with the predicted MEI type. Specifically, we examined whether, (a) the insertion could be annotated as the expected repeat family, such as LINE1, Alu, or SVA; (b) the annotation covered a substantial portion of the inserted sequence rather than only a short local fragment; (c) the annotated structure was compatible with the characteristic organization of the corresponding element type, such as the main body of LINE1, the typical size and composition of Alu, or the composite structure of SVA; and (d) for truncated or rearranged insertions, RepeatMasker still provided sufficient evidence to support assignment to a specific mobile element family. When RepeatMasker produced a clear and coherent repeat annotation consistent with the predicted insertion type, this was regarded as sequence-level support for a *bona fide* MEI (**Supplementary Tables 6 and 7**).

Based on these criteria, a candidate event was classified as a benchmark-missing true insertion only when both stable alignment support and RepeatMasker annotation consistent with the expected MEI type were present.

### Mendelian inconsistency evaluation

To assess genotyping reliability in family-based analyses, we evaluated Mendelian inconsistency in the CEPH1463 pedigree using CHM13 as the reference genome. Fifteen parent–offspring trios from the pedigree were included in this analysis. For each detected MEI locus, genotypes generated by each method were compared across the three members of a trio to determine whether the observed inheritance pattern was consistent with Mendelian transmission.

Genotypes were represented as 0/0 (homozygous reference), 0/1 (heterozygous), or 1/1 (homozygous alternate). A locus was classified as Mendelian inconsistent when the offspring genotype was incompatible with the parental genotype combination under standard diploid inheritance. Specifically, inconsistency included cases such as, (1) both parents being 0/0 while the offspring was 0/1 or 1/1; (2) both parents being 1/1 while the offspring was 0/0 or 0/1; and (3) one parent being 0/0 and the other 1/1 while the offspring was not 0/1. All other parental-offspring genotype combinations were considered Mendelian consistent.

For each trio, the Mendelian inconsistency rate was calculated as the proportion of inconsistent MEI loci among all loci with non-missing genotype calls for all three individuals. Trio-level inconsistency rates were then summarized across the 15 trios and compared between MEIsensor and MEHunter to evaluate the robustness of MEI genotyping in pedigree data. xTea and TLDR were not included in this analysis because their output did not provide genotype information suitable for trio-based Mendelian inconsistency evaluation.

### Centromeric MEI analysis

Centromeric MEI analysis was conducted using CHM13 as the reference genome. Centromeric intervals were first identified according to RepeatMasker annotation of CHM13, which was used to define the boundaries of major centromeric repeat regions. To determine the chromosomal origin and genomic placement of assembled sequences, assembled contigs were aligned to CHM13 with minimap2. Contigs showing primary alignments to centromeric regions were retained for subsequent analysis.

For these centromere-associated contigs, higher-order repeat (HOR) organization was further annotated using HiCAT^25^ to resolve local centromeric repeat structure. This annotation enabled assignment of contig segments to specific satellite repeat classes and HOR units within each centromeric region. MEIs were then mapped onto these annotated centromeric intervals based on their genomic coordinates, and insertions overlapping centromeric repeat regions were extracted for downstream analyses.

## Data availability

The CHM13 data were downloaded from Telomere-to-telomere consortium (https://github.com/nanopore-wgs-consortium/CHM13). The GRCh38 reference genome was downloaded from the Genome Reference Consortium via NCBI (https://www.ncbi.nlm.nih.gov/grc/human). The training, test, and validation data were downloaded from the Human Genome Structural Variation Consortium phase 3 (HGSVC3) (https://www.internationalgenome.org/data-portal/data-collection/hgsvc3), and showed in **Supplementary Tables 1-4**. The raw sequencing data and assembly results for the CEPH1463 pedigree were downloaded from the AWS Open Data program (s3://platinum-pedigree-data/) or the European Nucleotide Archive (ENA; BioProject: PRJEB86317).

## Code availability

Source code for MEIsensor is available for download at https://github.com/xjtu-omics/MEIsensor. The repository is free for noncommercial use by academic, government and nonprofit/not-for-profit institutions.

## Acknowledgements

This work is supported by the National Key R&D Program of China (grant nos. 2025YFC3410300, 2022YFC3400300), the National Natural Science Foundation of China (grant nos. 32422019, 32430017, 32125009, 32400509, 325B2021), the Natural Science Foundation of Shaanxi Province (grant no. 2024JC-JCQN-28), the Fundamental and Interdisciplinary Disciplines Breakthrough Plan of the Ministry of Education of China (JYB2025XDXM101), and the Fundamental Research Funds for the Central Universities (xzy012024088).

## Author contributions

X.Y. and K.Y. conceived and supervised the study. X.Y and Y.W. designed the model, performed data analyses, and wrote the manuscript. P.Z. contributed to the evaluation of MEIsensor results and downstream analyses. S.W. and Z.Z. performed centromeric region annotation. P.S., T.X., P.J., and X.Y. contributed to data analysis, result interpretation, and manuscript revision. All authors read and approved the final manuscript.

