## supplementary tables1-10, supplementary figures1-9, supplementary data1 for "MEIsensor: a deep-learning method for mobile element insertion discovery": supplementary_data1.pdf

### IGV screenshots of false-positive MEIs in HG00512

#### LINE1

LINE1-HG00512-chr11:38,791,107; (misclassification by MEIsensor)

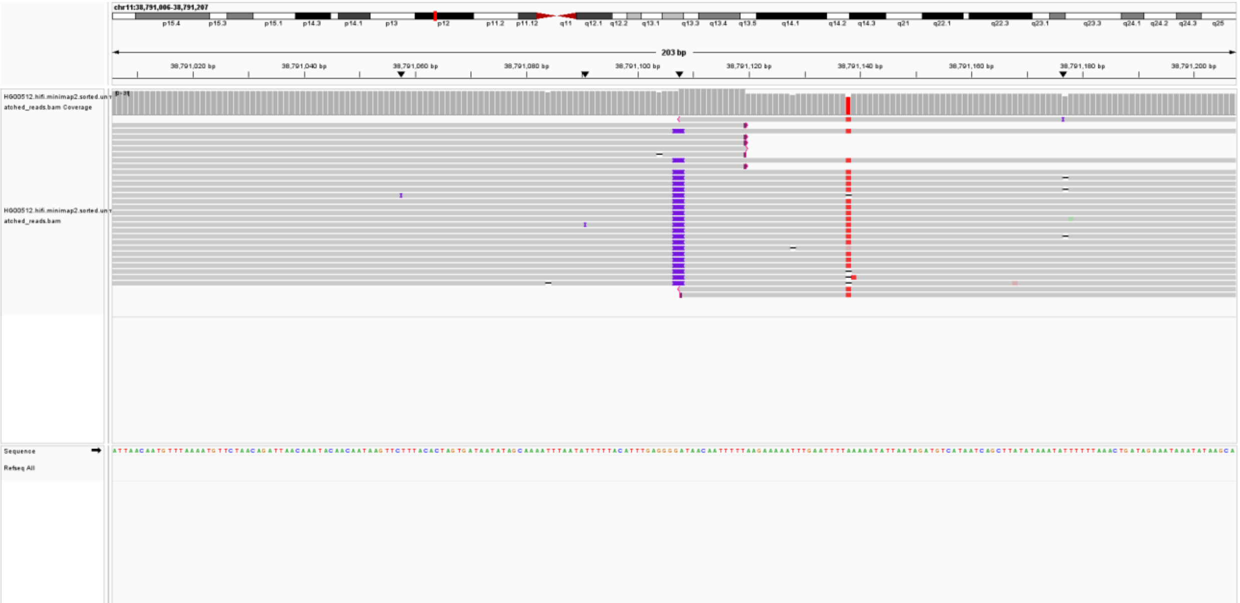

LINE1-HG00512-chr11:68,428,943; (misclassification by MEIsensor)

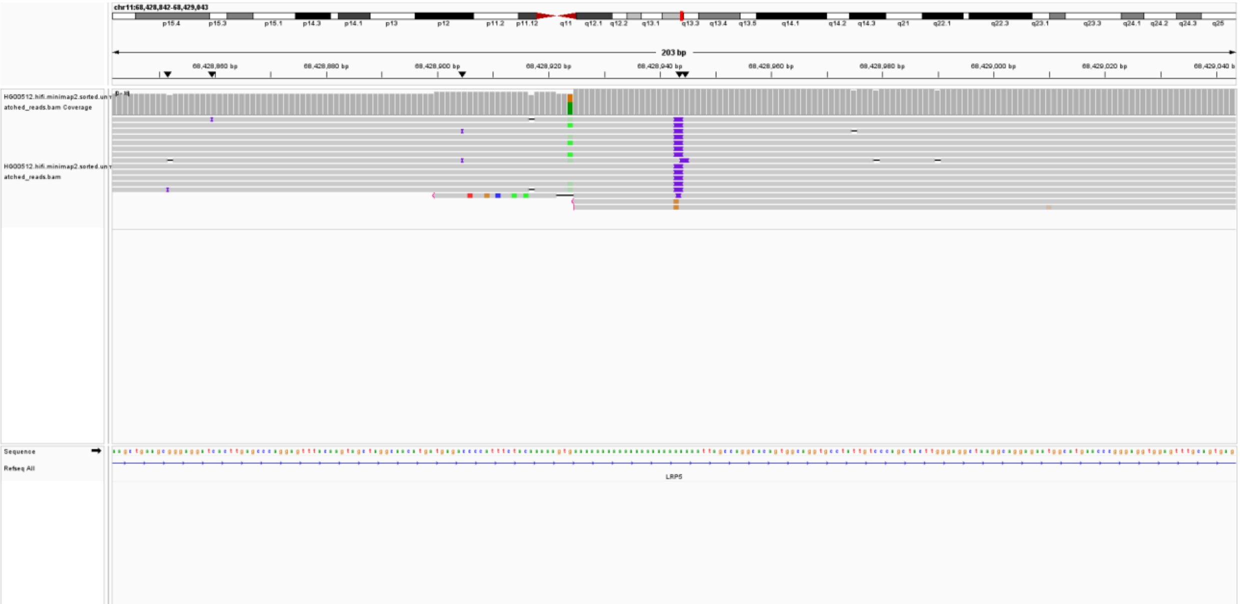

LINE1-HG00512-chr1:122,358,623; (lack reads support)

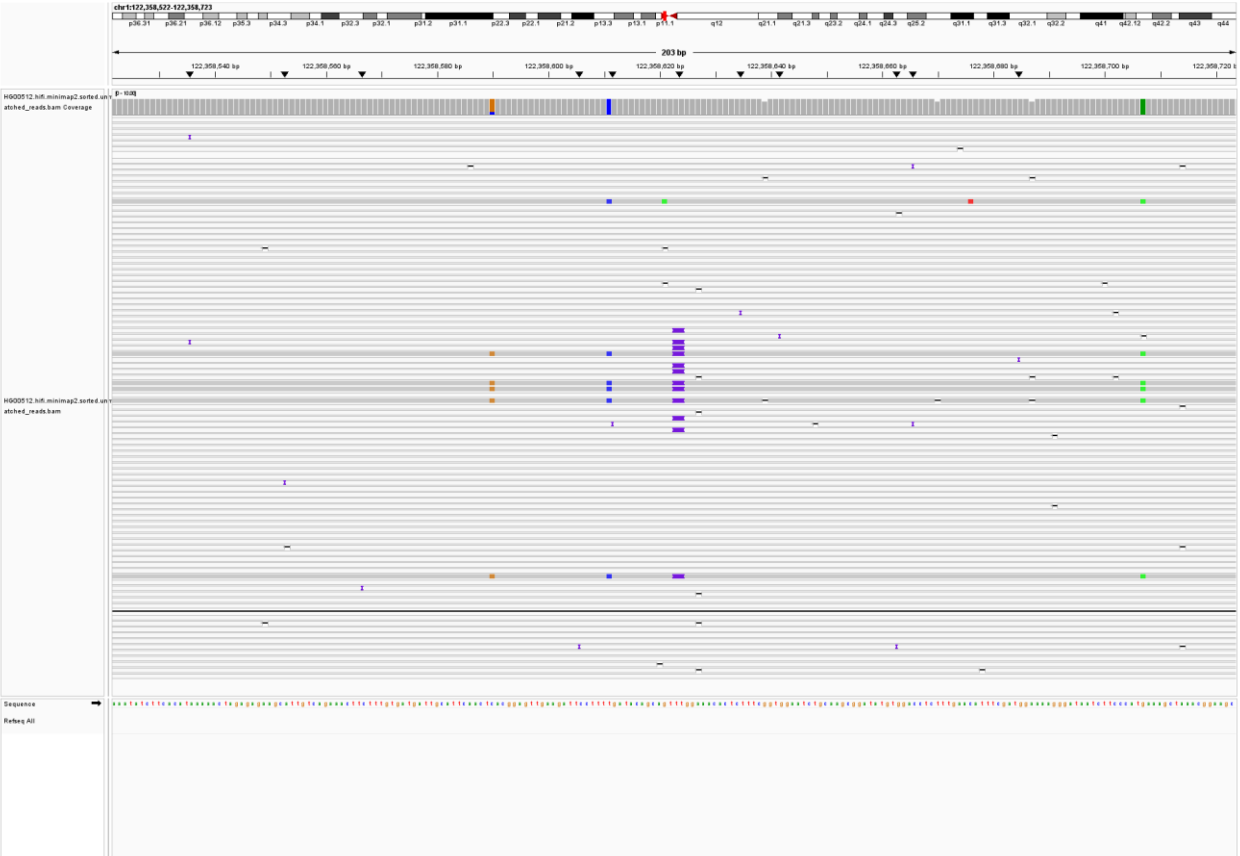

LINE1-HG00512-chr1:13,111,104; (lack reads support)

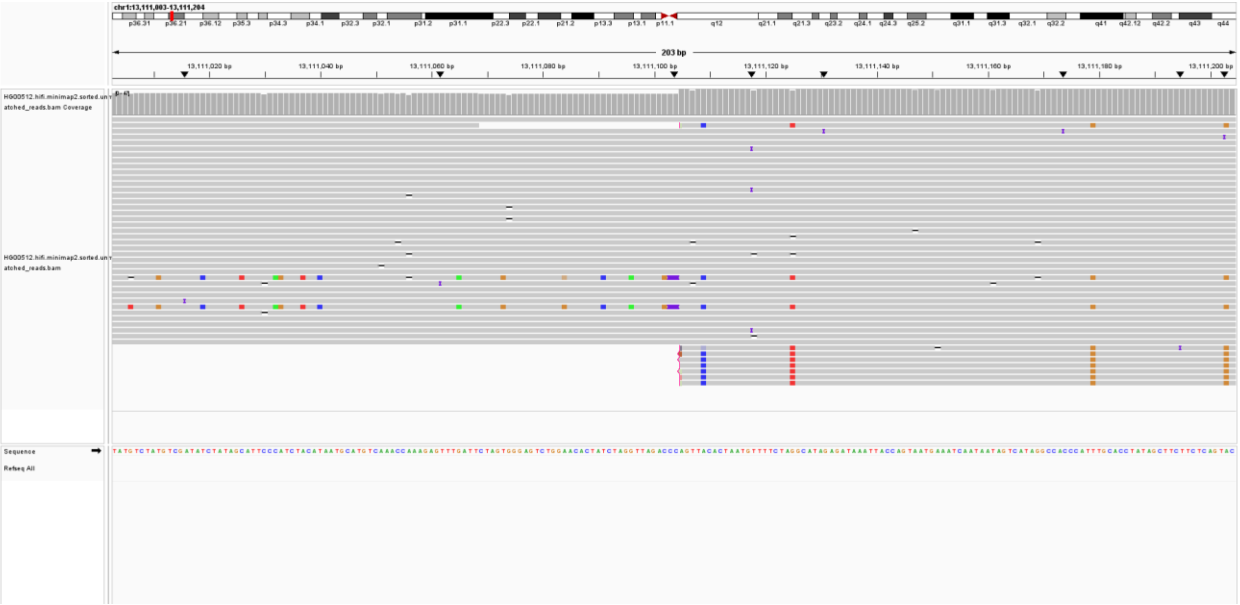

LINE1-HG00512-chr1:45,703,341; (misclassification by MEIsensor)

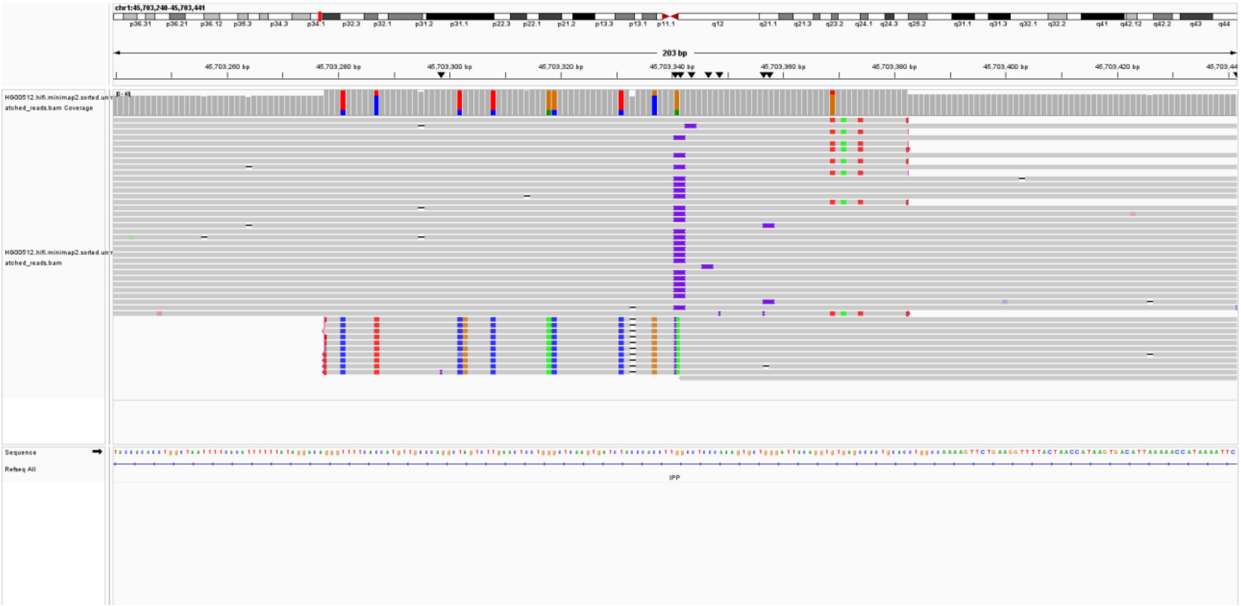

LINE1-HG00512-chr2:120,391,653; (misclassification by MEIsensor)

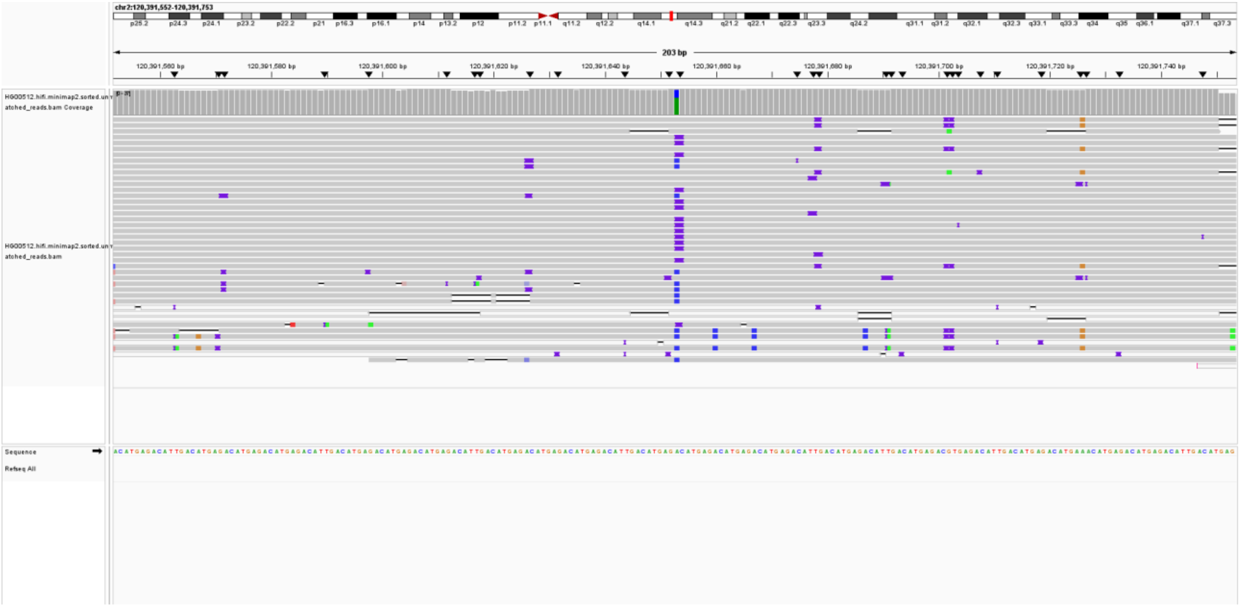

LINE1-HG00512-chr2:205,345,784; (misclassification by MEIsensor)

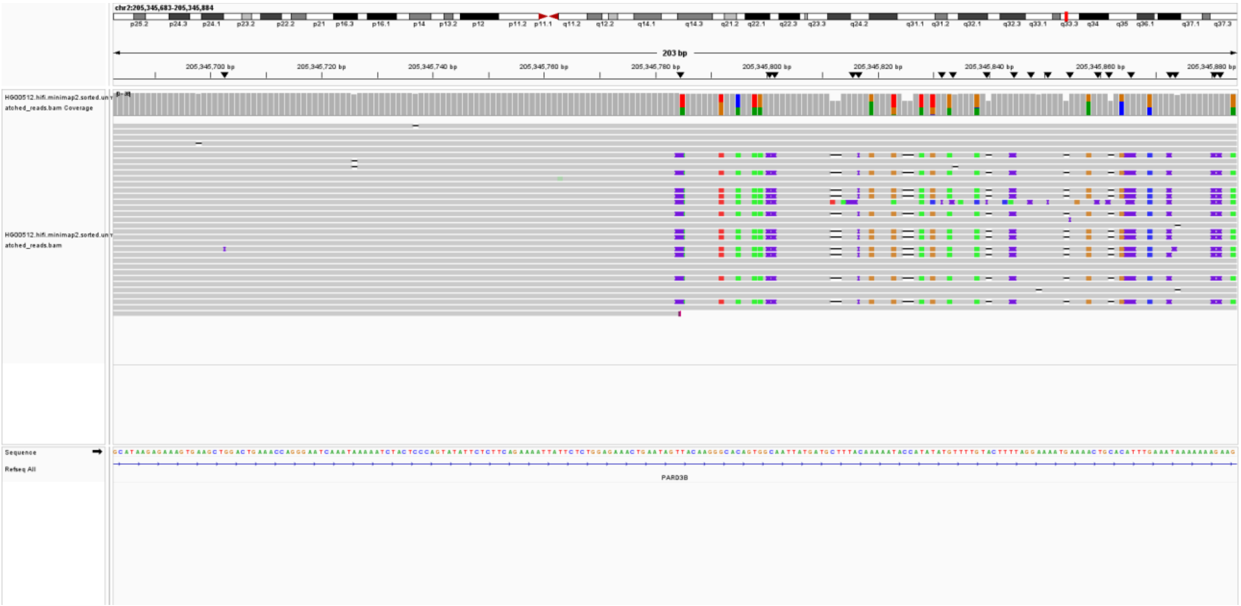

LINE1-HG00512-chr3:32,674,902; (misclassification by MEIsensor)

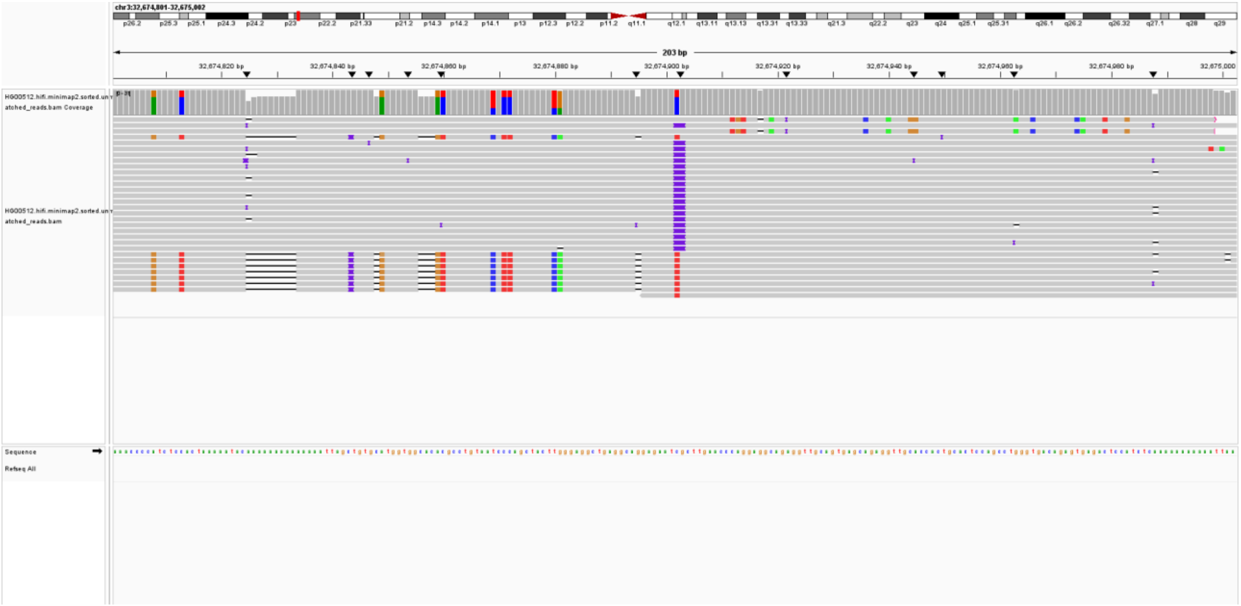

LINE1-HG00512-chr3:58,786,706; (misclassification by MEIsensor)

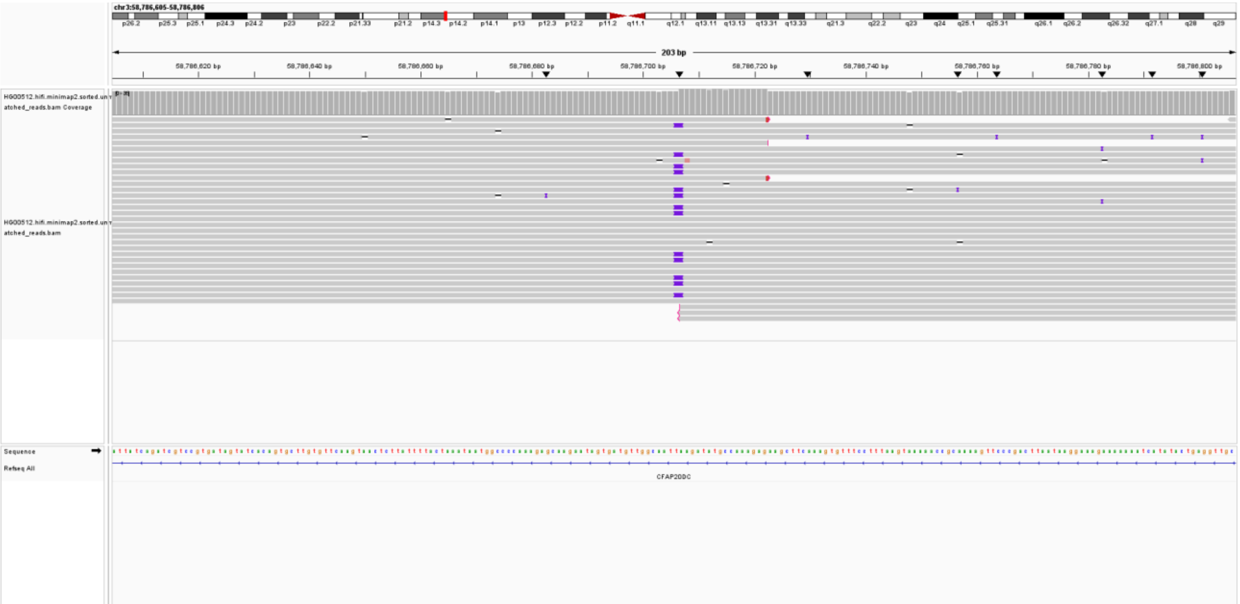

LINE1-HG00512-chr5:157,778,905; (lack reads support)

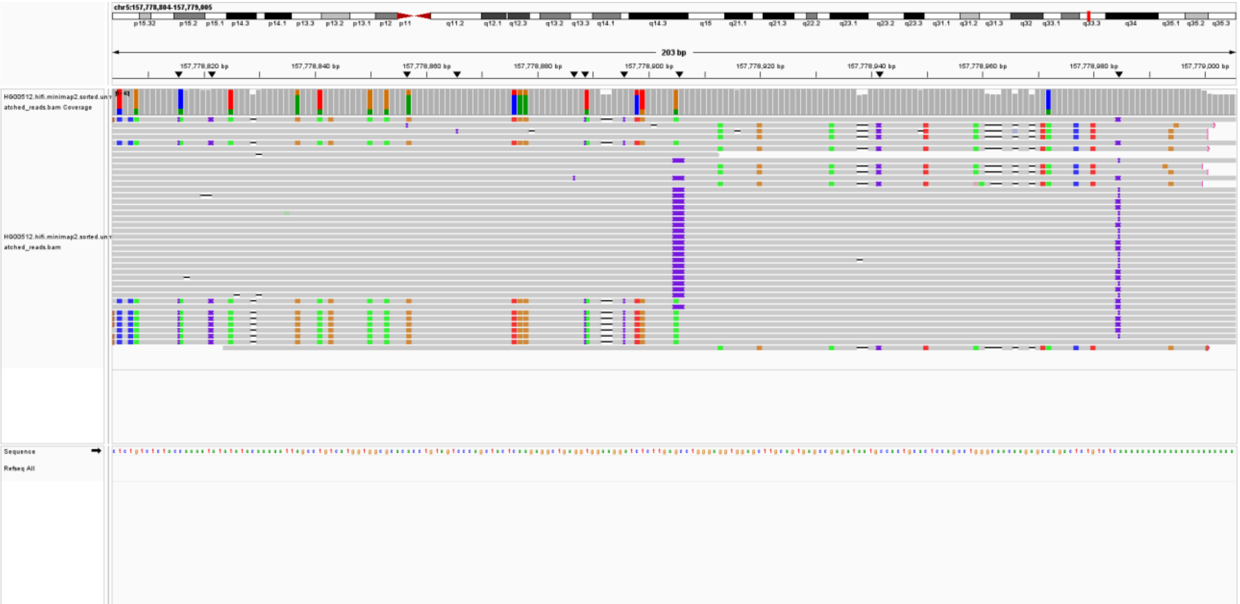

LINE1-HG00512-chr5:46,694,429; (misclassification by MEIsensor)

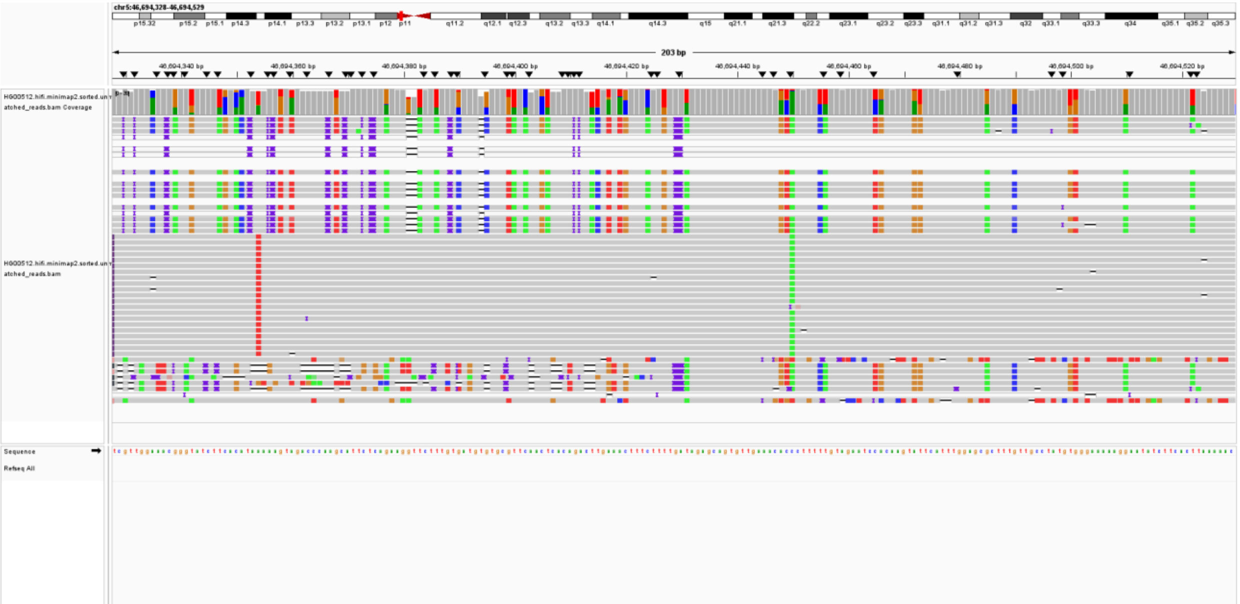

LINE1-HG00512-chr6:149,983,904; (misclassification by MEIsensor)

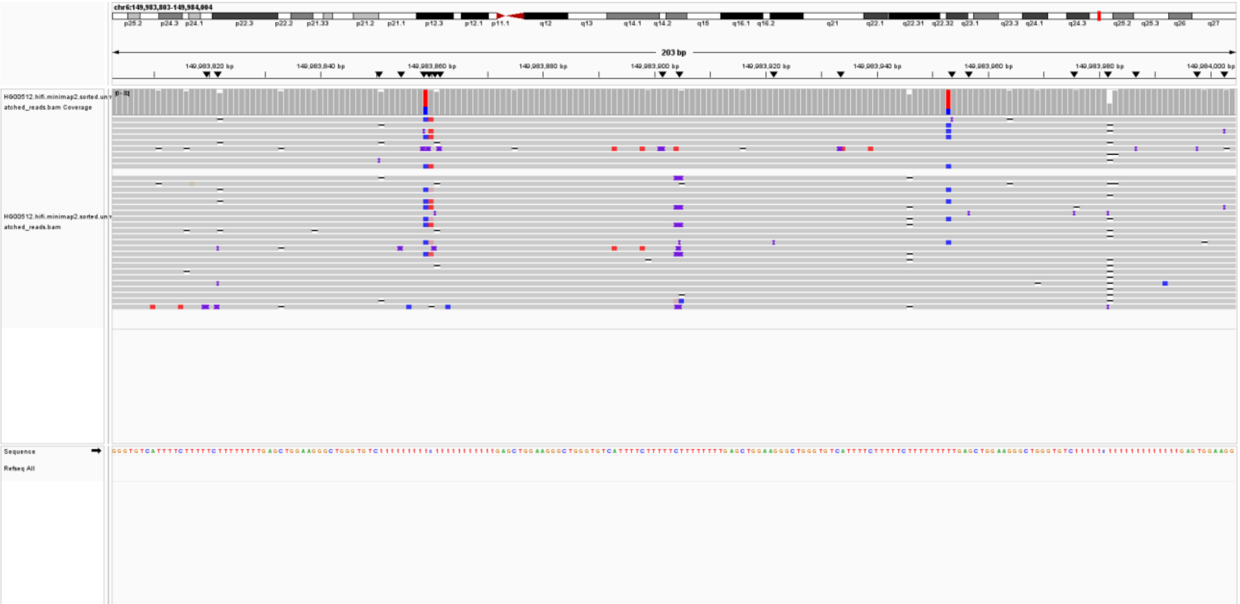

LINE1-HG00512-chr7:80,006,050; (misclassification by MEIsensor)

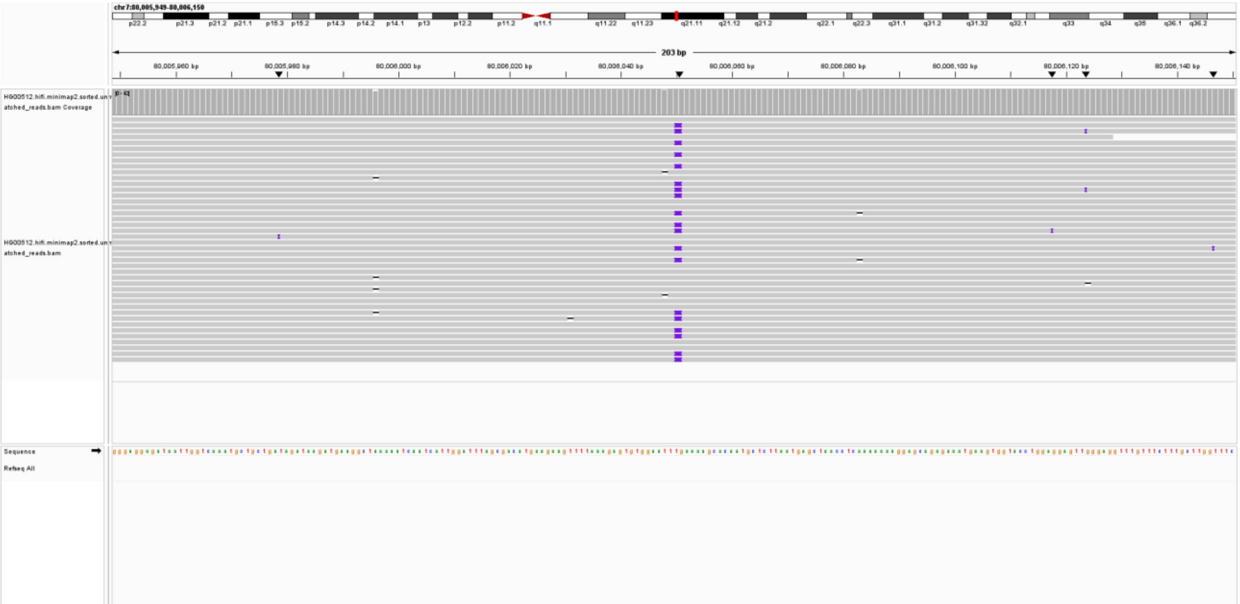

LINE1-HG00512-chrX:322,517; (misclassification by MEIsensor)

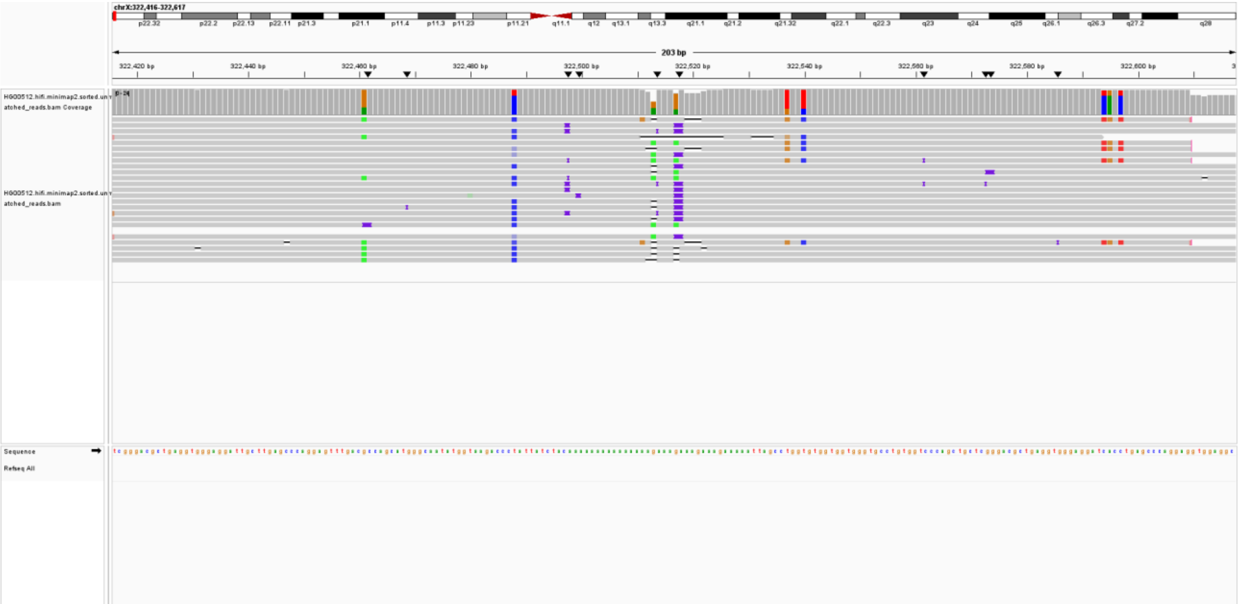

LINE1-HG00512-chrX:5,047,636; (misclassification by MEIsensor)

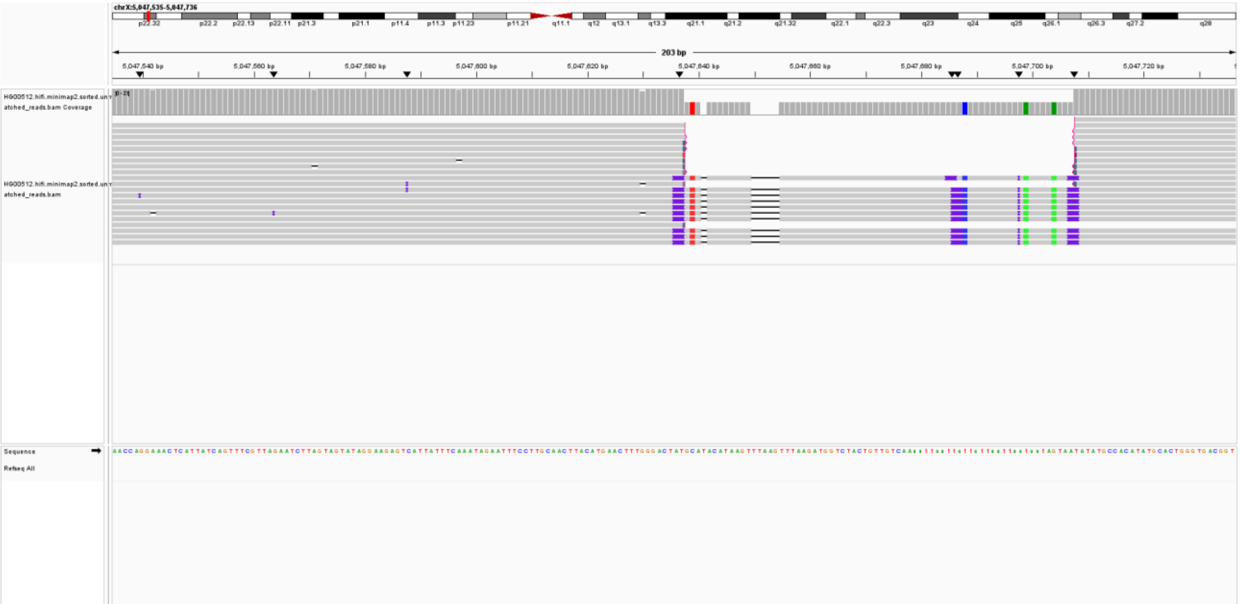

### ALU

ALU-HG00512-chr10:133,761,992; (lack reads support)

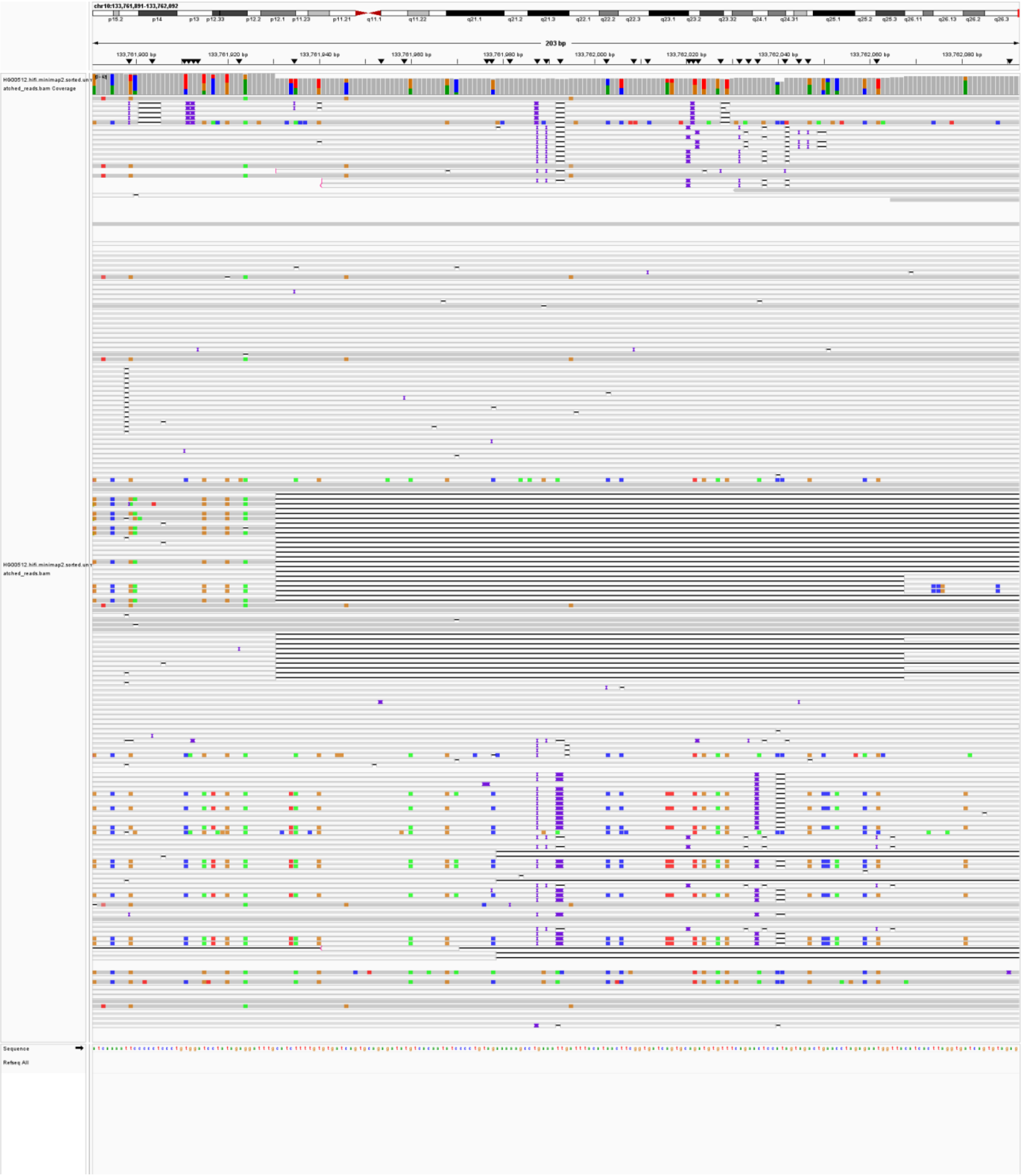

ALU-HG00512-chr10:15,400,028; (lack reads support)

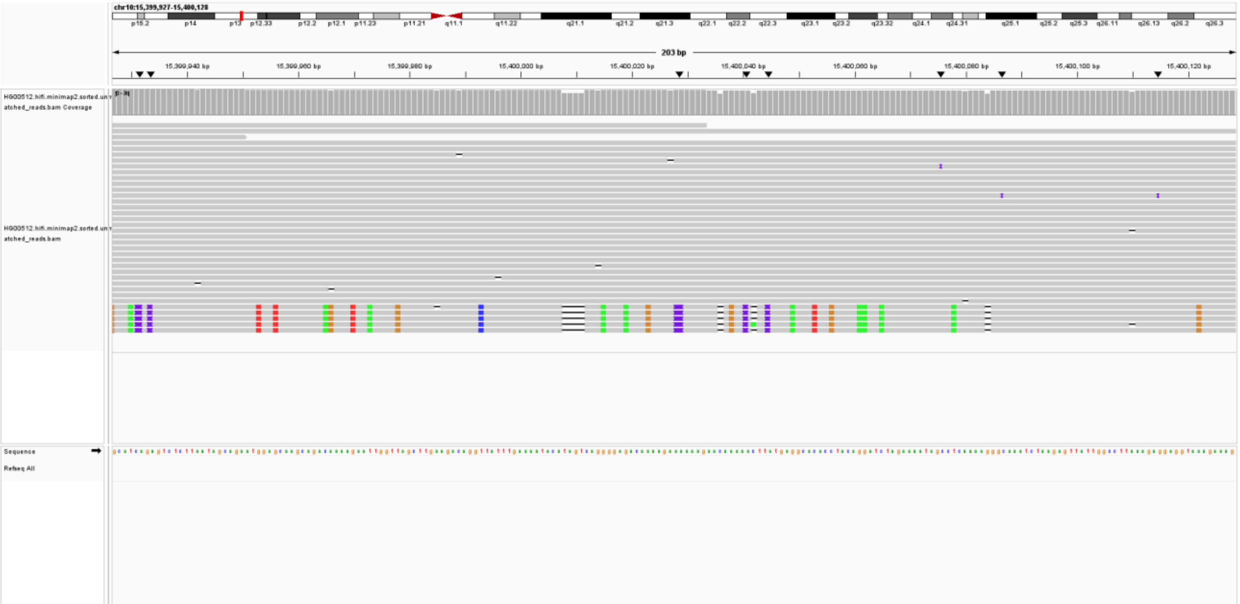

ALU-HG00512-chr10:26,627,268; (lack reads support)

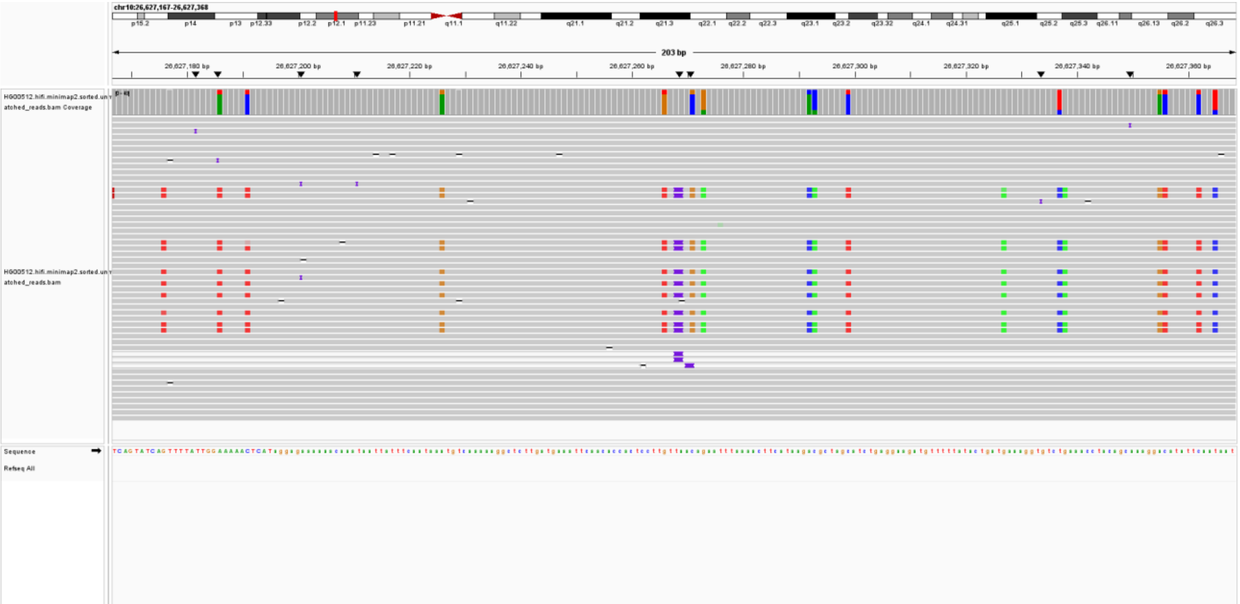

ALU-HG00512-chr10:26,938,142; (lack reads support)

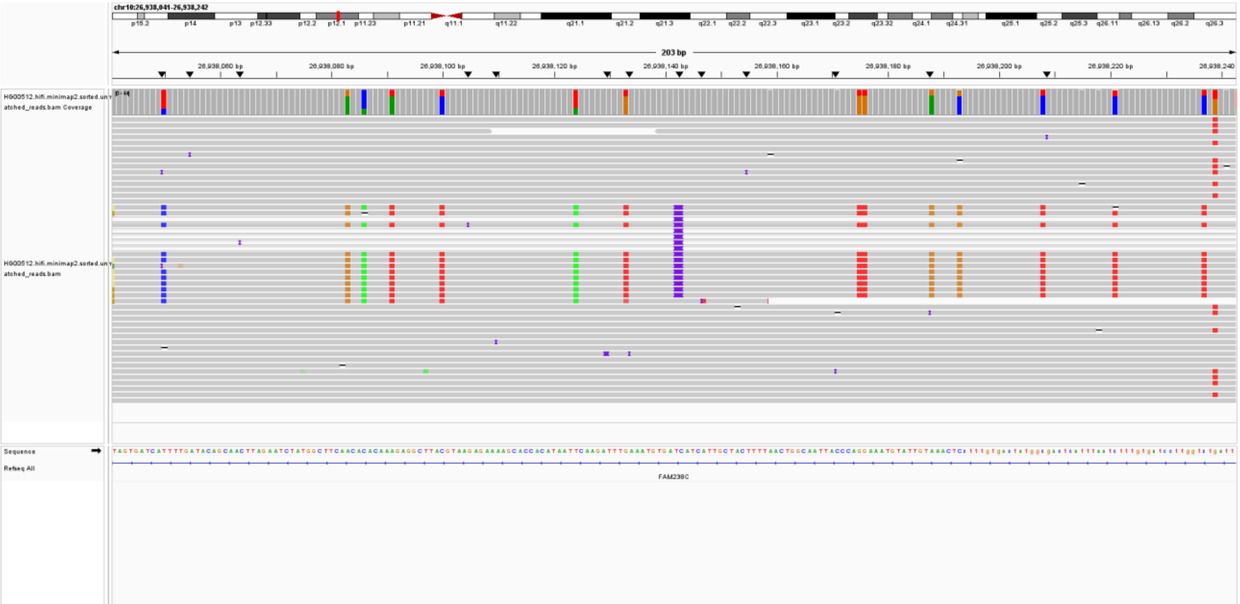

ALU-HG00512-chr10:27,331,443; (lack reads support)

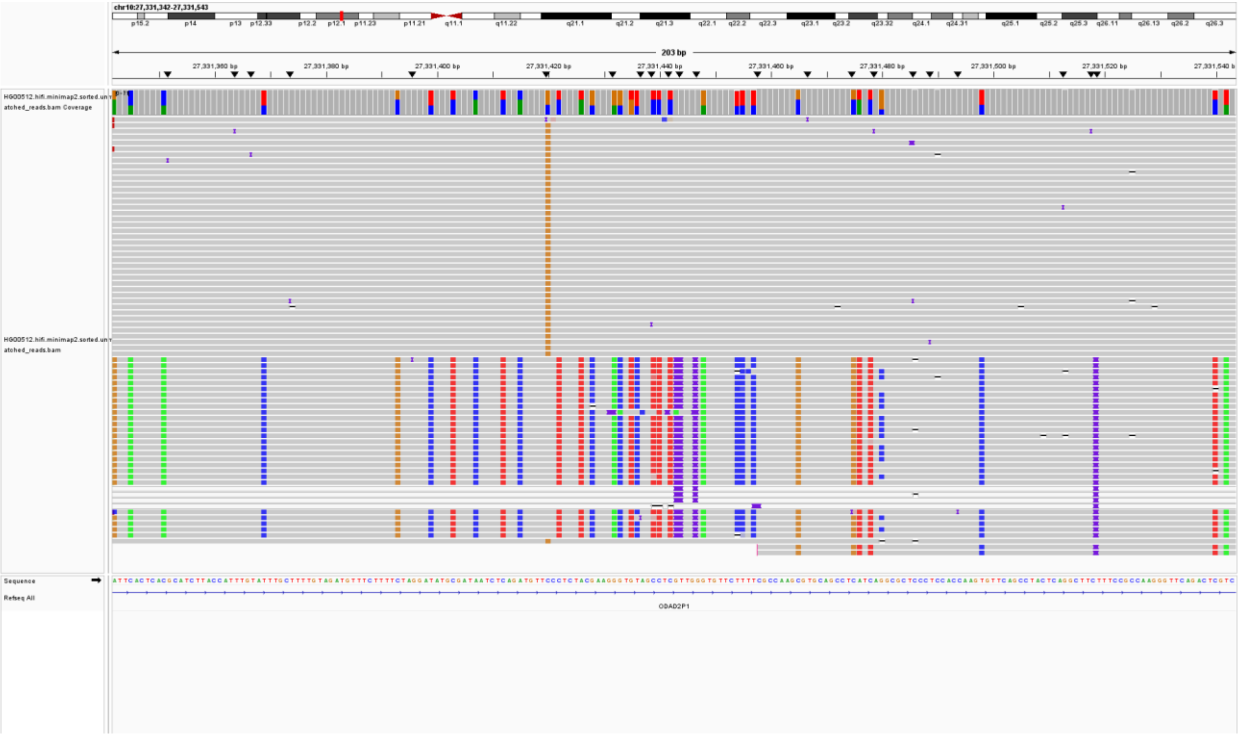

ALU-HG00512-chr13:27,215,551; (lack reads support)

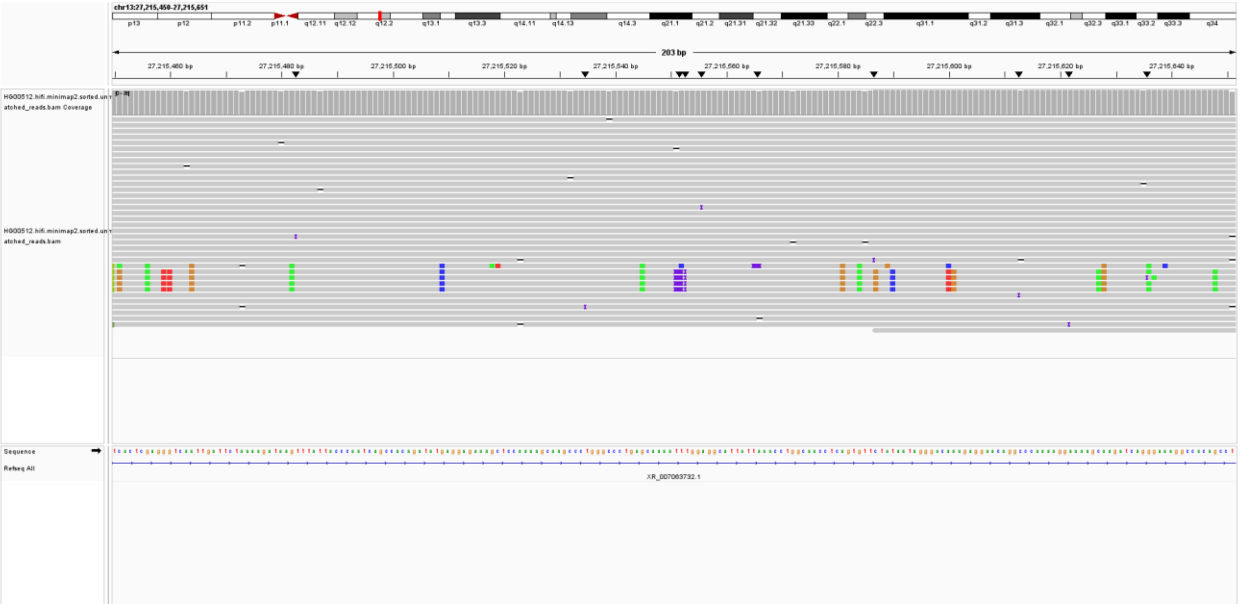

ALU-HG00512-chr13:41,485,435; (lack reads support)

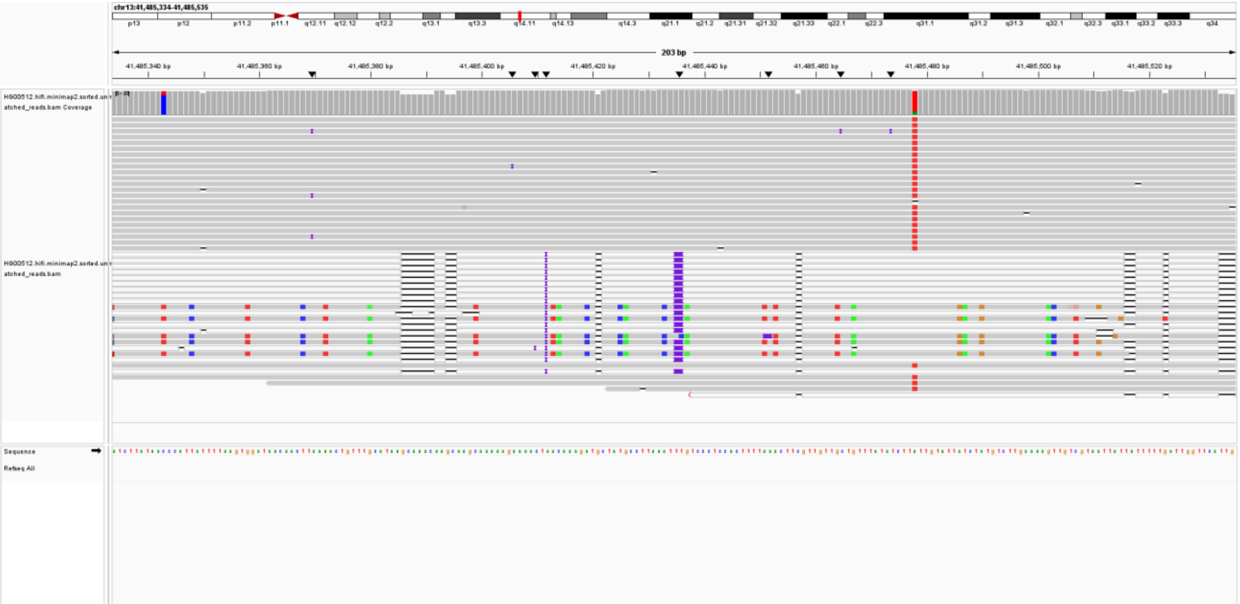

ALU-HG00512-chr15:90,671,366; (misclassification by MEIsensor)

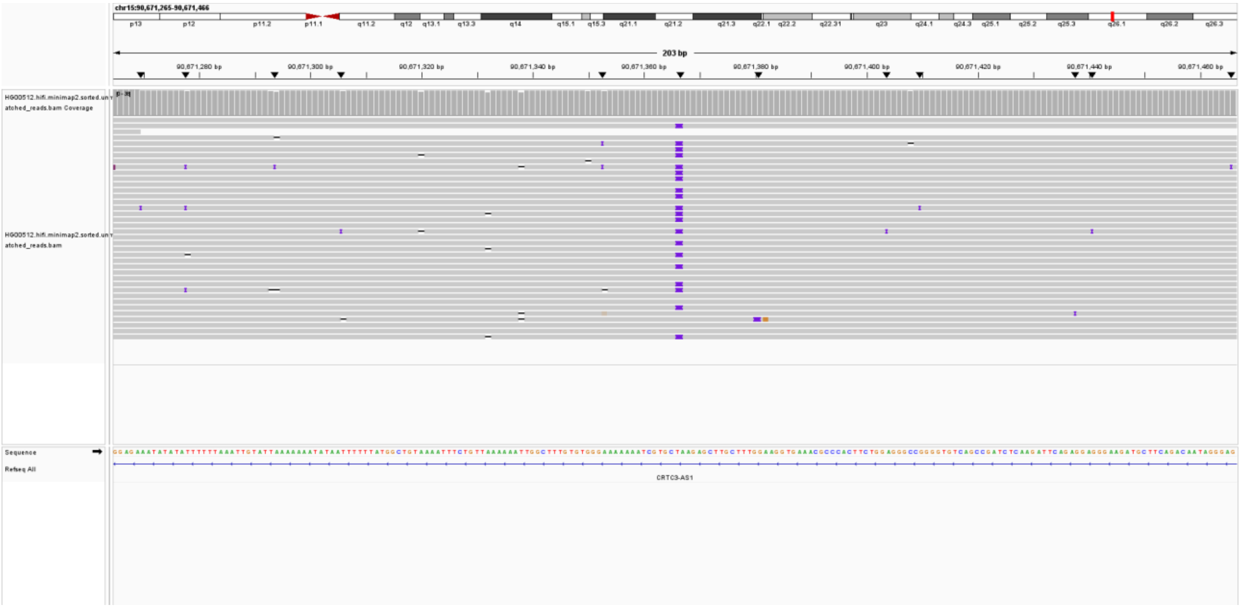

ALU-HG00512-chr20:29,202,607; (lack reads support)

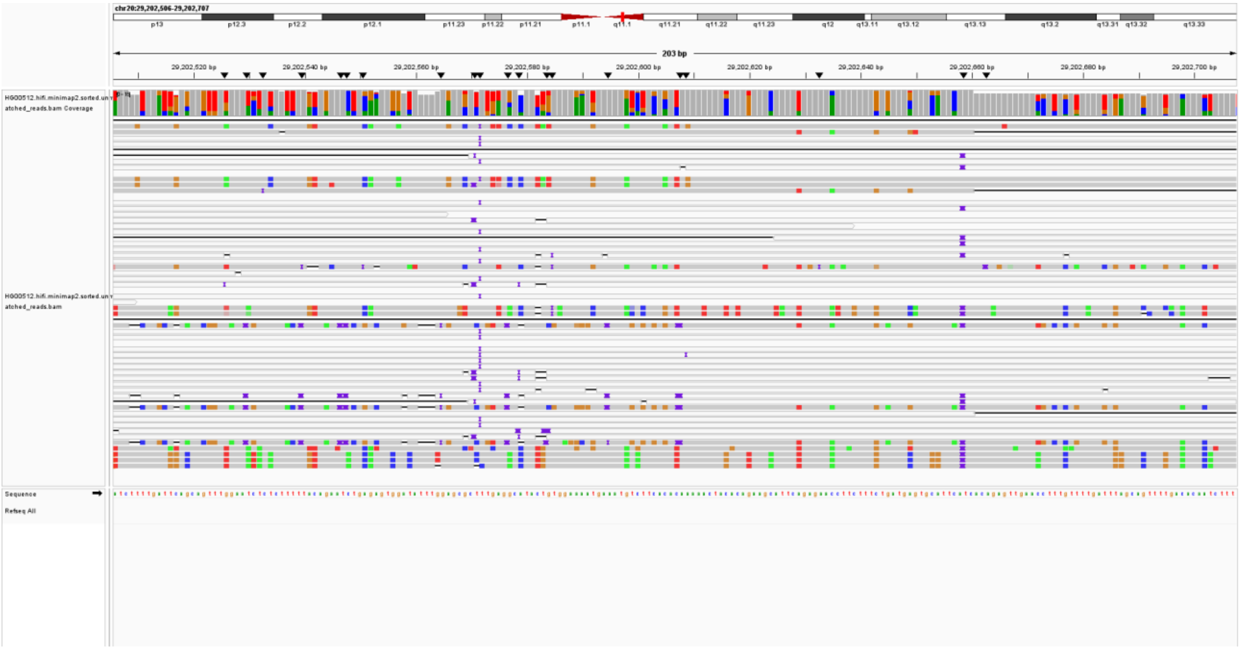

ALU-HG00512-chr16:33,029,593; (lack reads support)

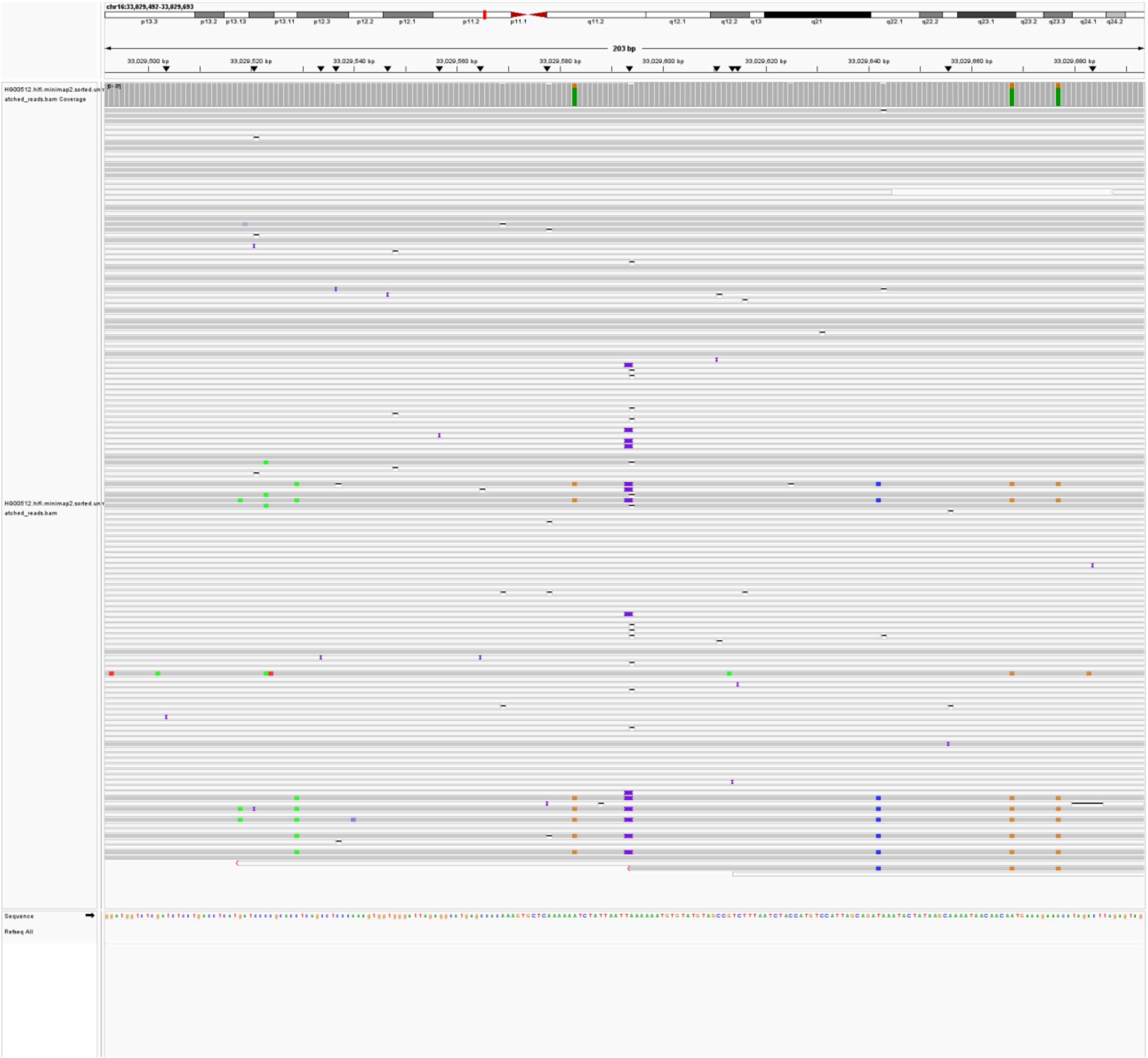

ALU-HG00512-chr16:90,145,128; (lack reads support)

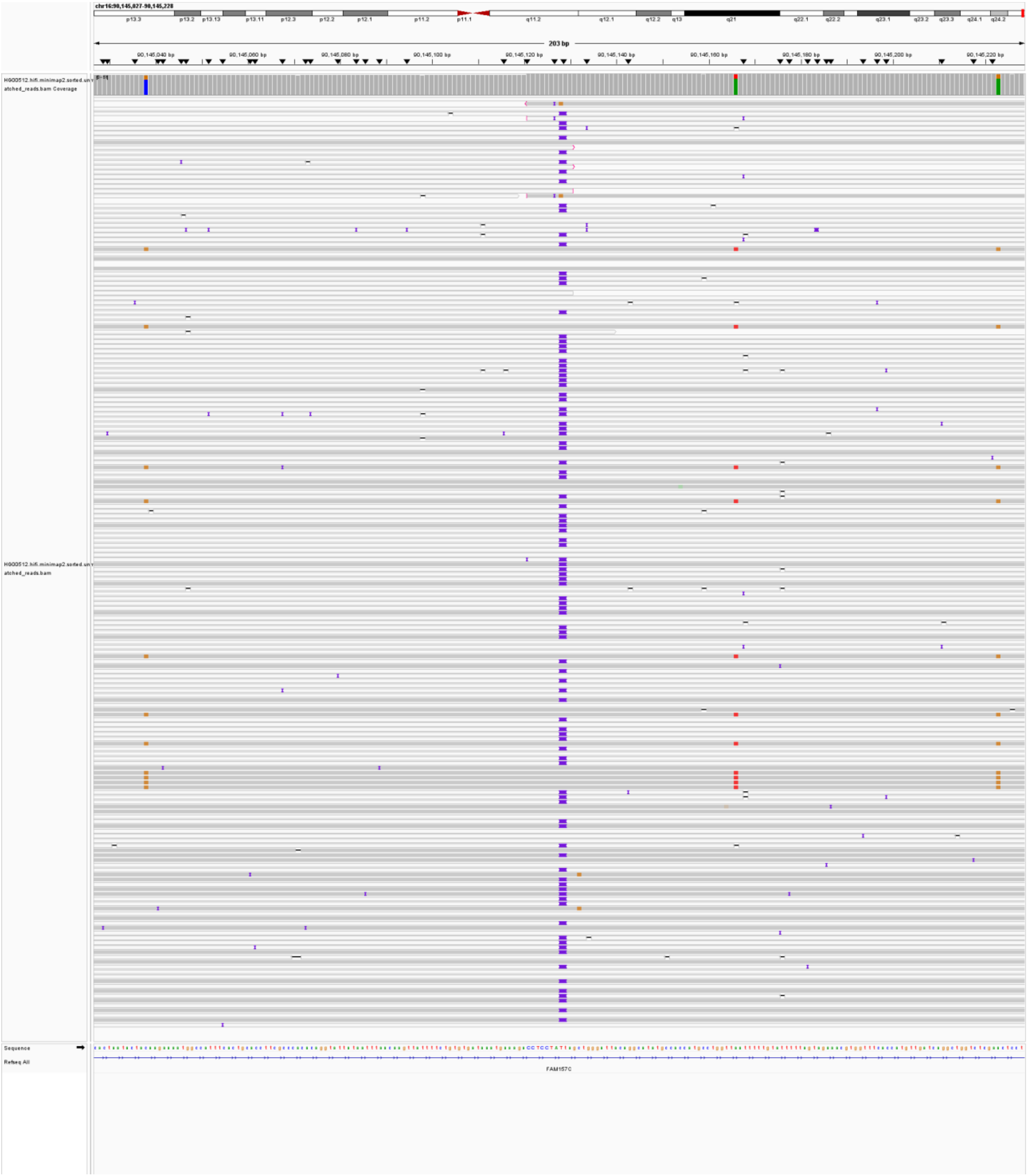

ALU-HG00512-chr17:36,272,223; (lack reads support)

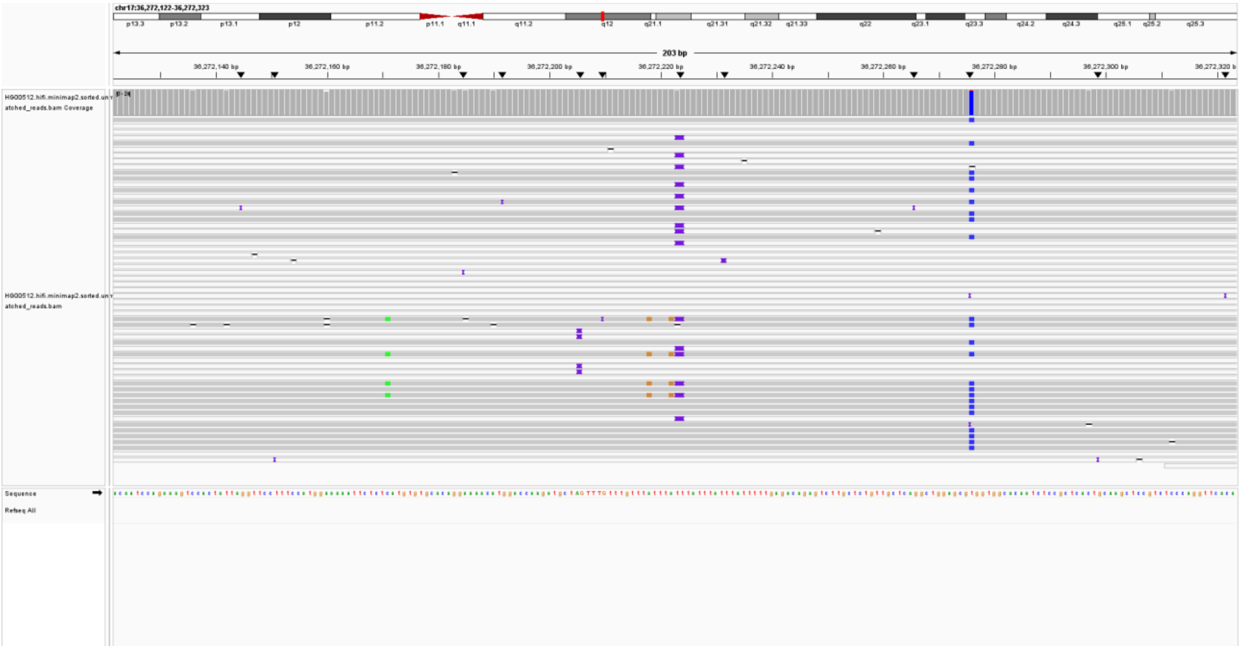

ALU-HG00512-chr17:81,352,658; (misclassification by MEIsensor)

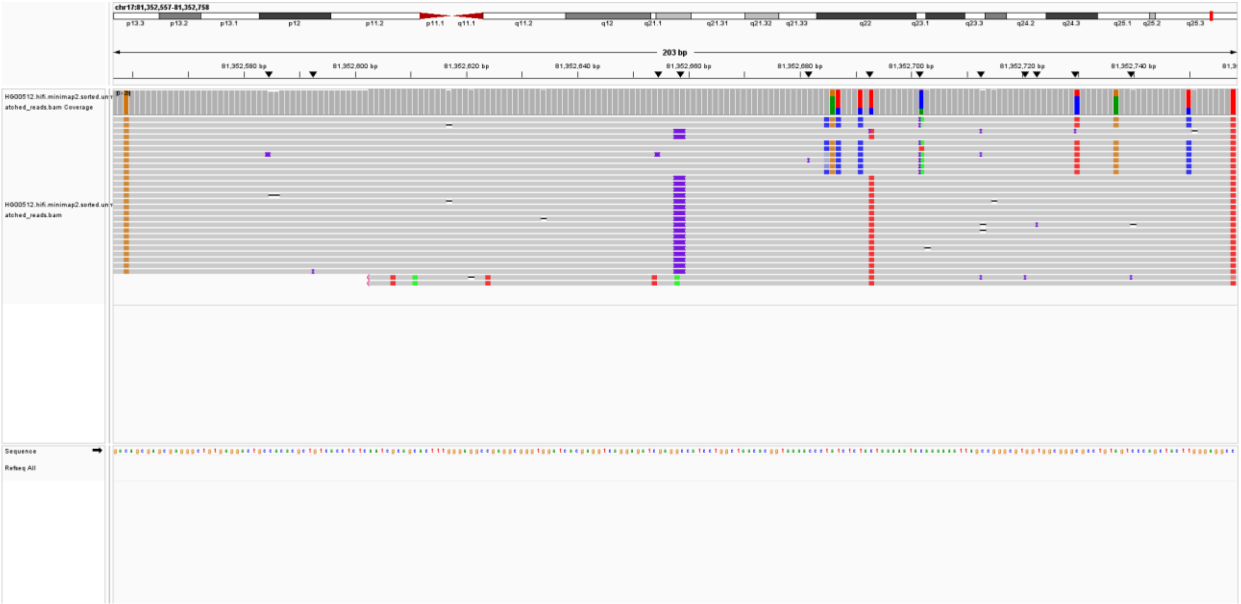

ALU-HG00512-chr19:54,753,605; (lack reads support)

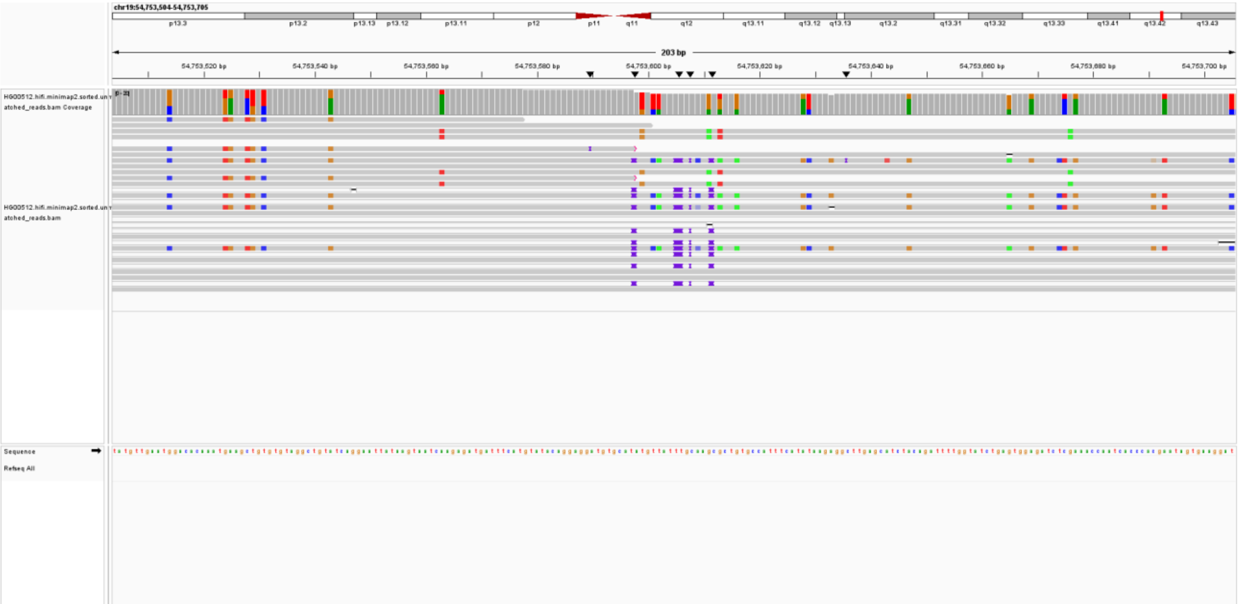


ALU-HG00512-chr1:231,796,350; (lack reads support)

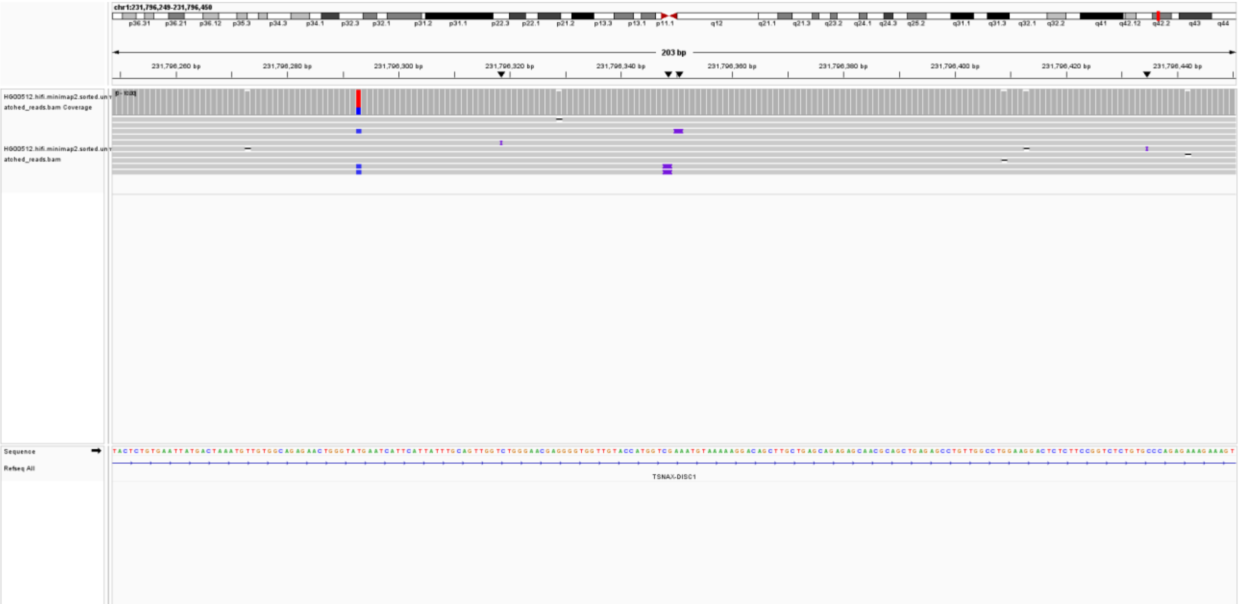

ALU-HG00512-chr20:29,258,308; (lack reads support)

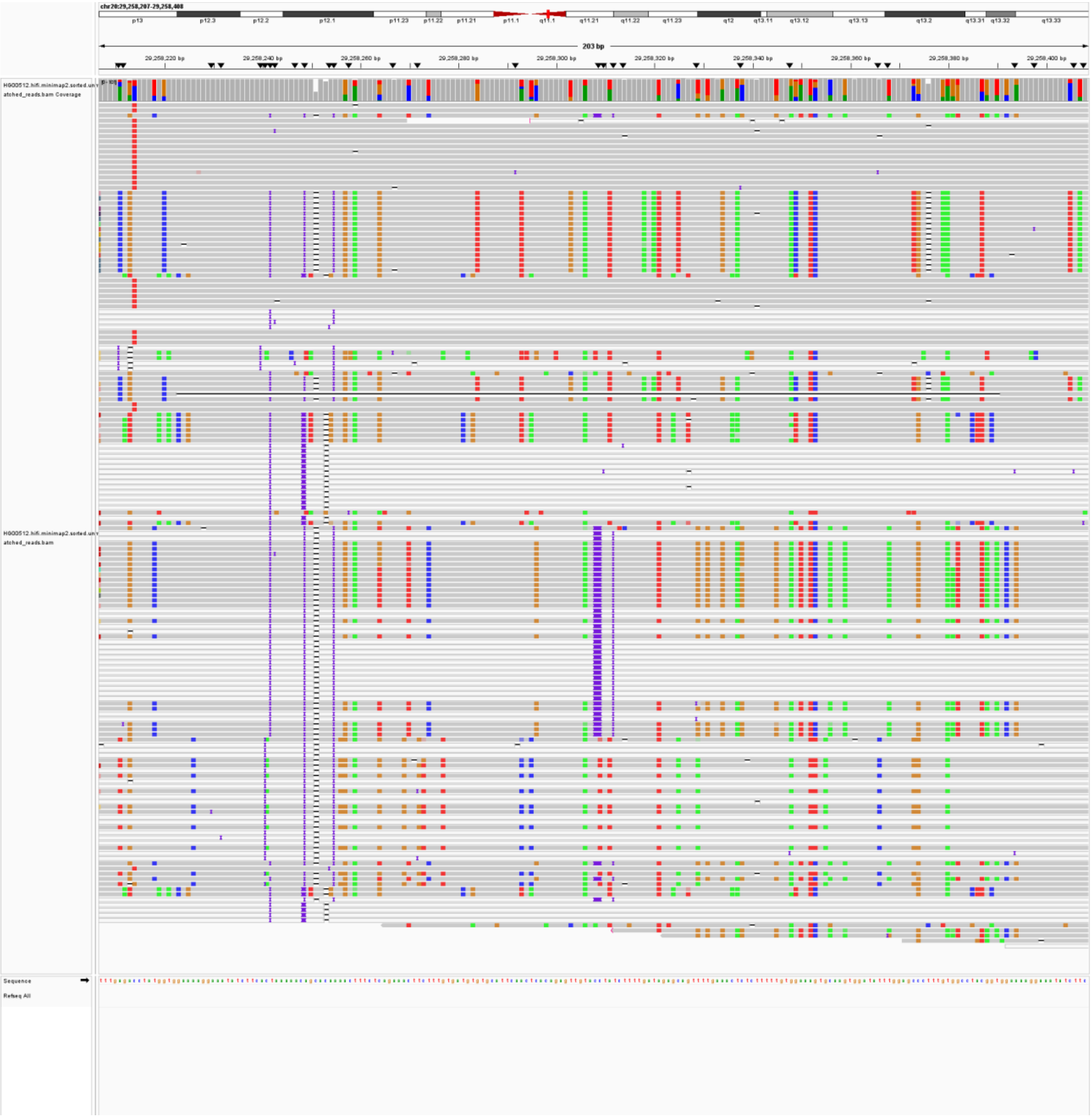

ALU-HG00512-chr20:29,264,446; (lack reads support)

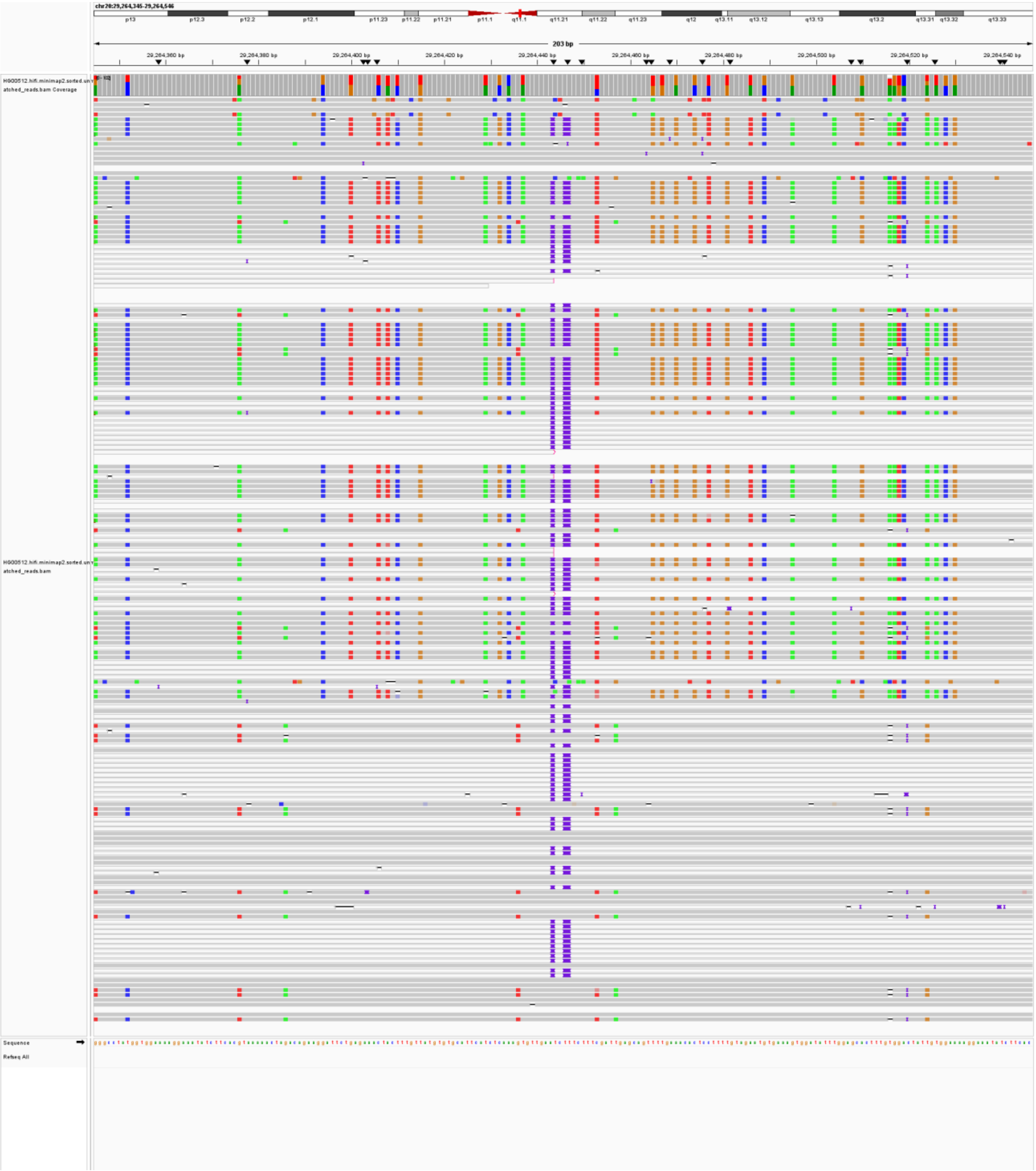

ALU-HG00512-chr20:30,215,832; (lack reads support)

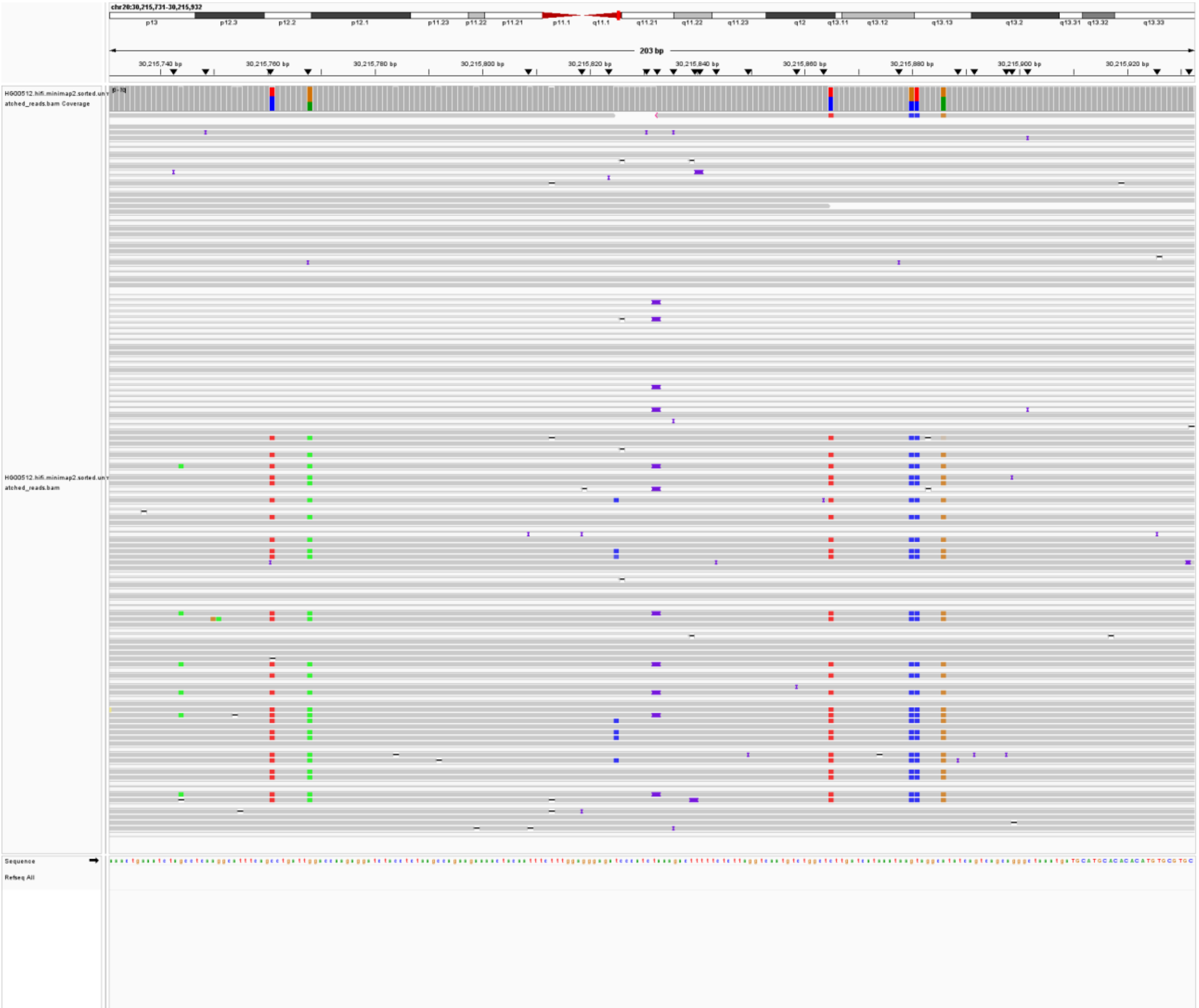

ALU-HG00512-chr21:23,927,857; (misclassification by MEIsensor)

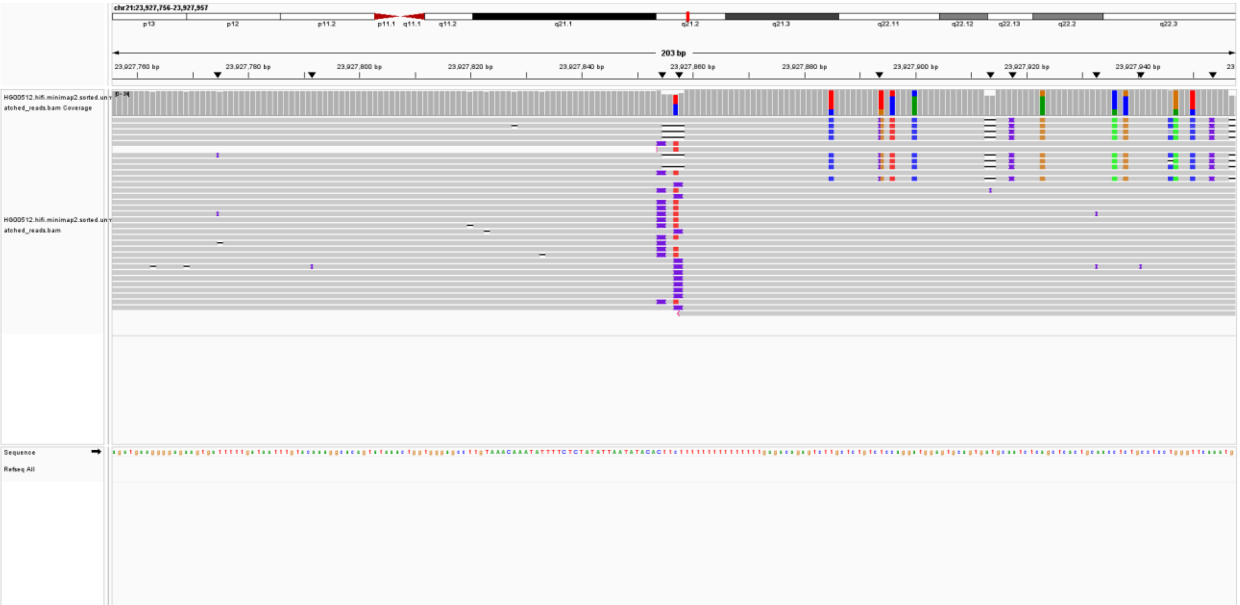

ALU-HG00512-chr21:32,428,506; (lack reads support)

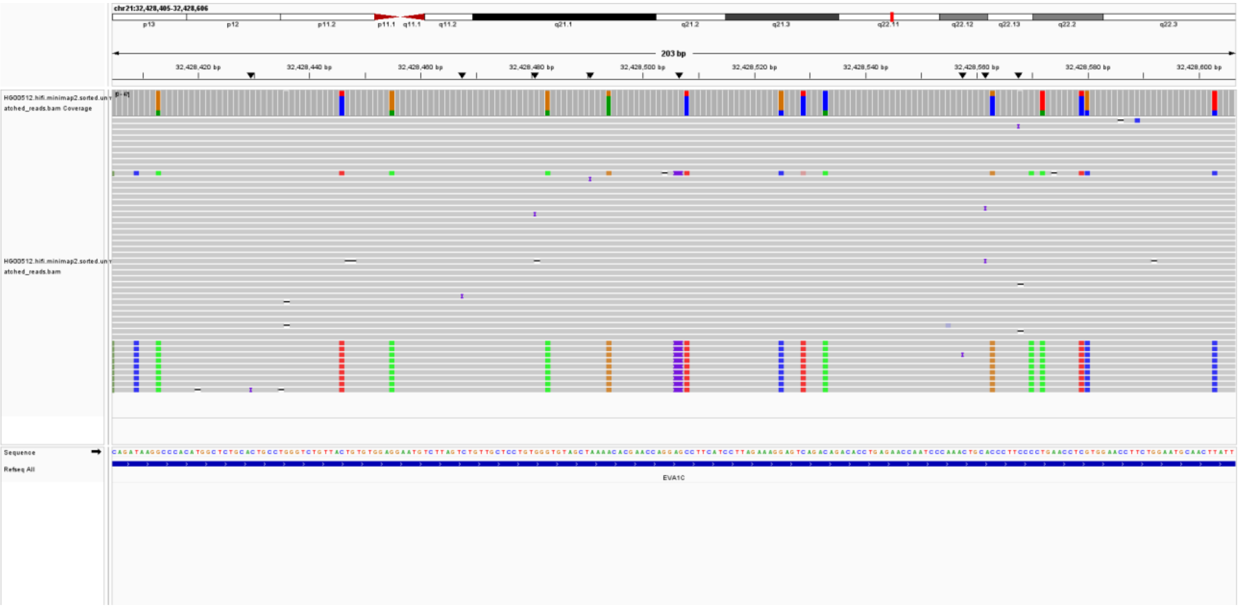

ALU-HG00512-chr21:6,696,470; (lack reads support)

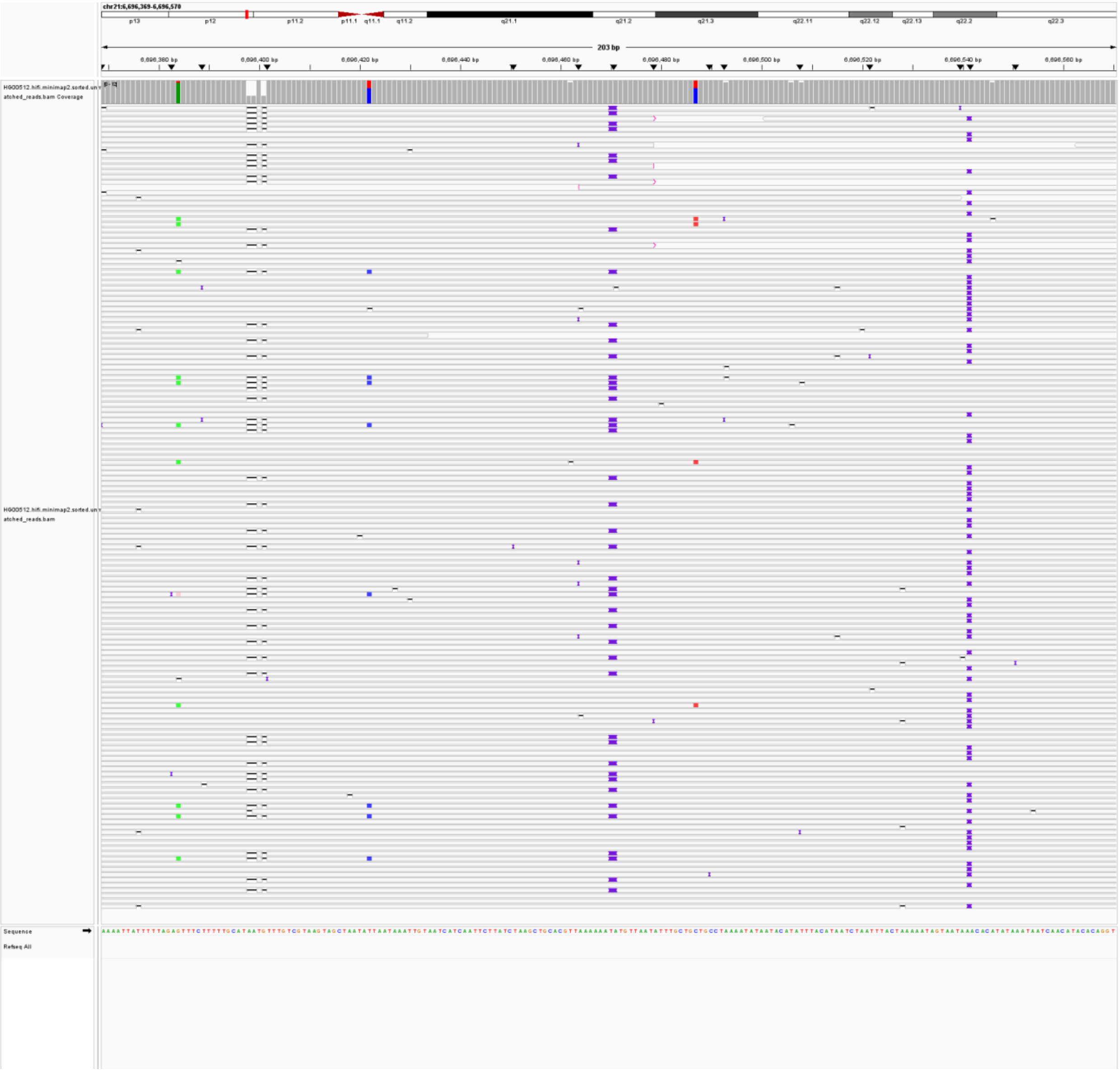

ALU-HG00512-chr21:7,023,540; (lack reads support)

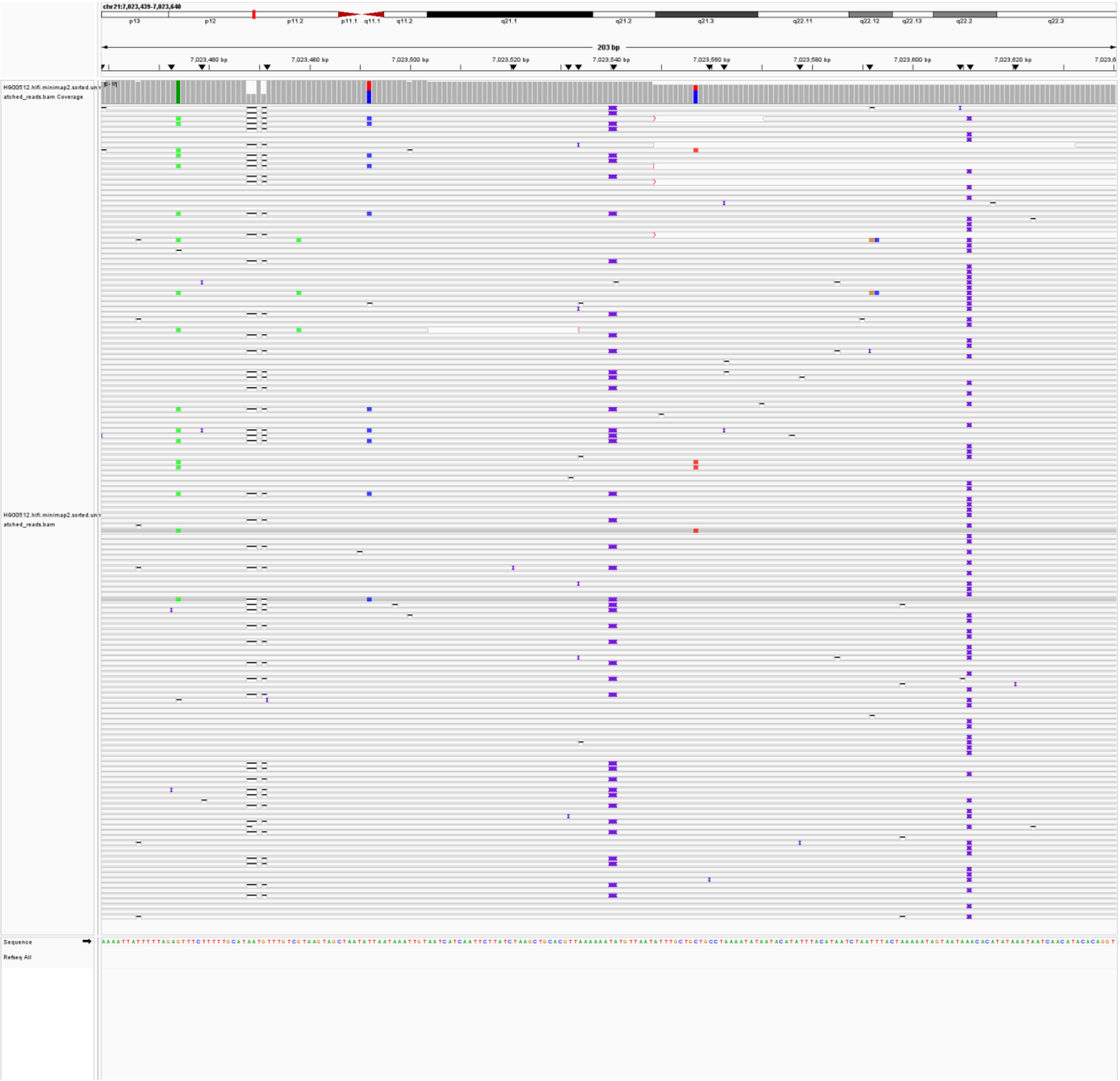

ALU-HG00512-chr21:7,244,007; (lack reads support)

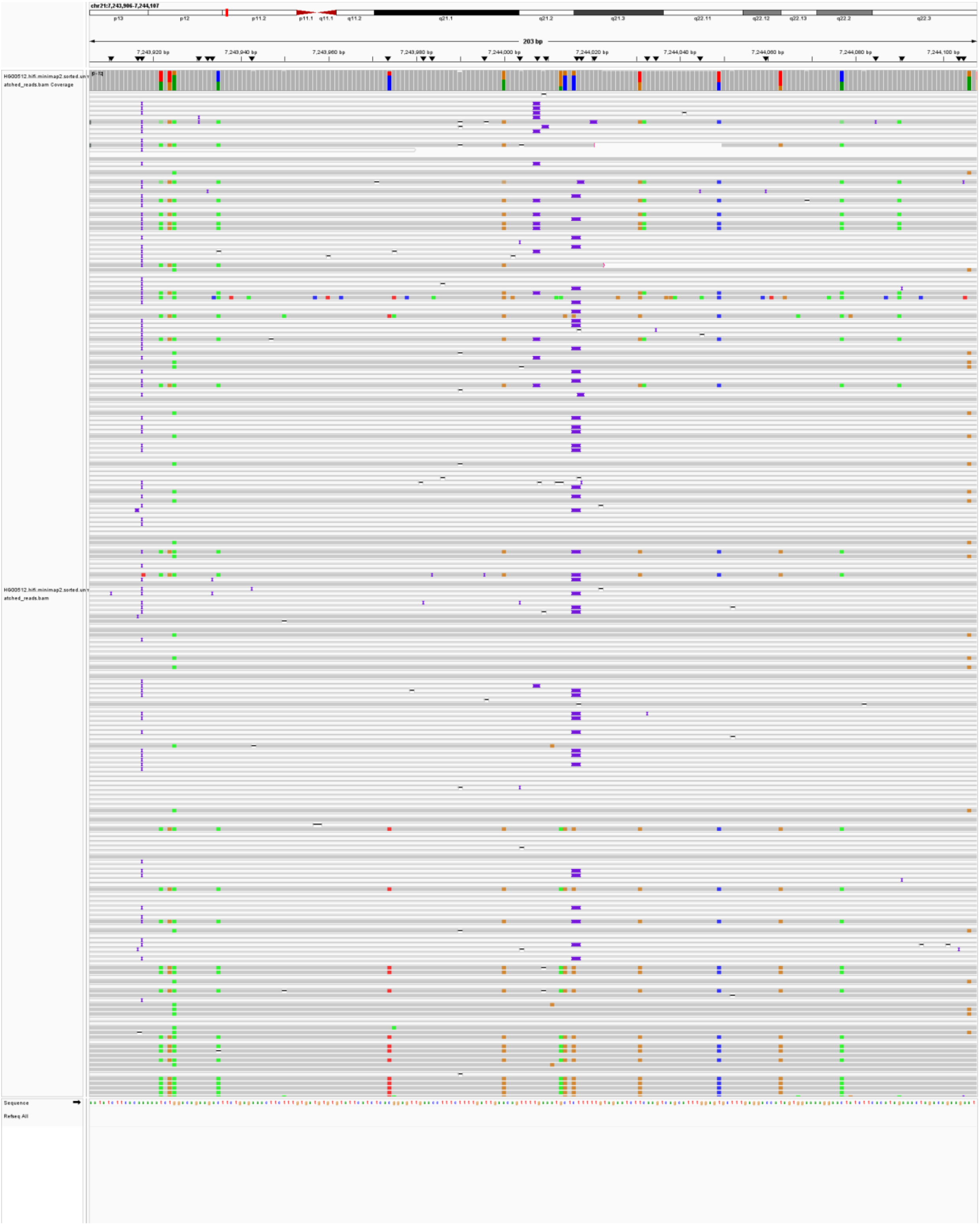

ALU-HG00512-chr21:7,405,318; (lack reads support)

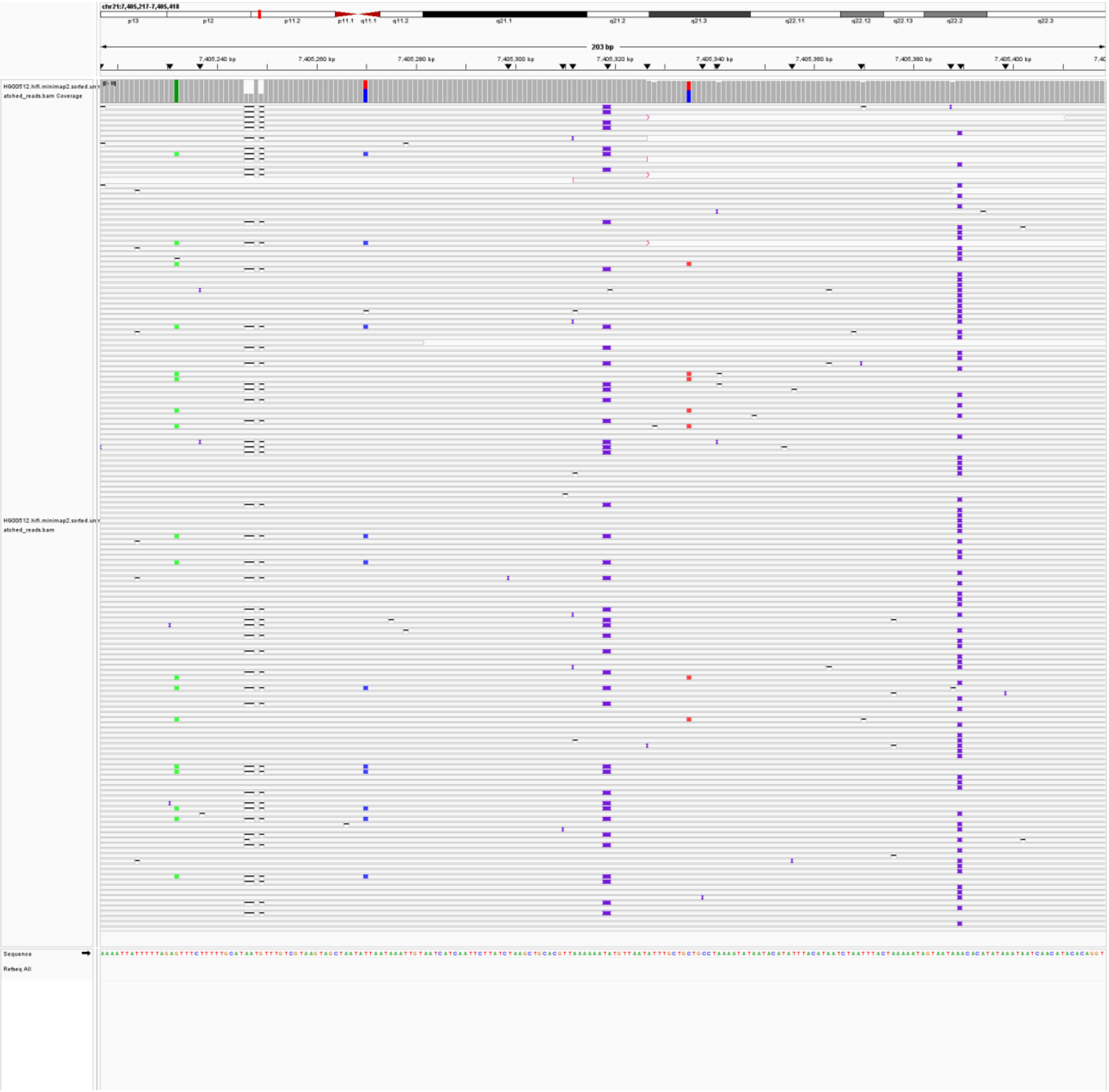

ALU-HG00512-chr21:9,328,301; (lack reads support)

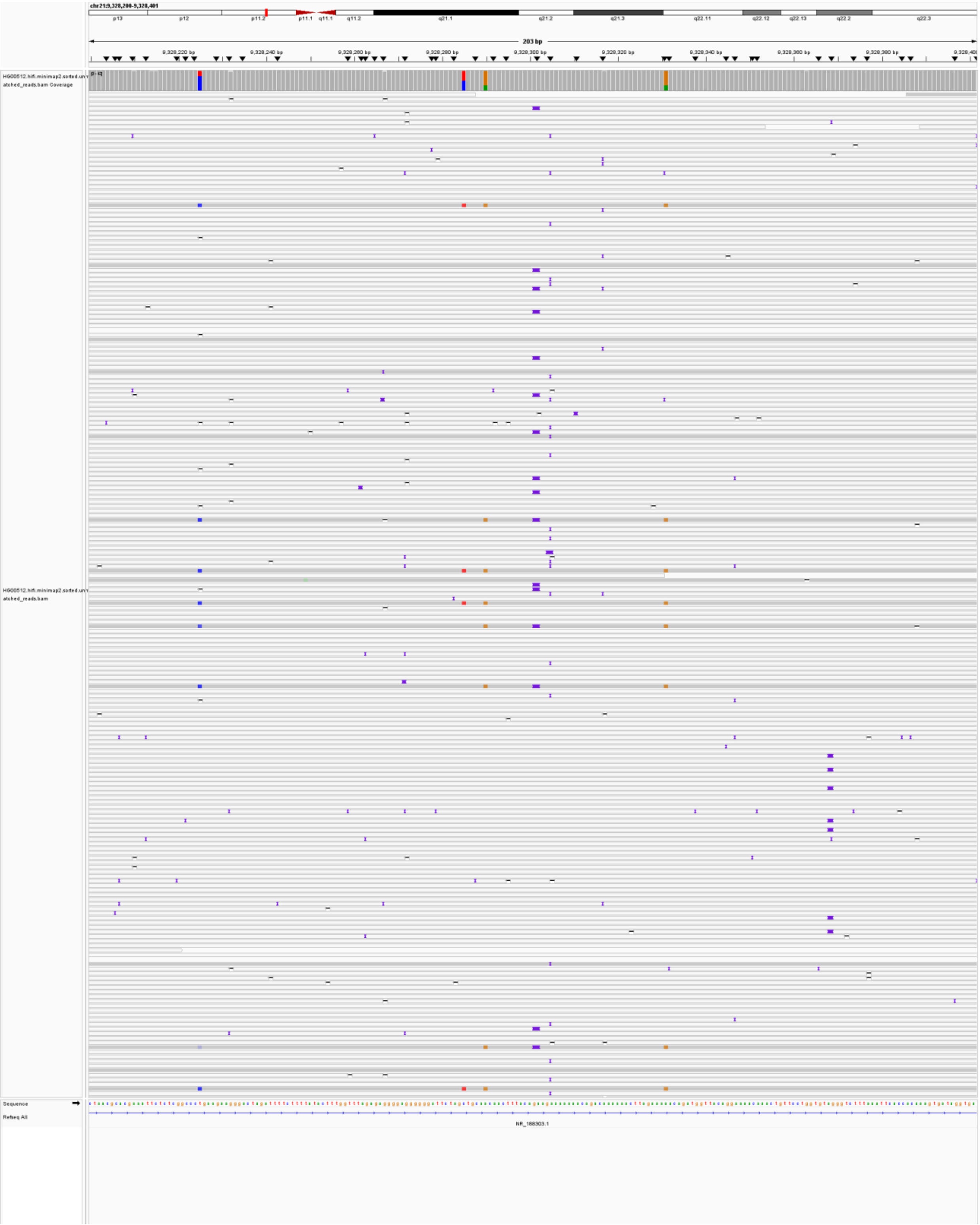

ALU-HG00512-chr22:12,423,578; (lack reads support)

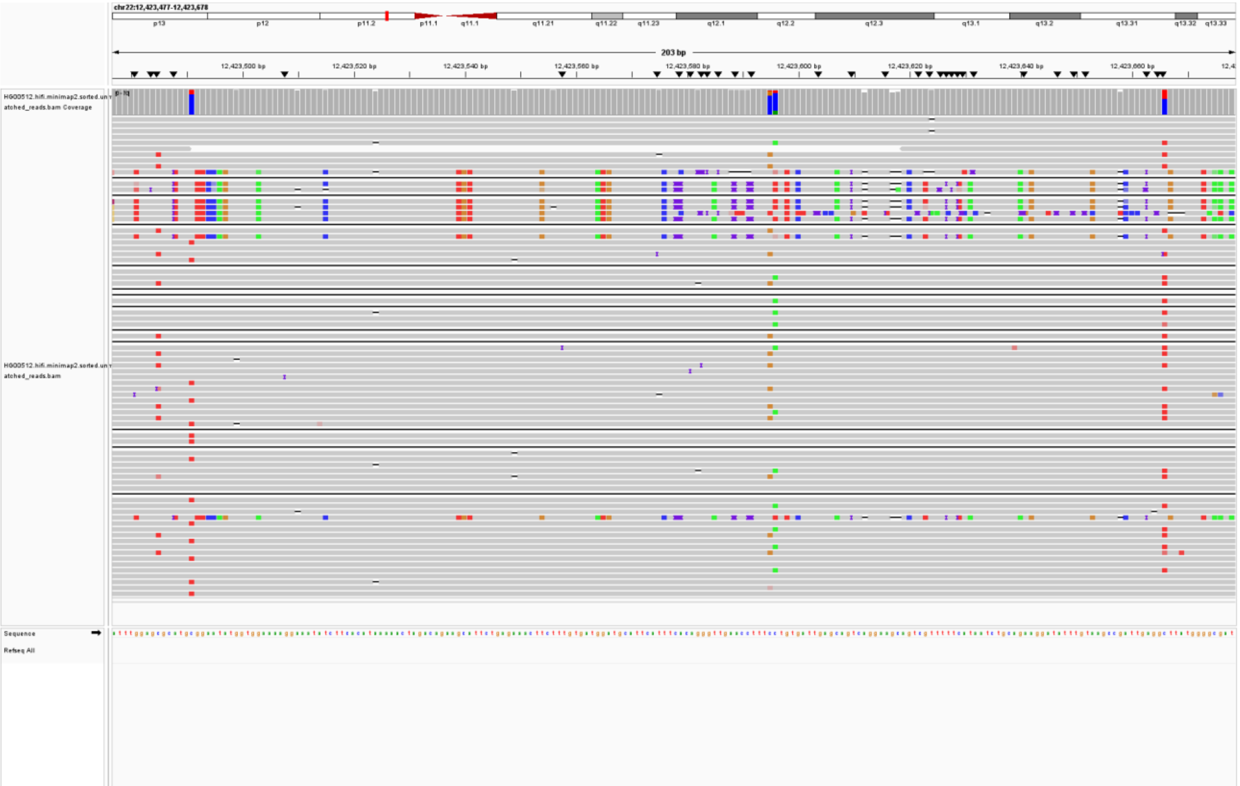

ALU-HG00512-chr22:15,698,177; (lack reads support)

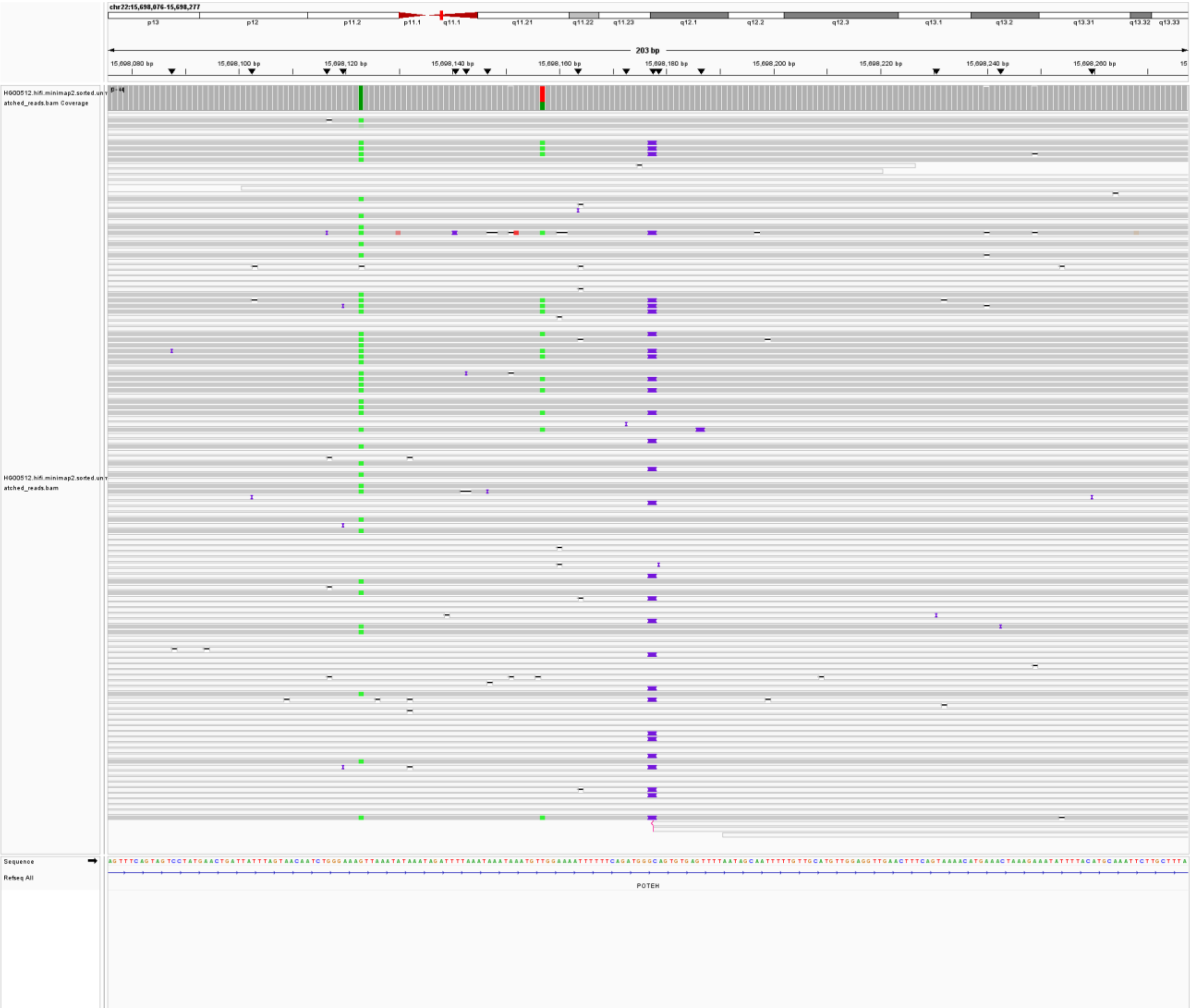

ALU-HG00512-chr22:49,879,744; (lack reads support)

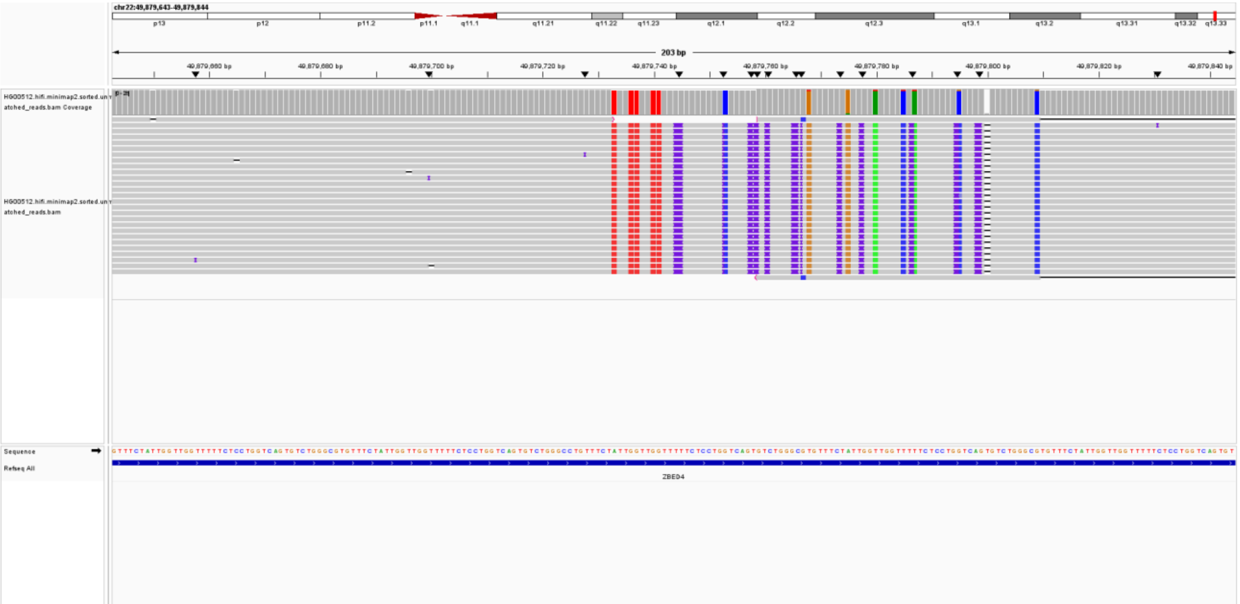

ALU-HG00512-chr2:13,374,169; (lack reads support)

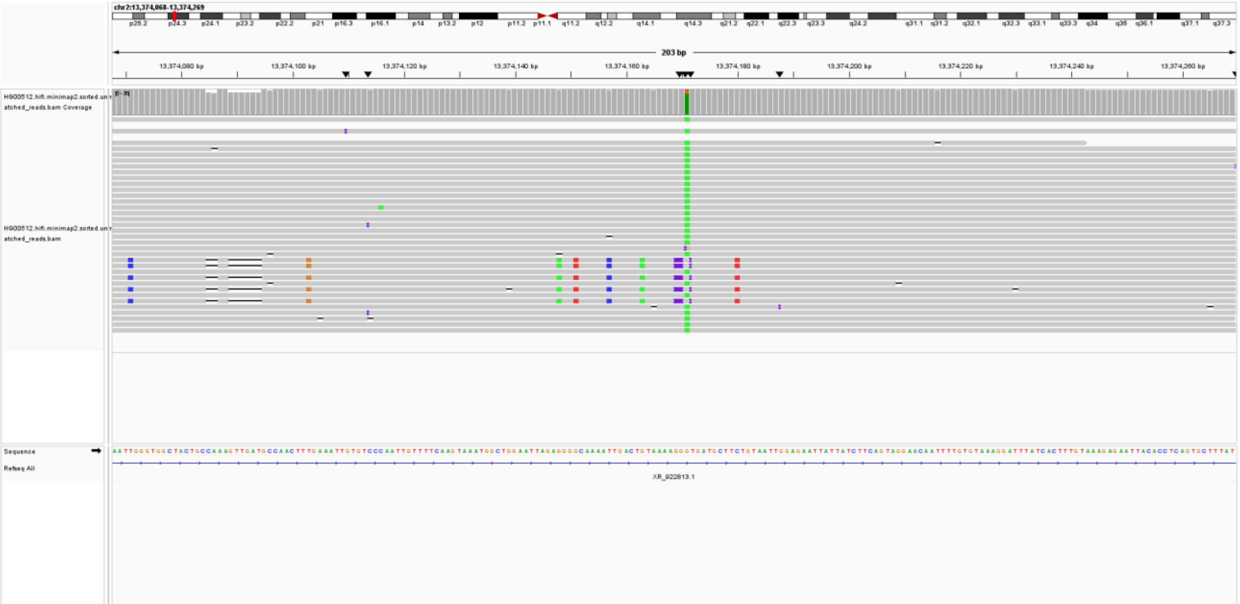

ALU-HG00512-chr2:170,128,752; (lack reads support)

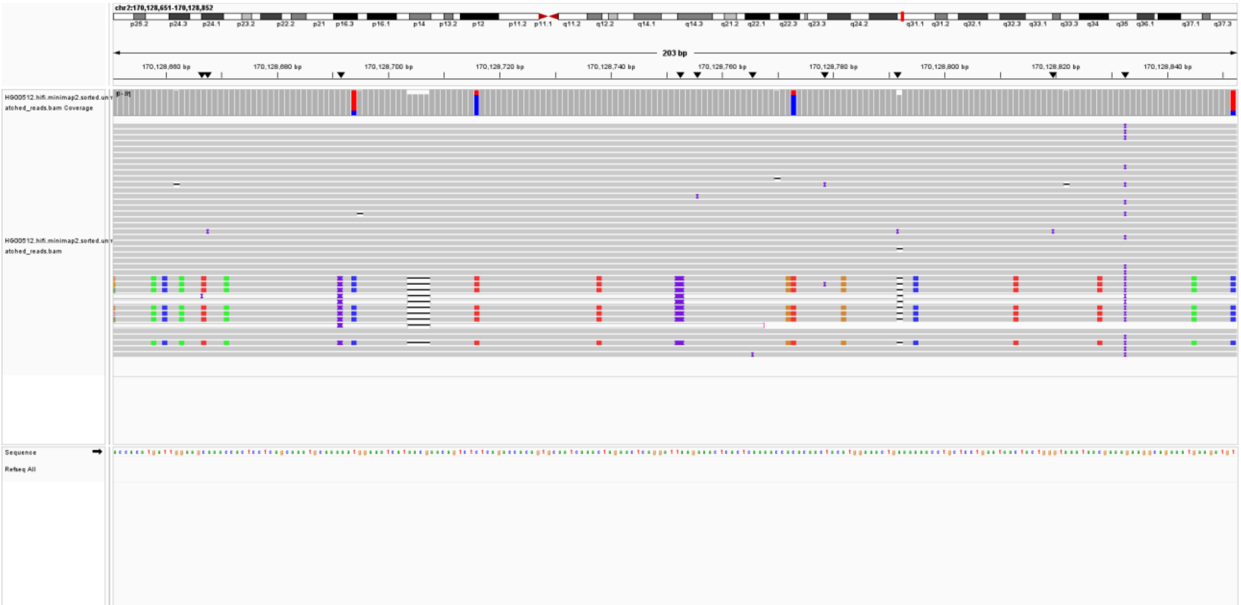

ALU-HG00512-chr3:39,174,528; (misclassification by MEIsensor)

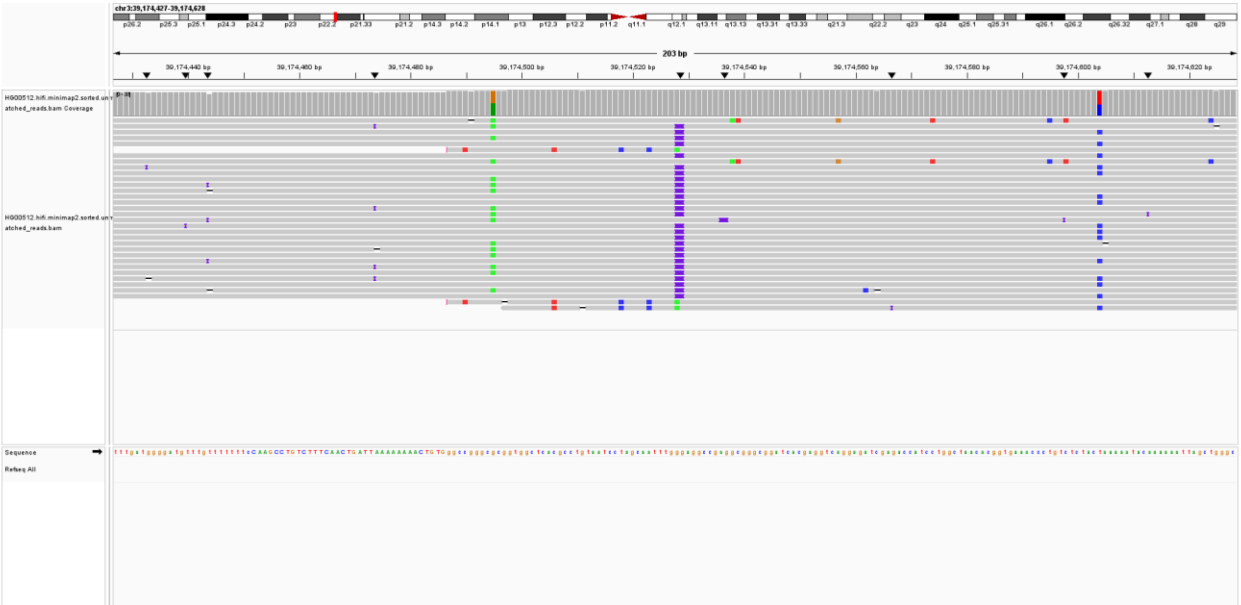

ALU-HG00512-chr3:58,436,542; (lack reads support)

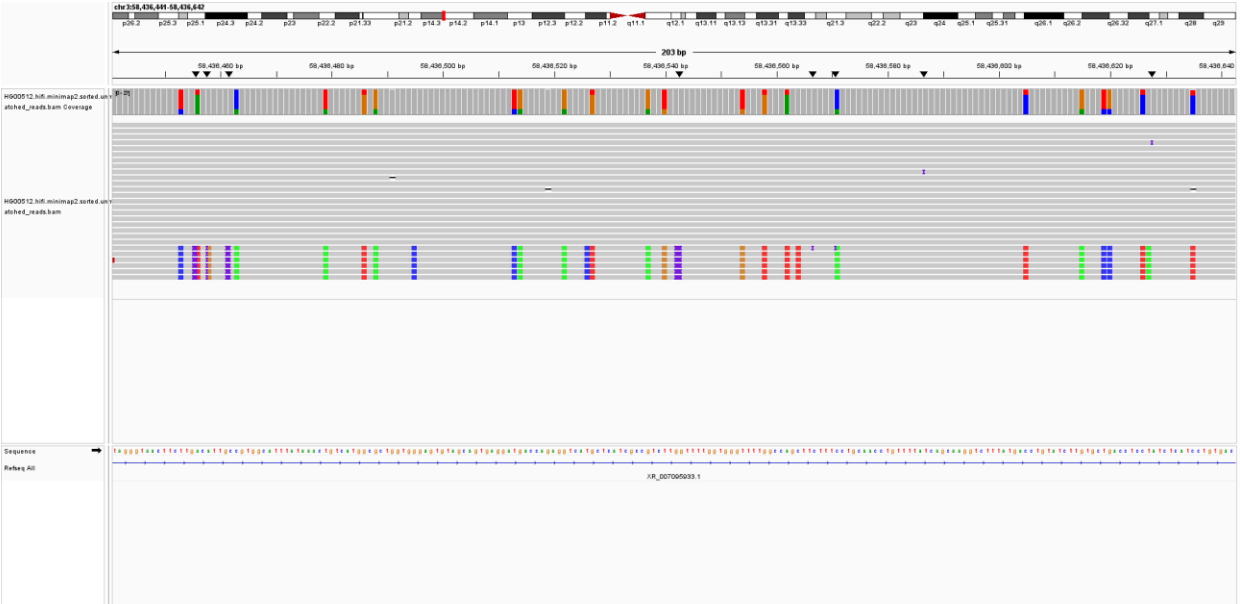

ALU-HG00512-chr4:158,660,816; (misclassification by MEIsensor)

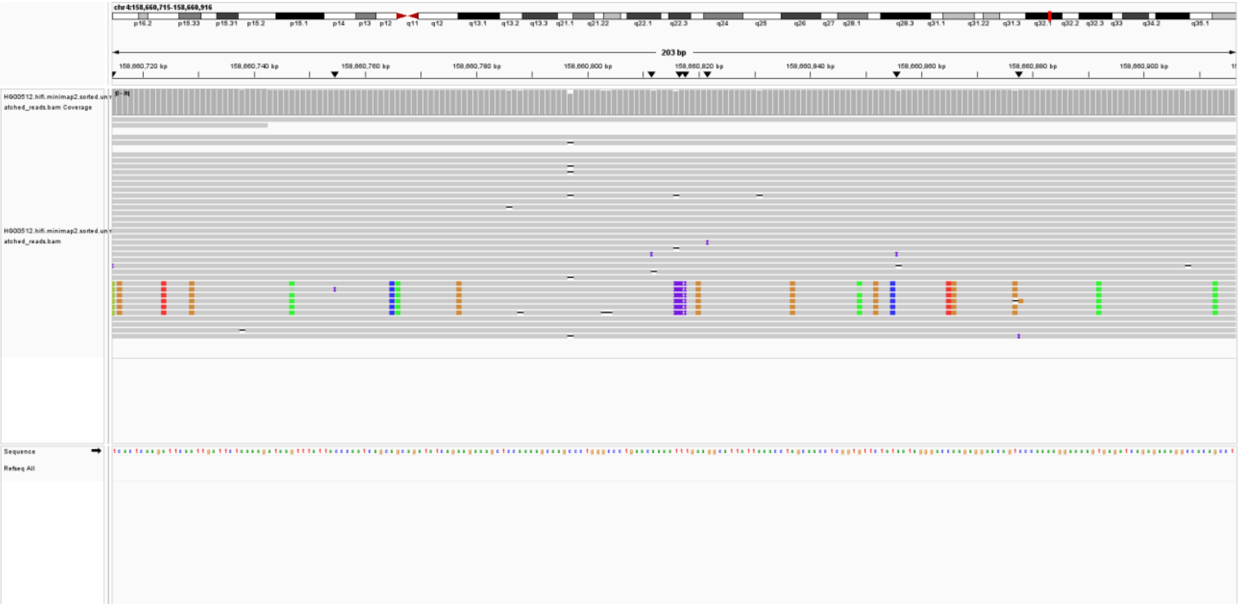

ALU-HG00512-chr6:104,042,792; (misclassification by MEIsensor)

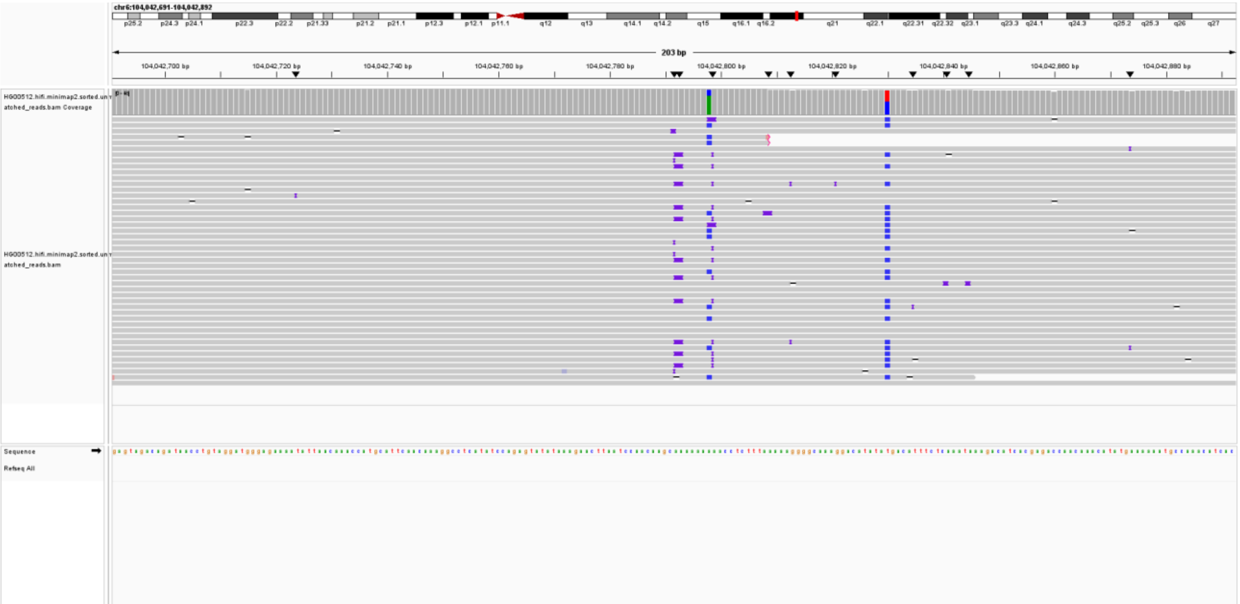

ALU-HG00512-chr6:31,041,860; (lack reads support)

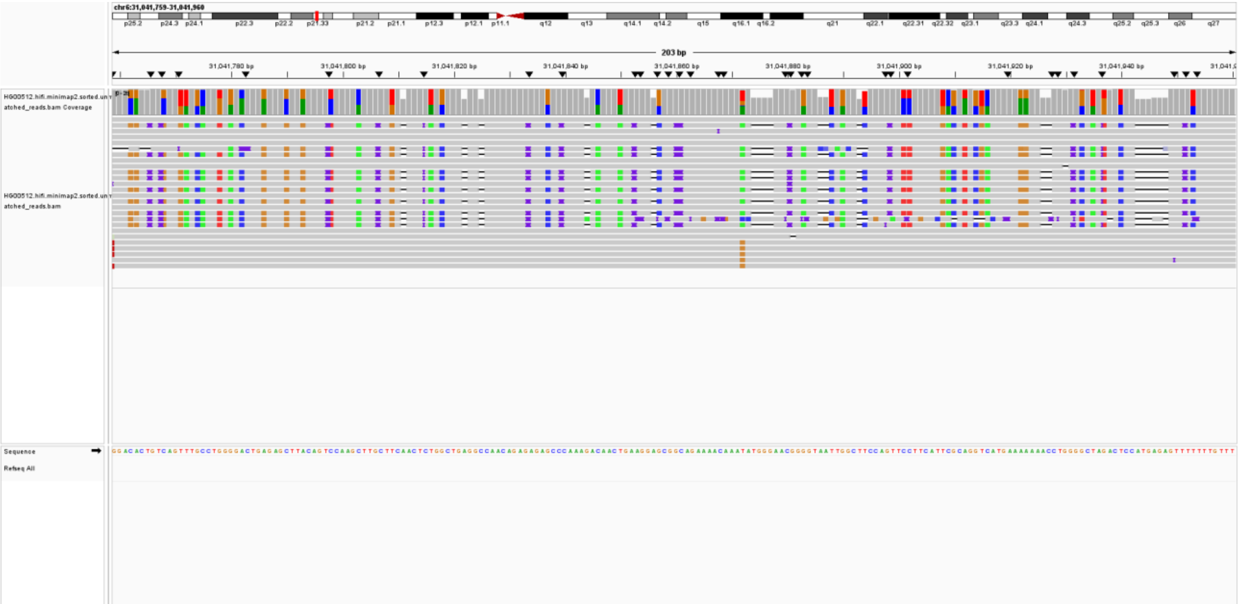

ALU-HG00512-chr7:102,549,903; (lack reads support)

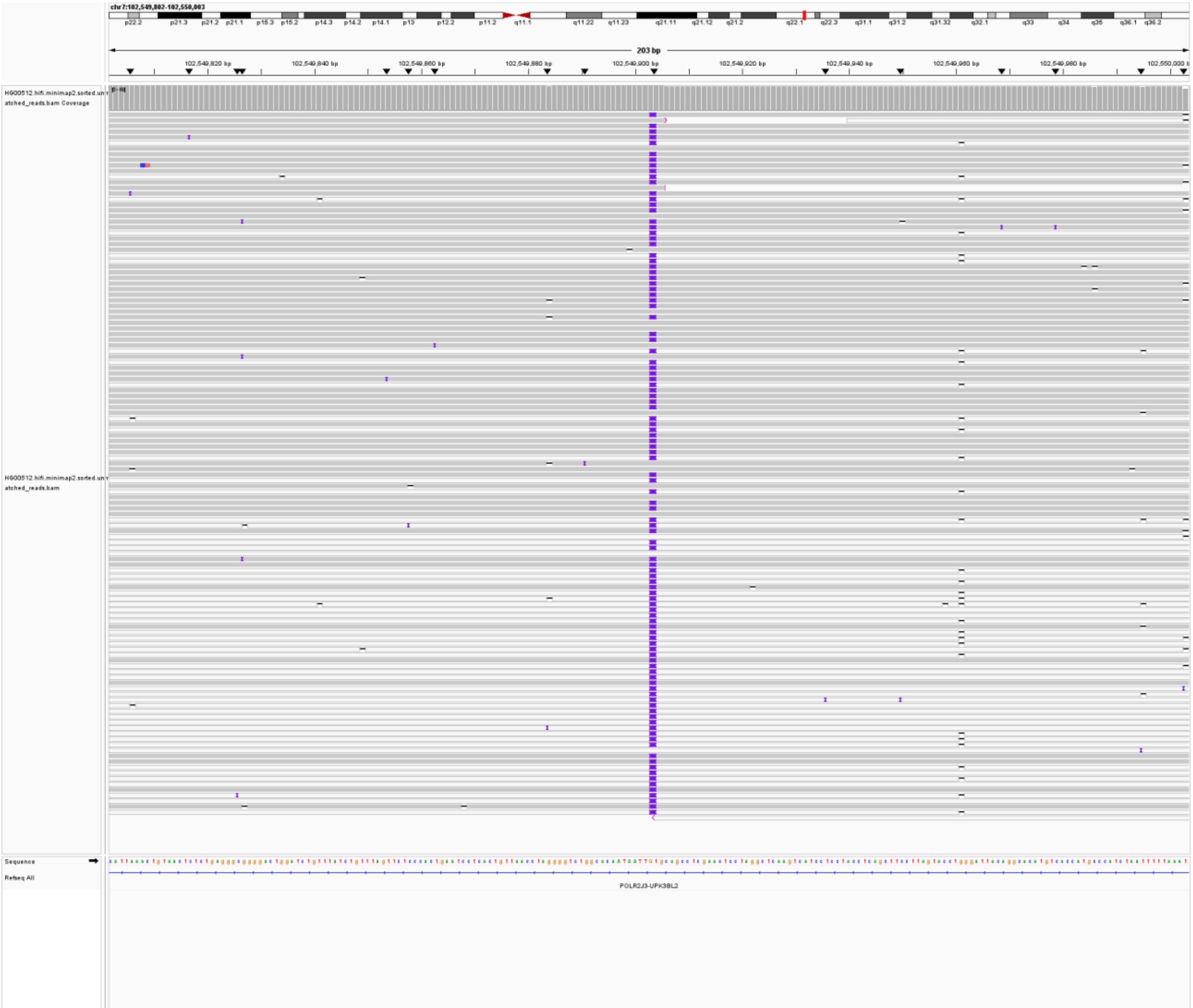

ALU-HG00512-chr7:102,648,997; (lack reads support)

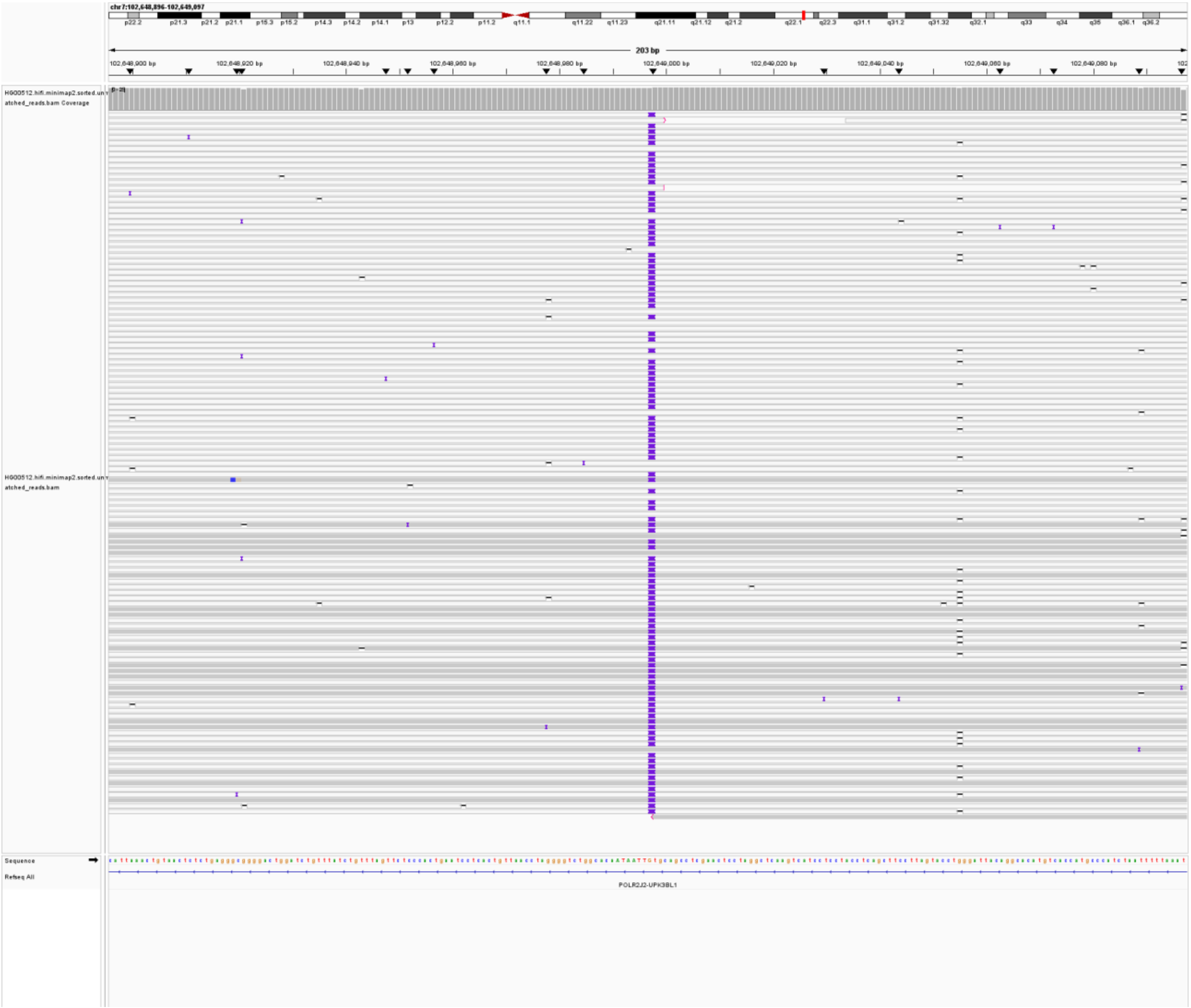

ALU-HG00512-chr7:142,452,258; (lack reads support)

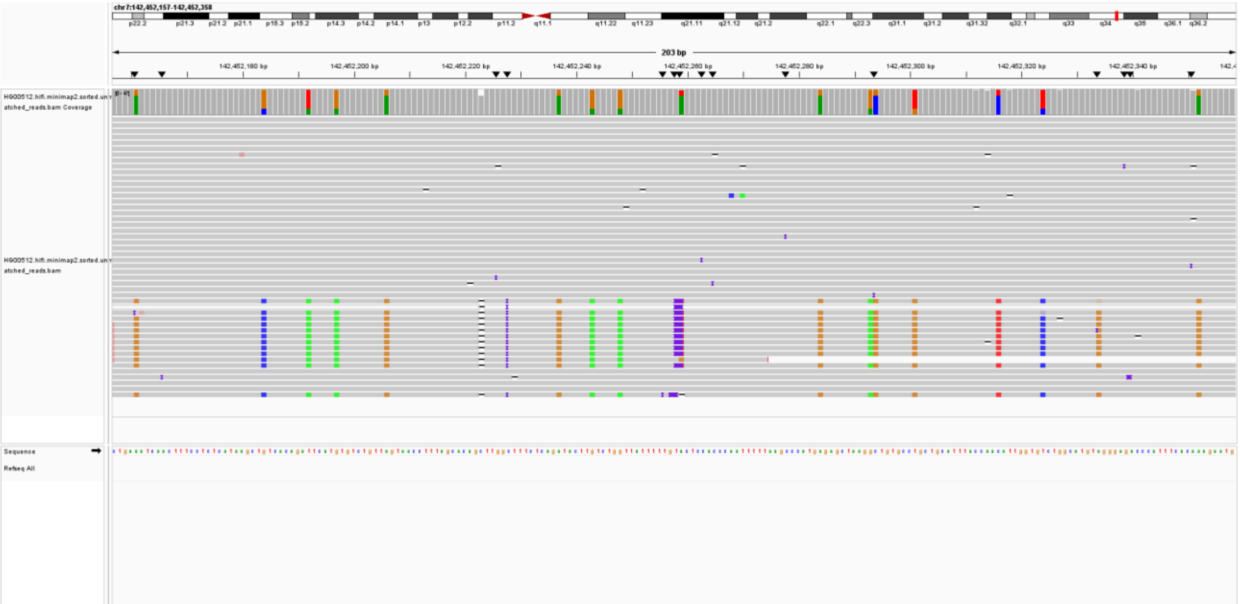

ALU-HG00512-chr7:142,458,367; (lack reads support)

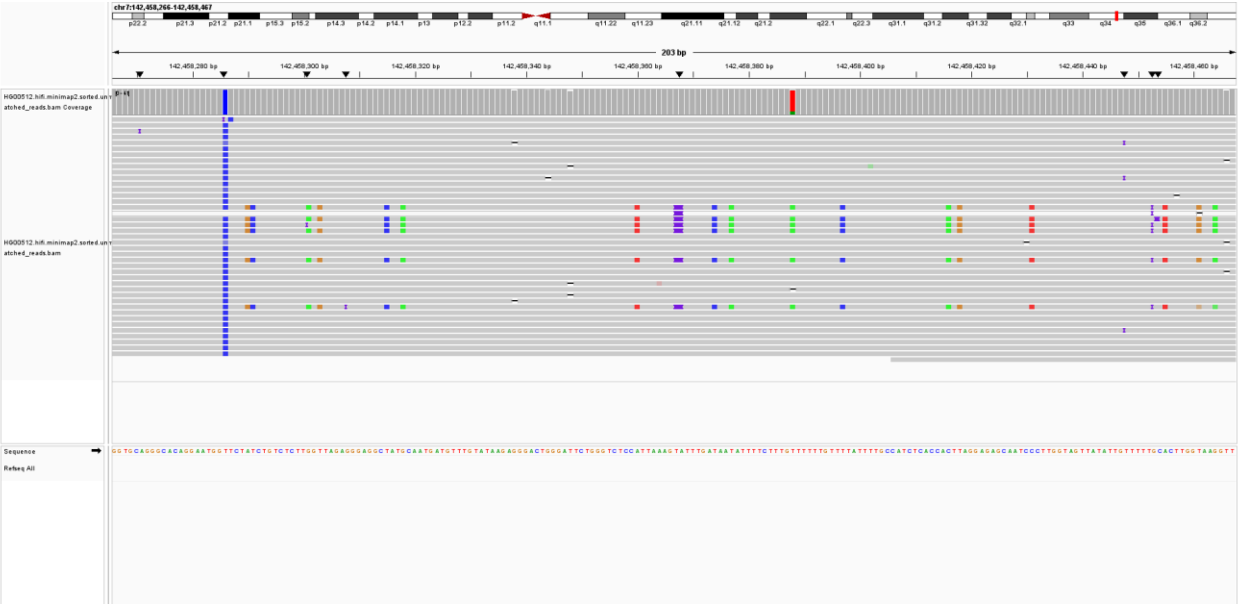

ALU-HG00512-chr7:142,477,316; (lack reads support)

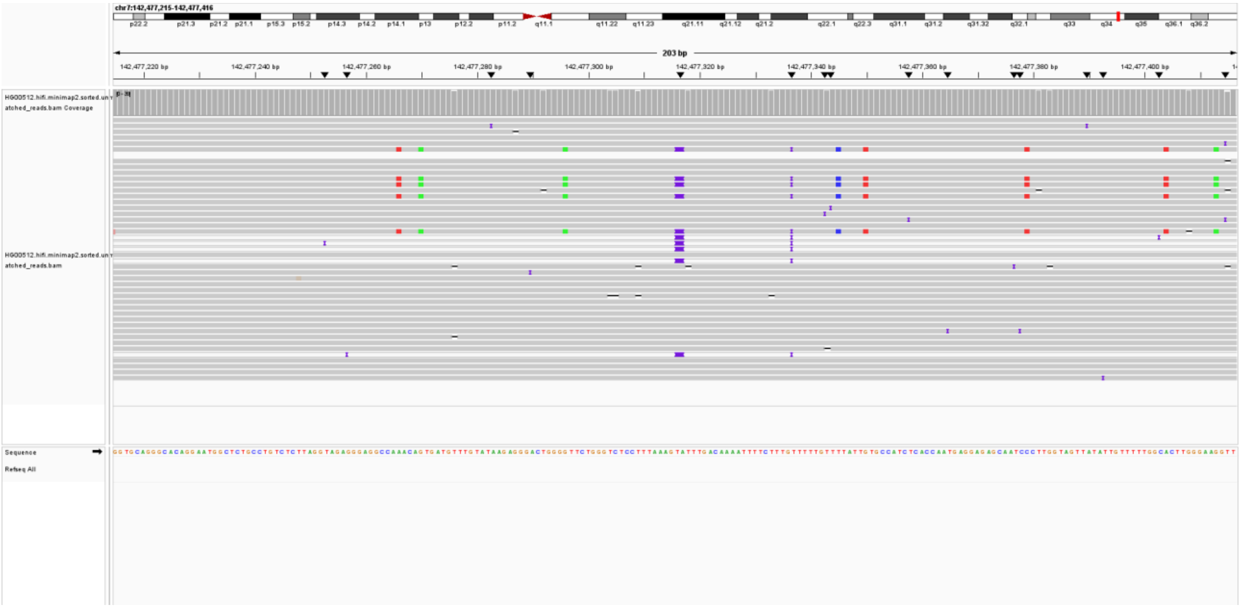

ALU-HG00512-chr7:58,183,285; (lack reads support)

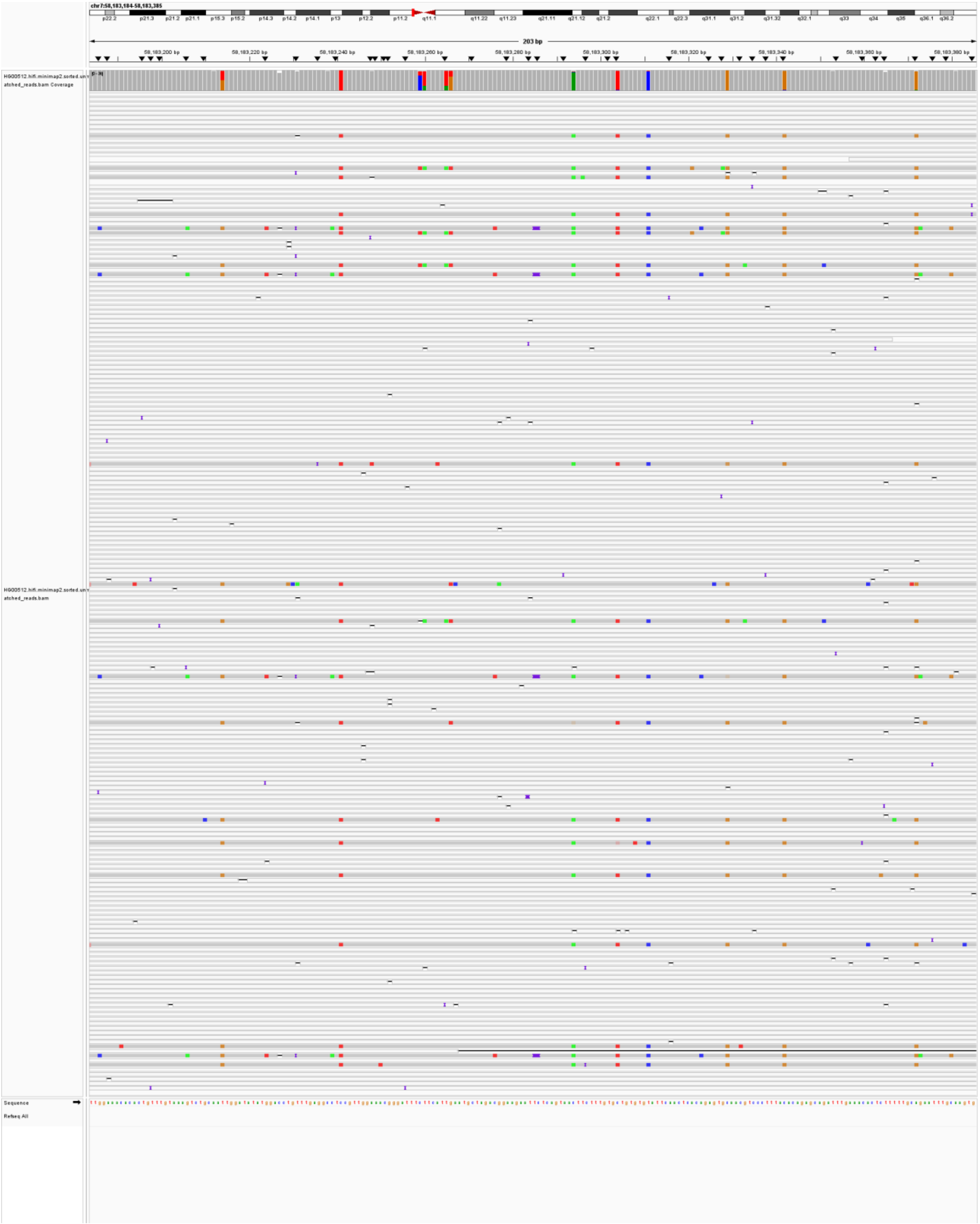

ALU-HG00512-chr7:58,197,577; (lack reads support)

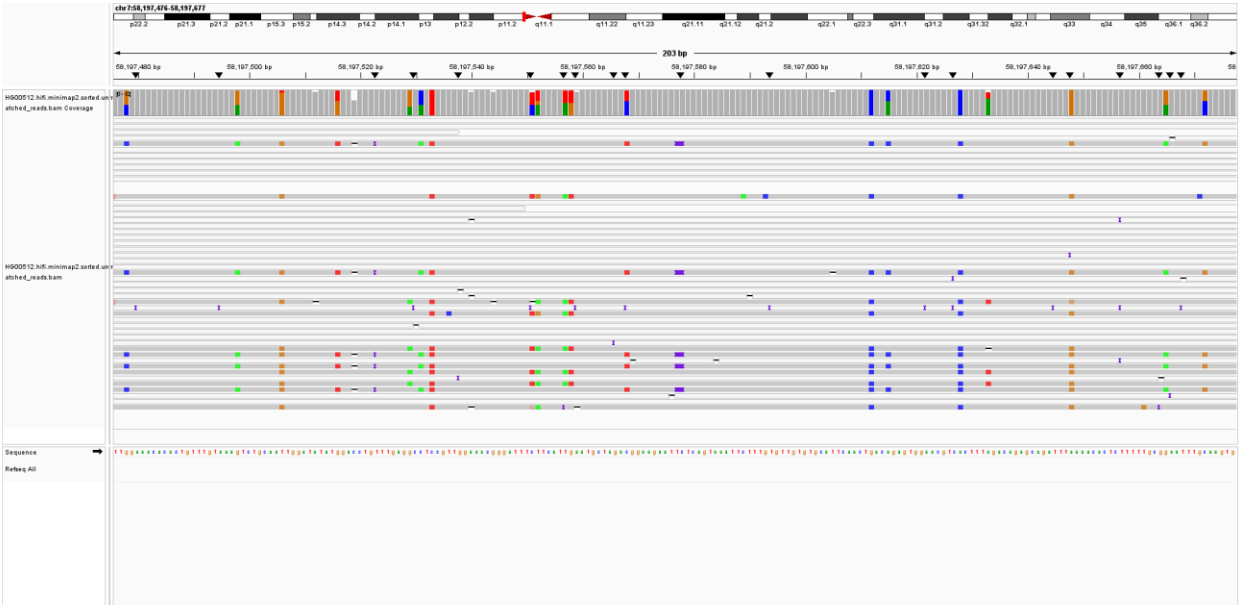

ALU-HG00512-chr7:59,140,991; (lack reads support)

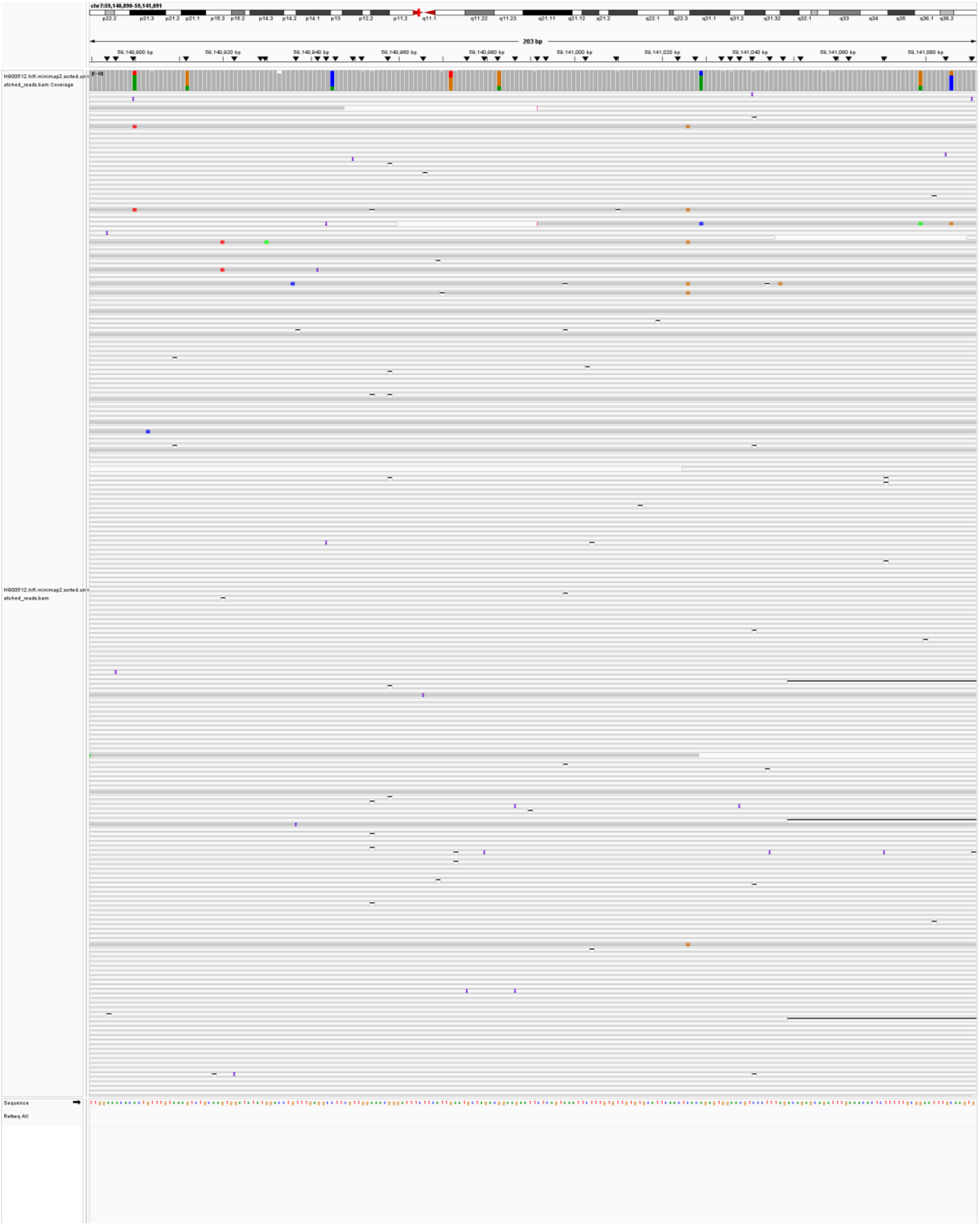

ALU-HG00512-chr9:129,426,080; (lack reads support)

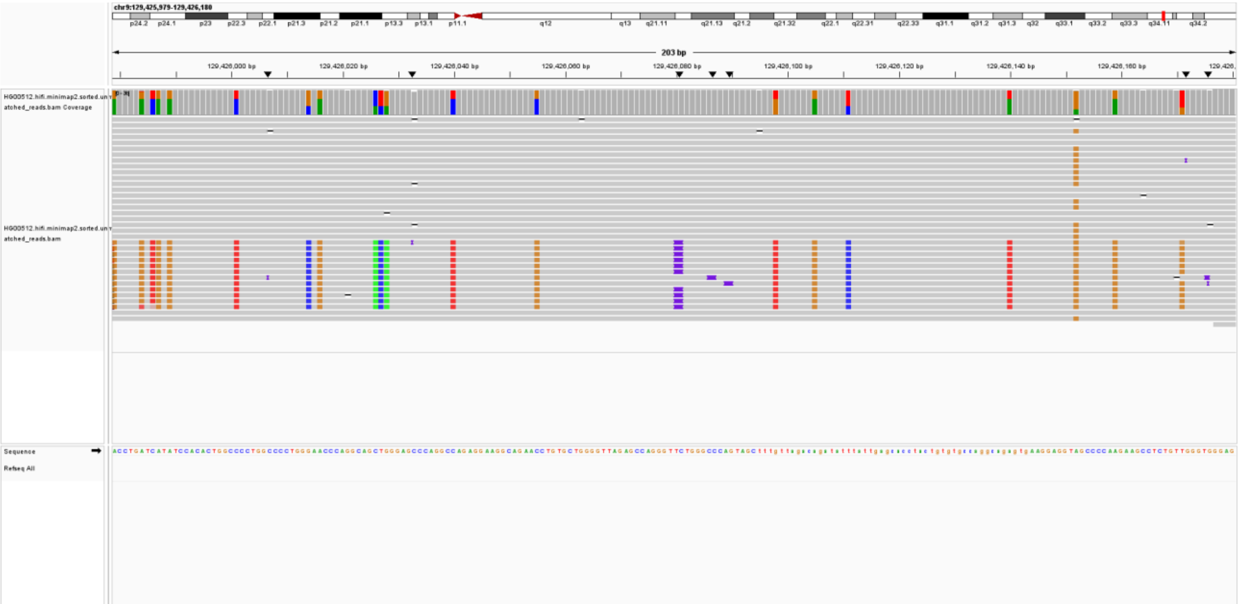

ALU-HG00512-chr9:41,975,916; (lack reads support)

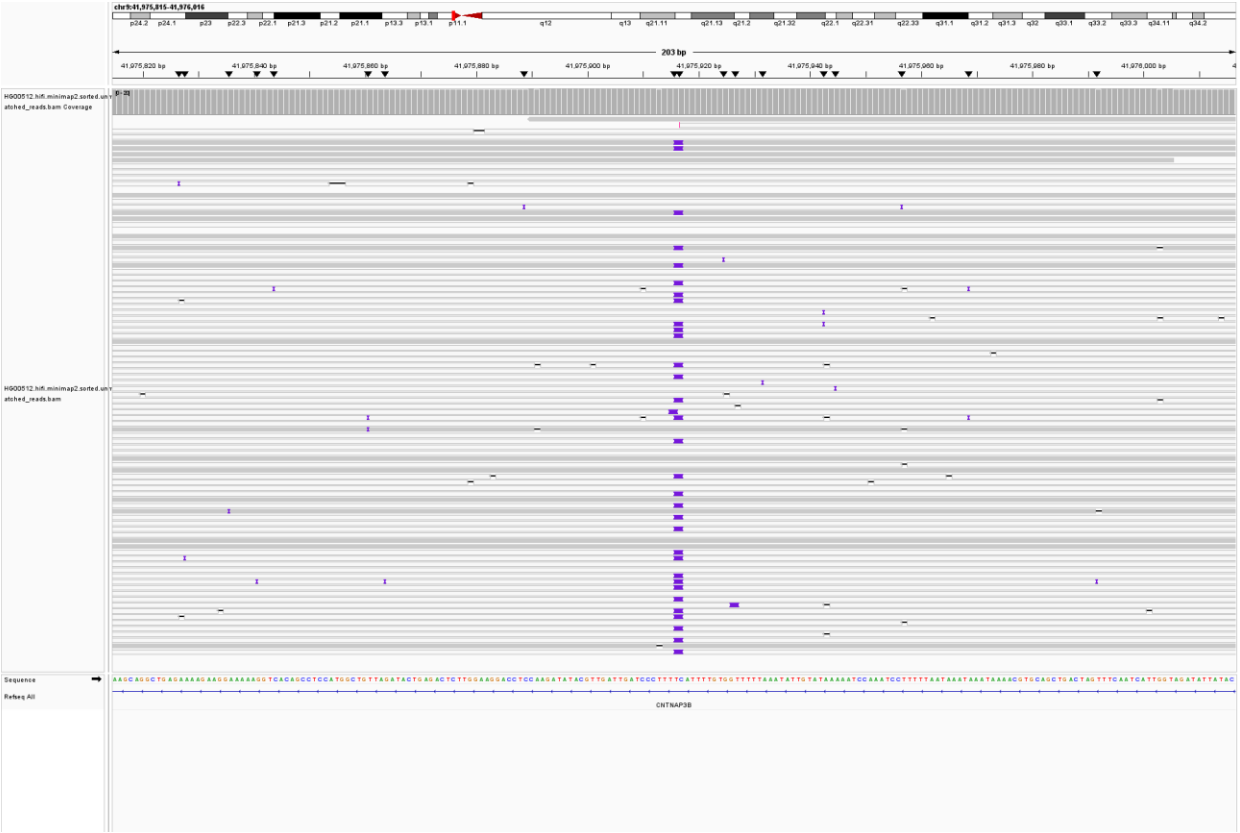

ALU-HG00512-chr9:62,644,734; (lack reads support)

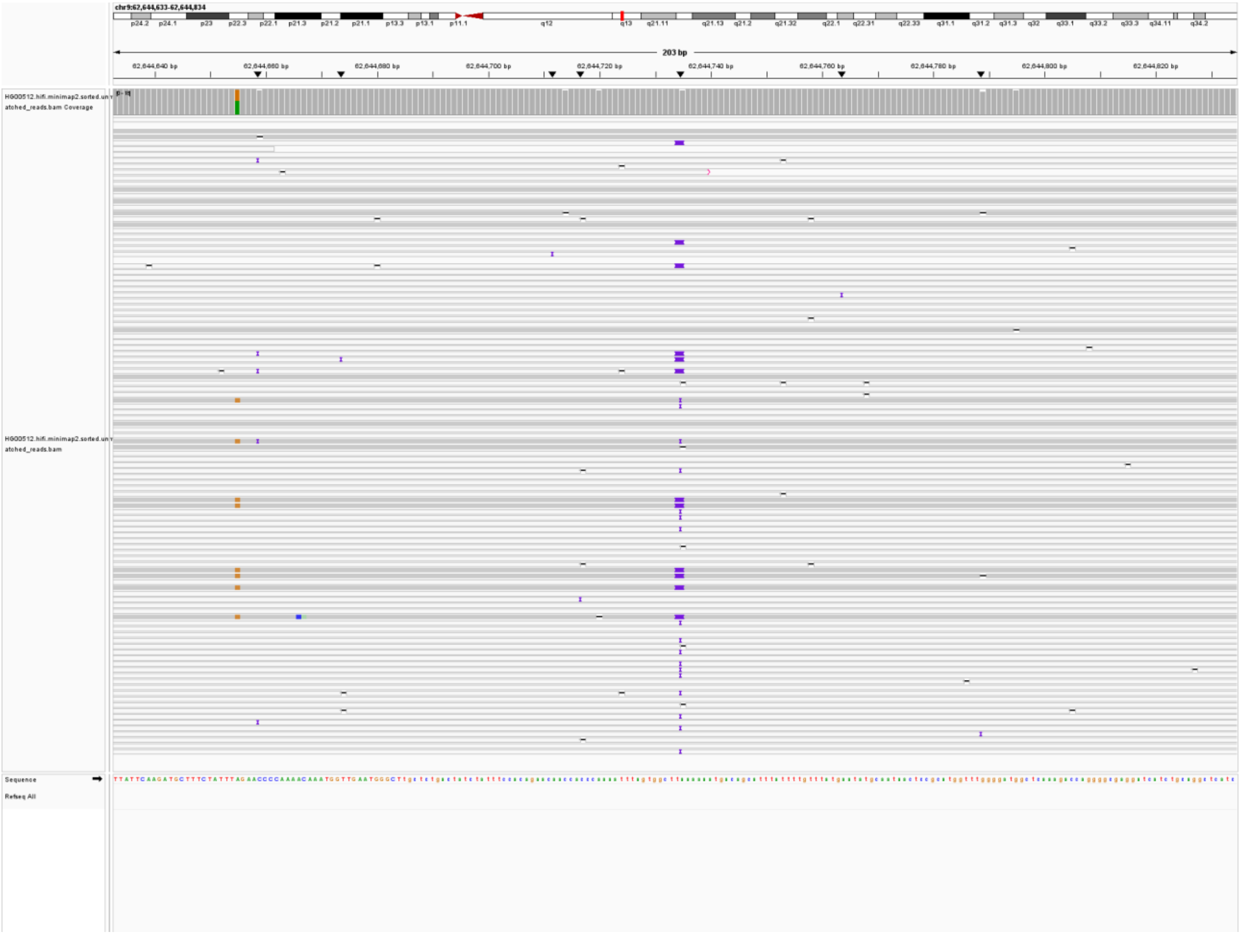

ALU-HG00512-chr9:65,302,833; (lack reads support)

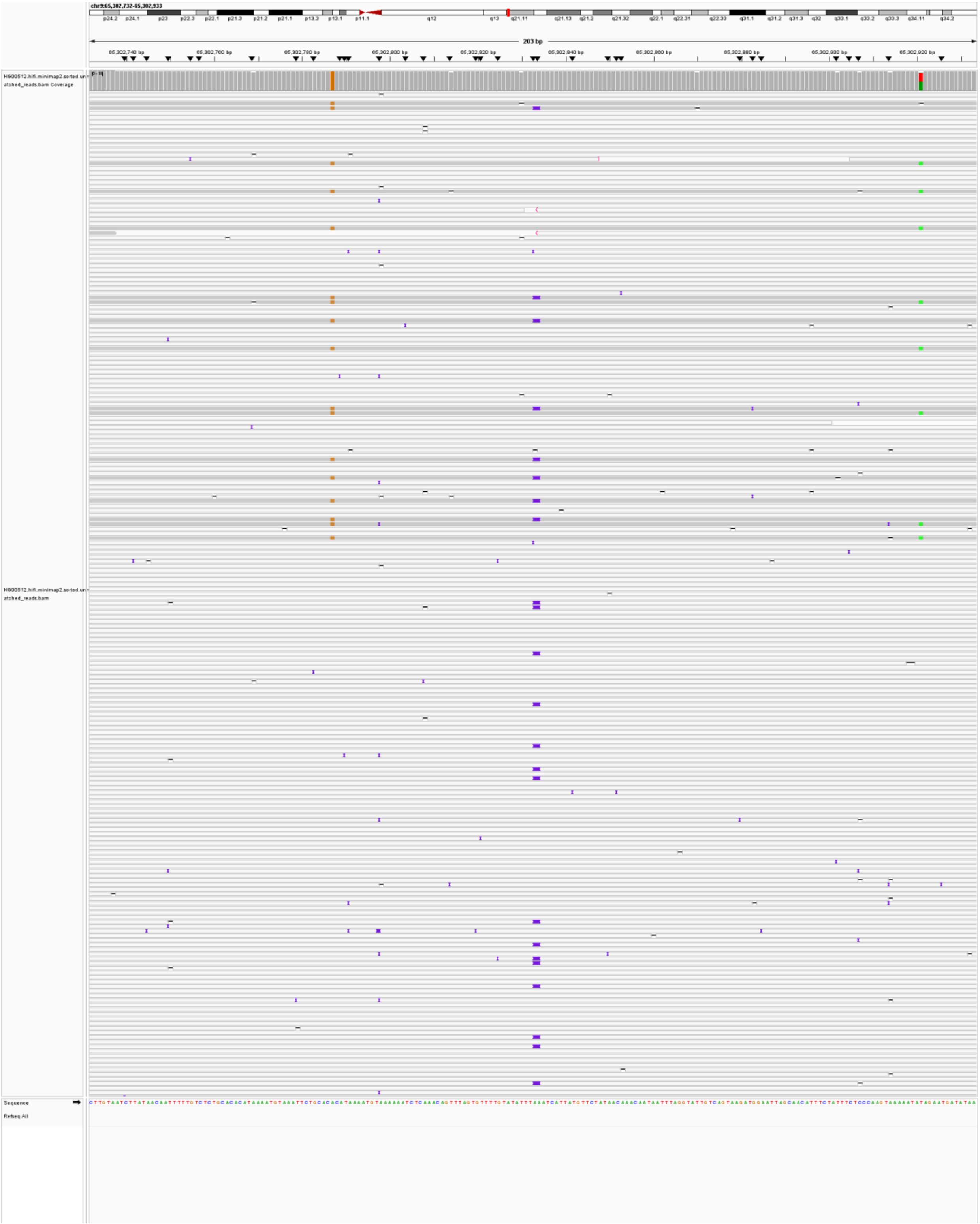

### SVA

SVA-HG00512-chr16:32,427,863; (lack reads support)

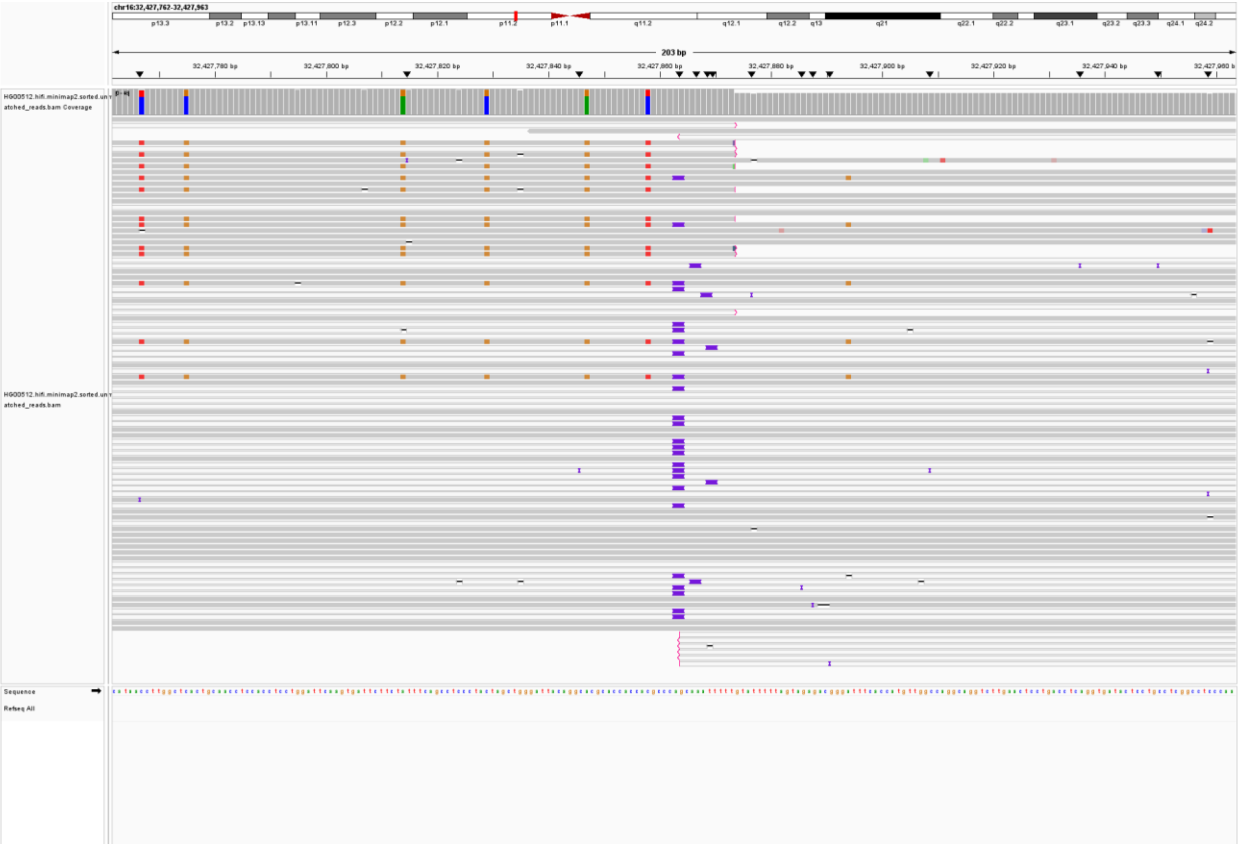

SVA-HG00512-chr16:494,827; (misclassification by MEIsensor)

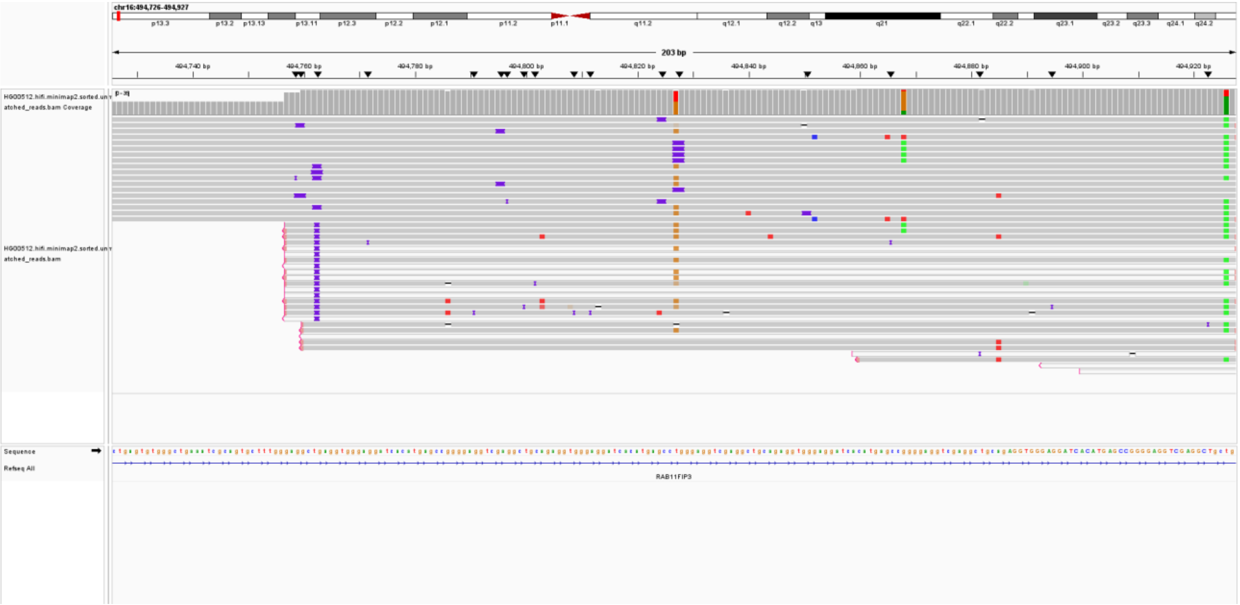

SVA-HG00512-chr19:8,422,955; (misclassification by MEIsensor)

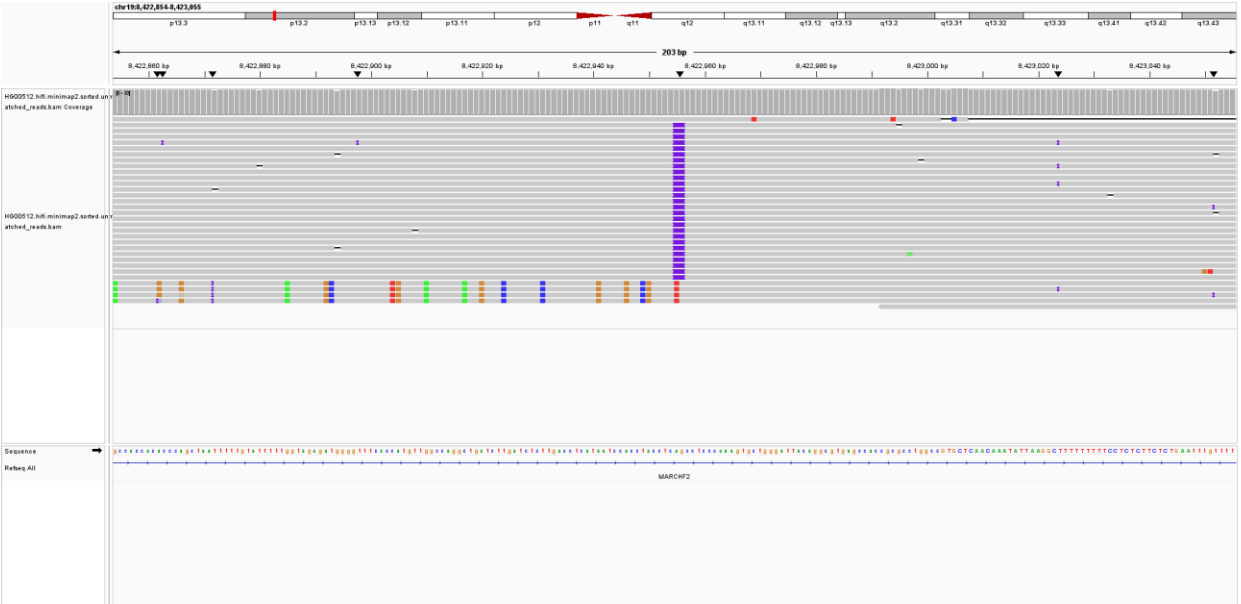

SVA-HG00512-chr1:45,637,522; (misclassification by MEIsensor)

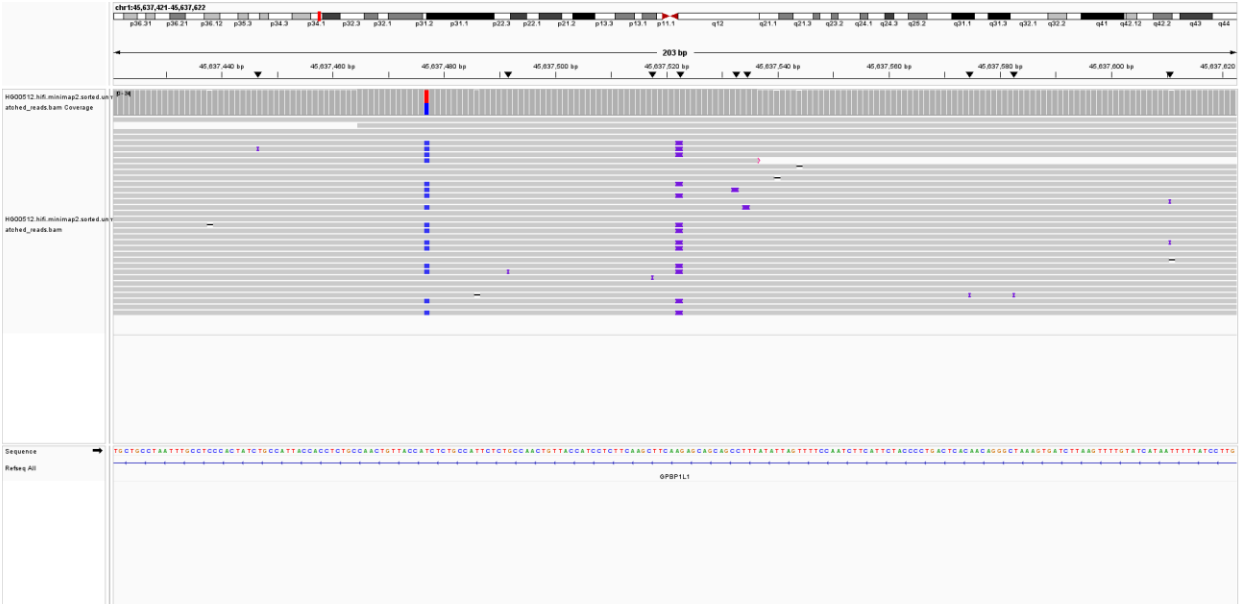

SVA-HG00512-chr21:5,694,954; (lack reads support)

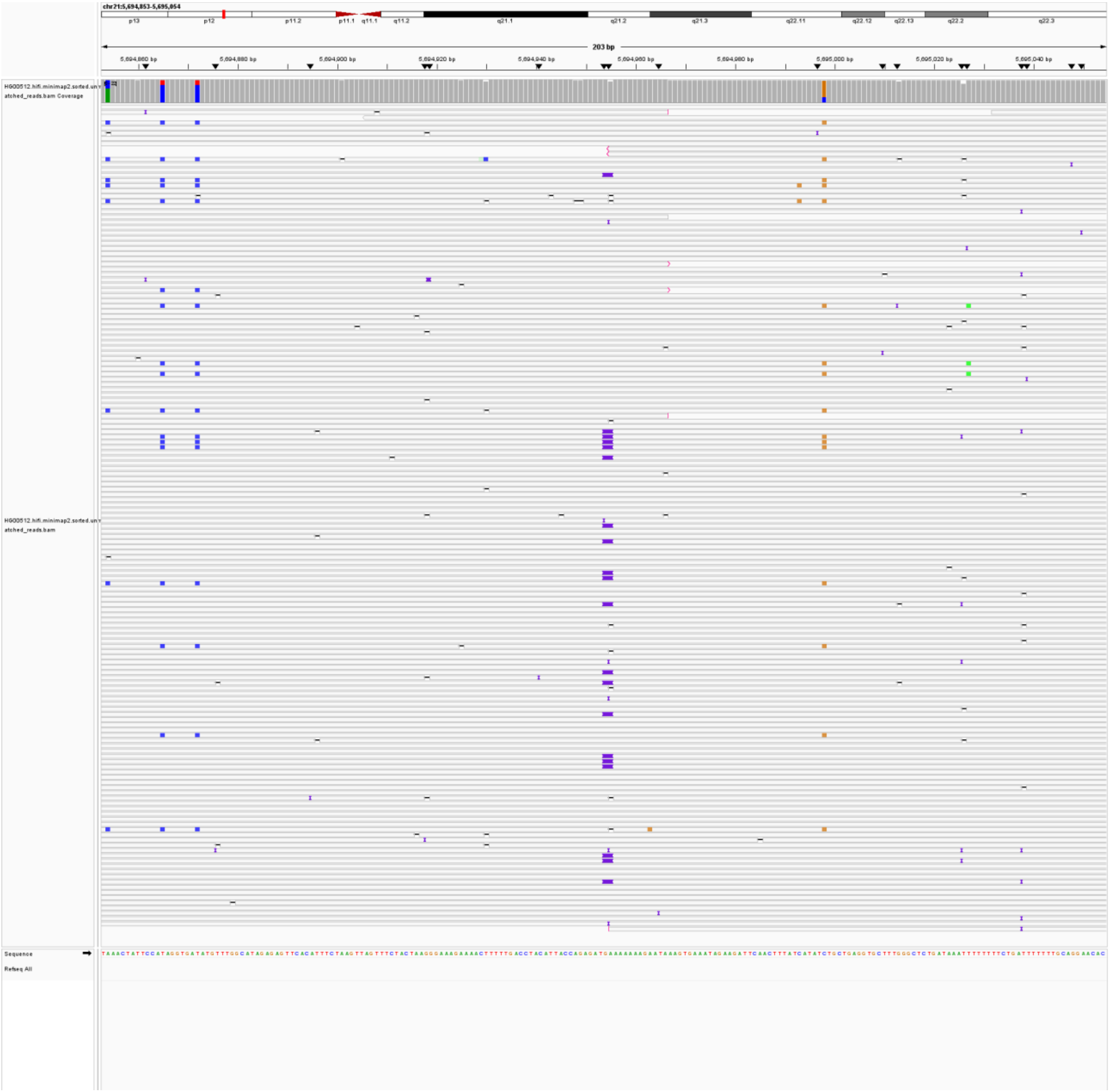

SVA-HG00512-chr22:11,296,543; (lack reads support)

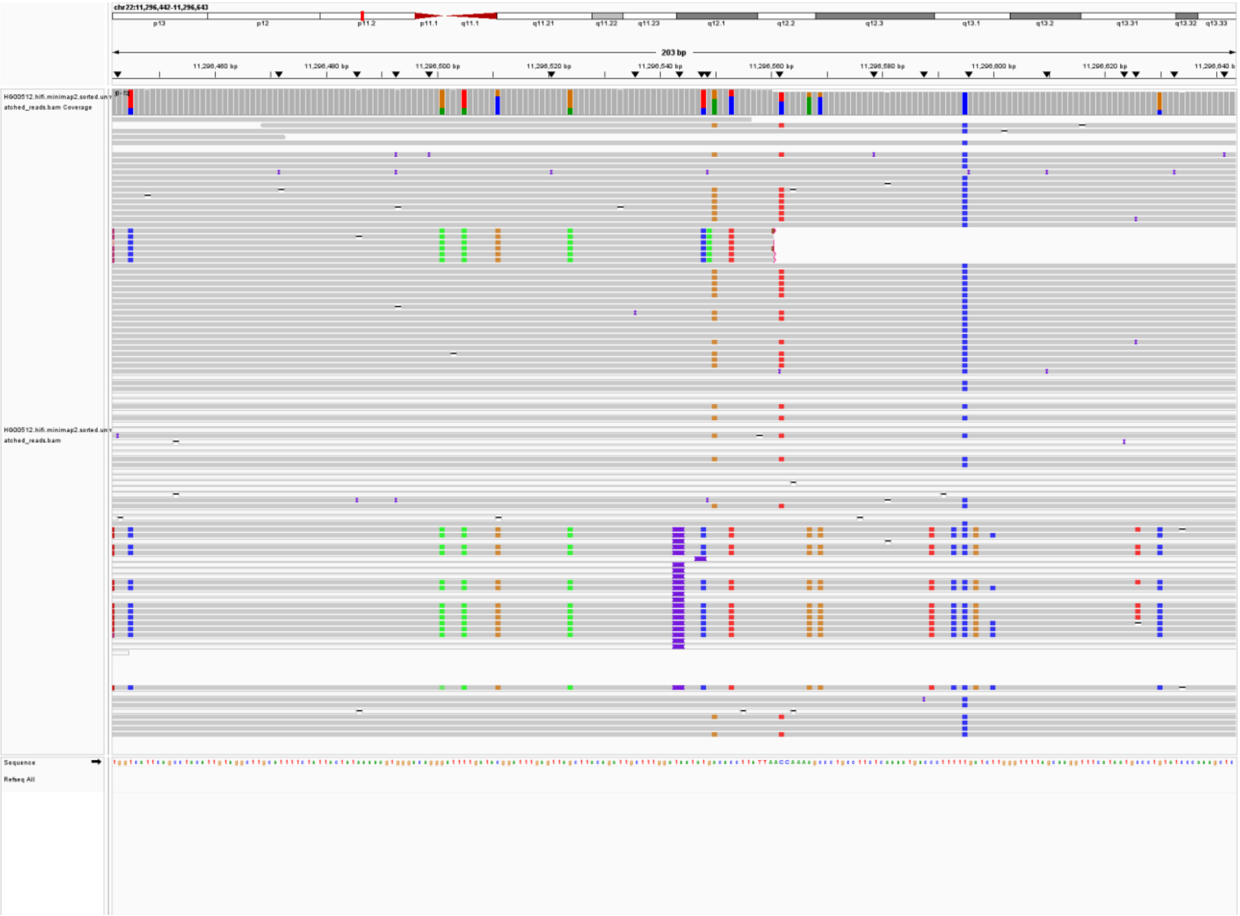

SVA-HG00512-chr6:110,956,988; (lack reads support)

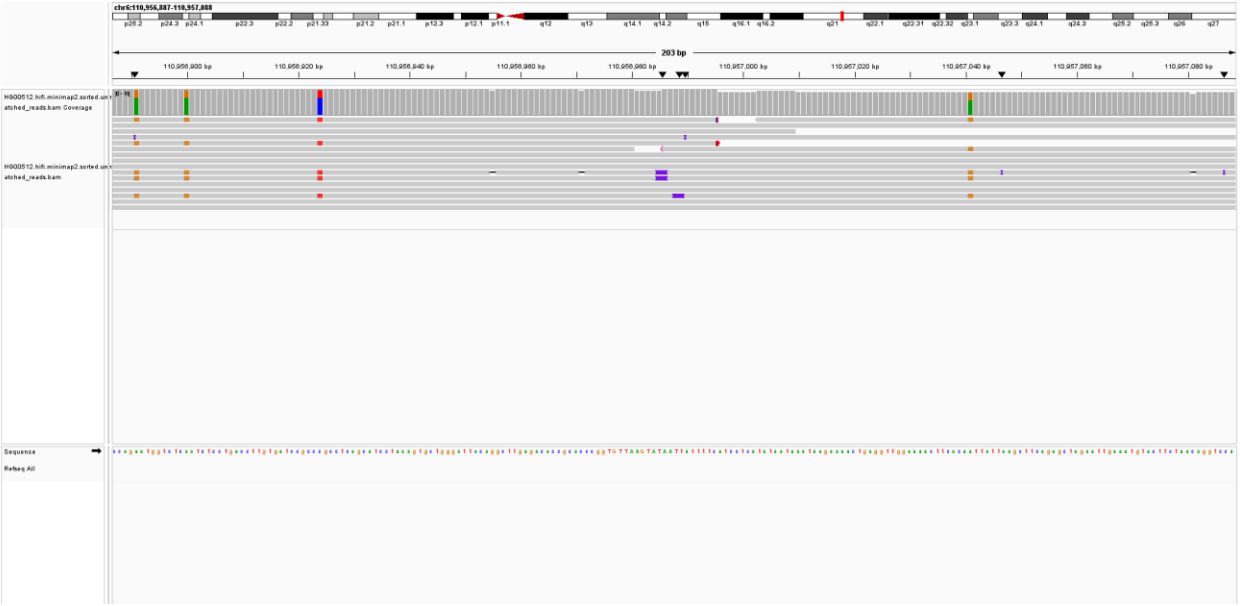

SVA-HG00512-chr6:24,683,759; (misclassification by MEIsensor)

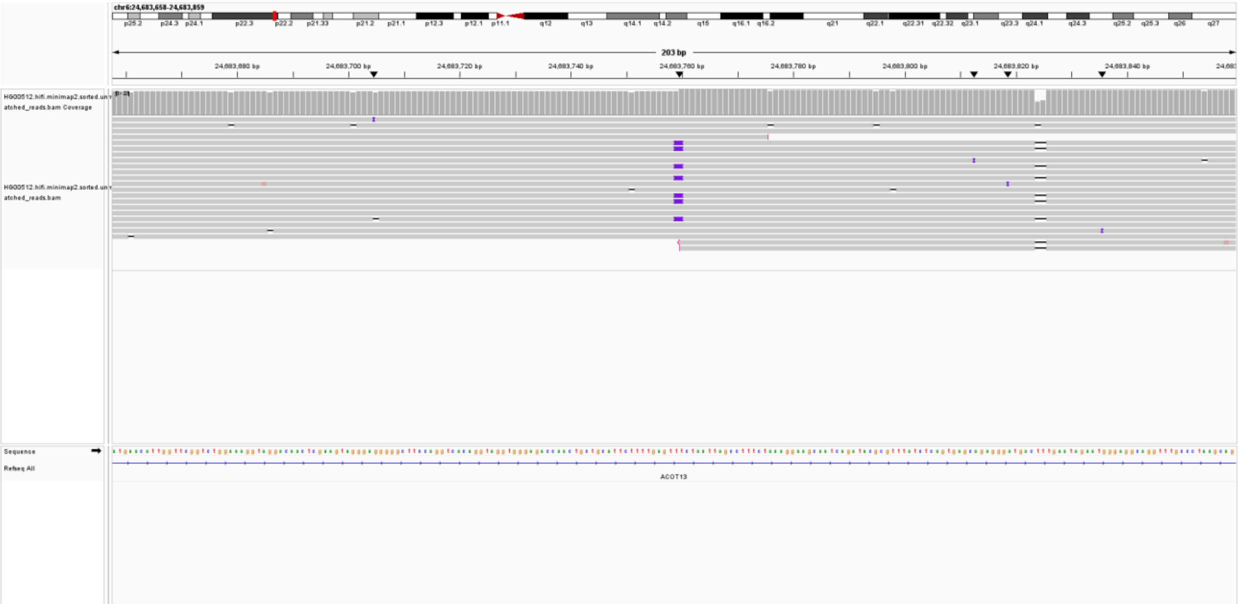

SVA-HG00512-chr7:64,197,477; (lack reads support)

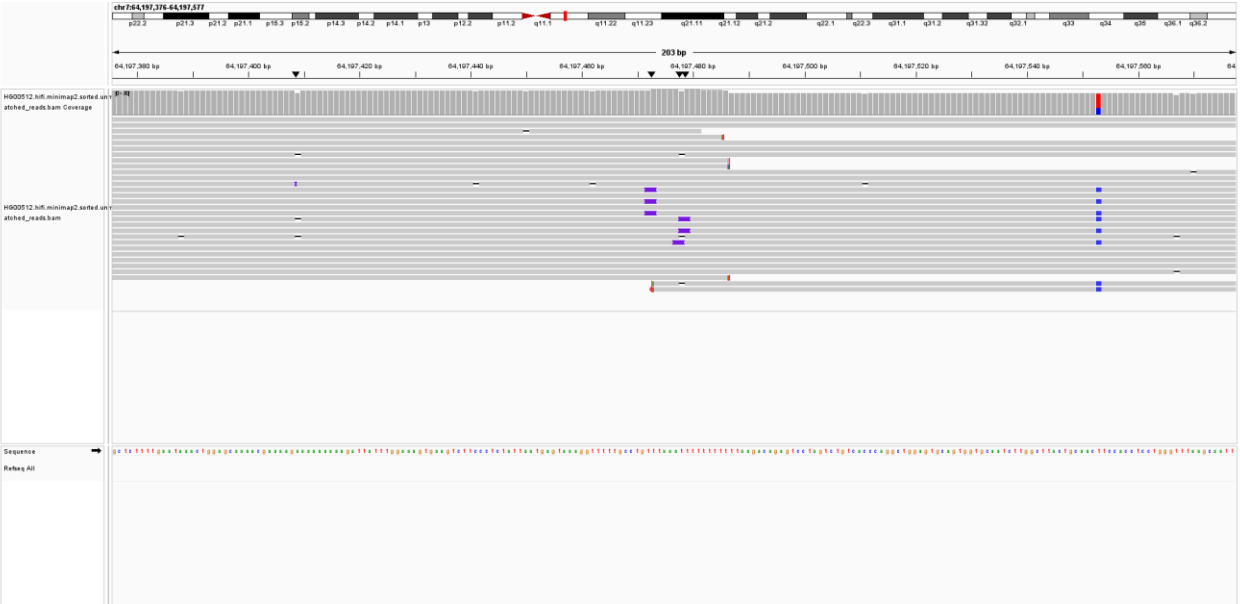
