## supplementary tables1-10, supplementary figures1-9, supplementary data1 for "MEIsensor: a deep-learning method for mobile element insertion discovery": supplementary_file.pdf

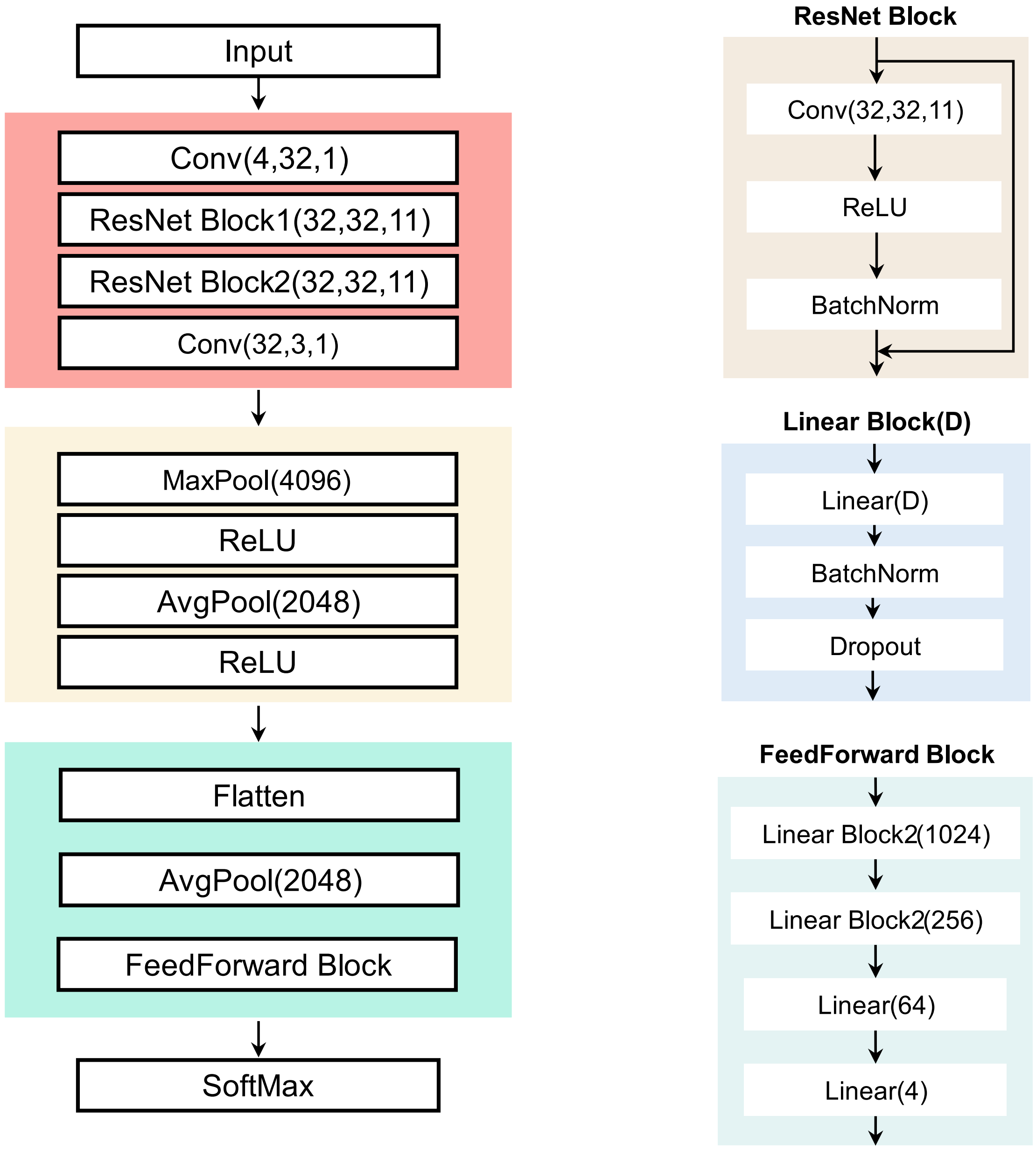


**Supplementary Fig. 1| Detailed architecture of the MEIsensor deep learning model.** The model processes one-hot encoded nucleotide sequences through a hierarchical architecture for feature extraction and classification. A 1D convolution, stacked ResNet blocks, and a transition convolution capture sequence patterns, followed by max and average pooling for dimensionality reduction and feature integration. Fully connected layers and a final SoftMax layer then classify insertions as Alu, LINE1, SVA, or background. The lower panels show the ResNet, linear, and feedforward blocks used in the network.

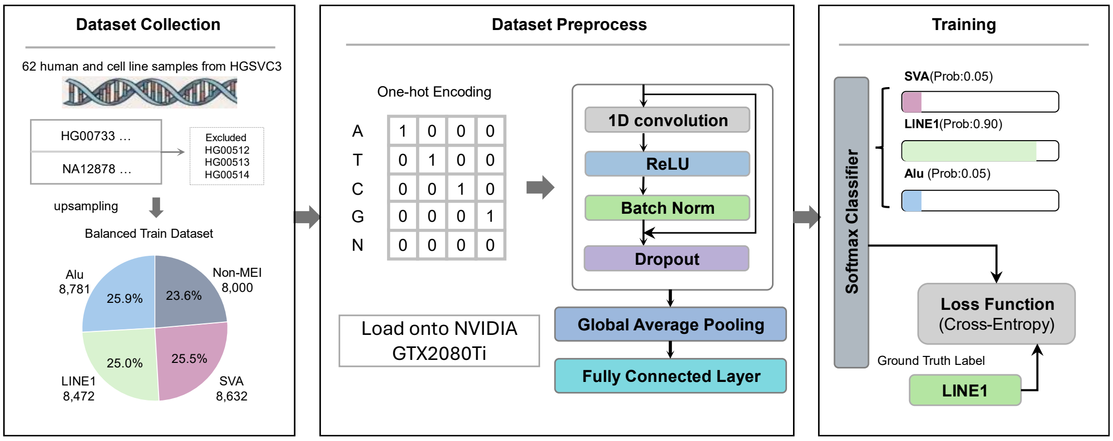


**Supplementary Fig. 2| Overview of the training procedure of the MEIsensor deep learning model.**

The left panel shows dataset collection and class balancing. MEI sequences were obtained from 62 human and cell-line samples from HGSVC3, with HG00512, HG00513, and HG00514 excluded from model development and reserved for independent evaluation. The remaining samples were used to construct the training dataset, and minority classes, including LINE1 and SVA insertions, were upsampled to reduce class imbalance relative to Alu and non-MEI insertions. The middle panel illustrates sequence preprocessing and model architecture. Insertion sequences were converted into one-hot encoded representations and processed by a one-dimensional convolutional neural network composed of residual blocks, ReLU activation, batch normalization, dropout, global average pooling, and a fully connected layer. Both encoded inputs and model computation were loaded onto an NVIDIA GTX 2080 Ti GPU for training. The right panel shows supervised optimization, in which the extracted features were passed through a SoftMax classifier to generate class probabilities for Alu, LINE1, and SVA insertions, and model parameters were optimized using cross-entropy loss against ground-truth labels.

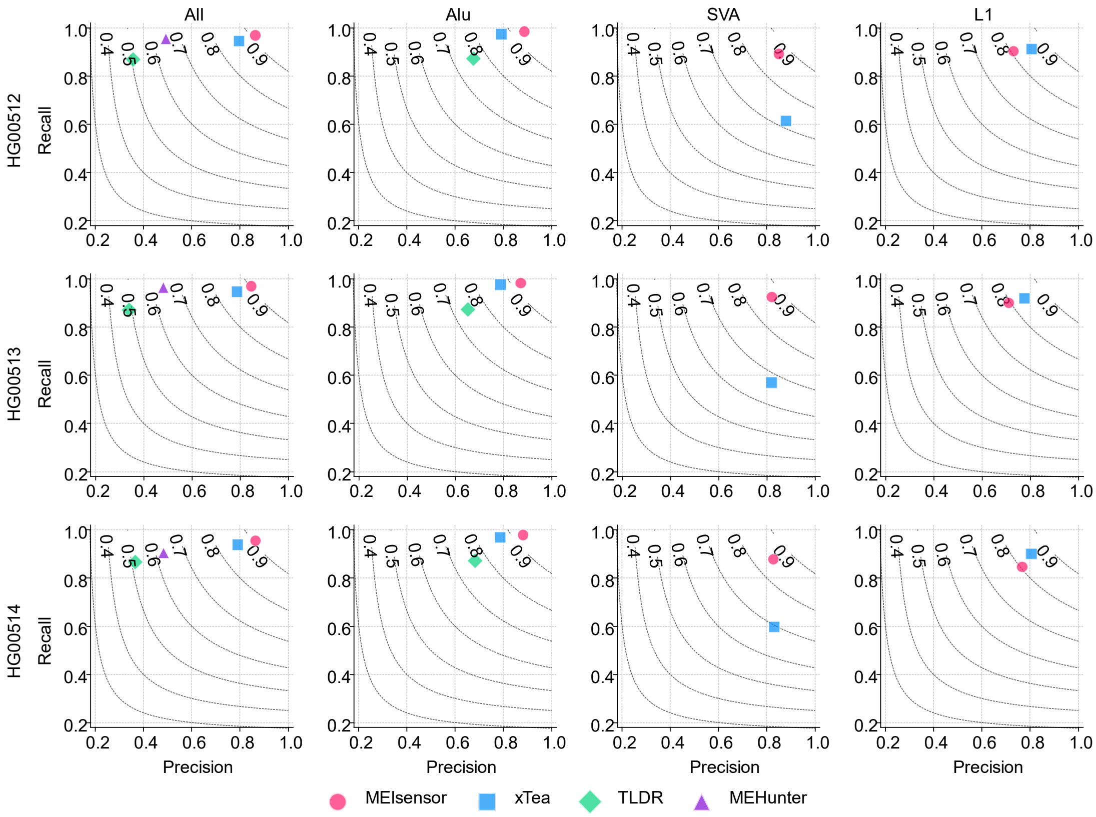


**Supplementary Fig. 3| Detection performance of MEIsensor, xTea, TLDR, and MEHunter onHG00512, HG00513, and HG00514.**

Precision–recall (PR) plots are shown separately for each sample, with panels reporting results for all MEIs, and Alu, LINE1, and SVA insertions. Each point denotes the overall precision and recall under a unified filtering scheme; dashed curves indicate iso-F1 contours.

LINE1-HG00512-chr1:125,012,475

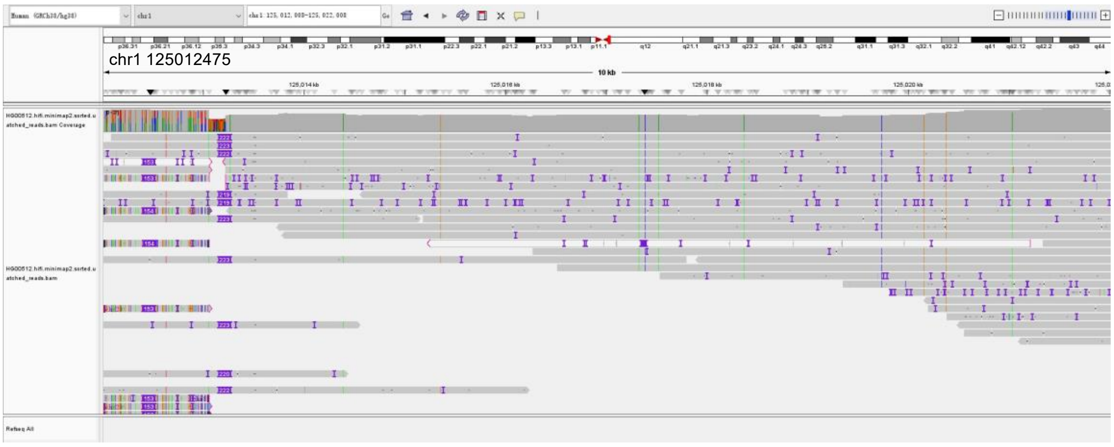


LINE1-HG00512-chr1:13,161,043

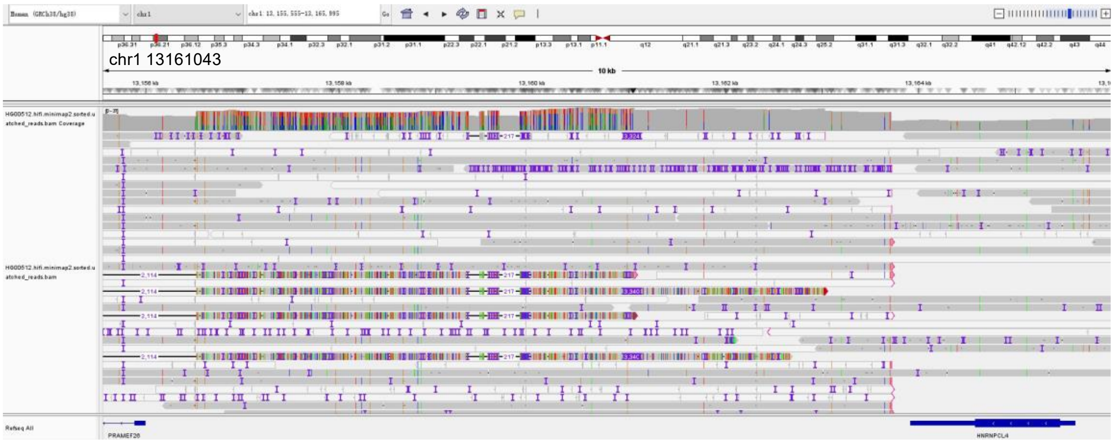


LINE1-HG00512-chr15:45,026,631

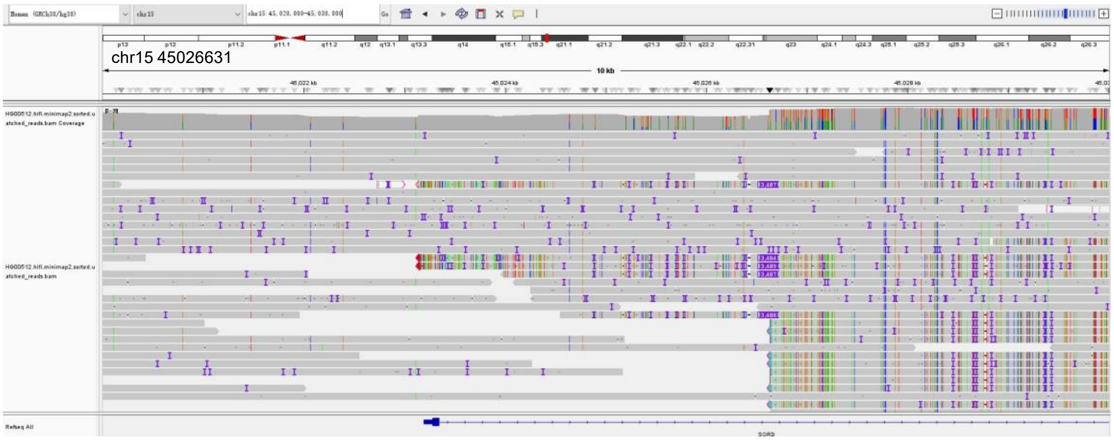

LINE1-HG00512-chr20:28,576,053

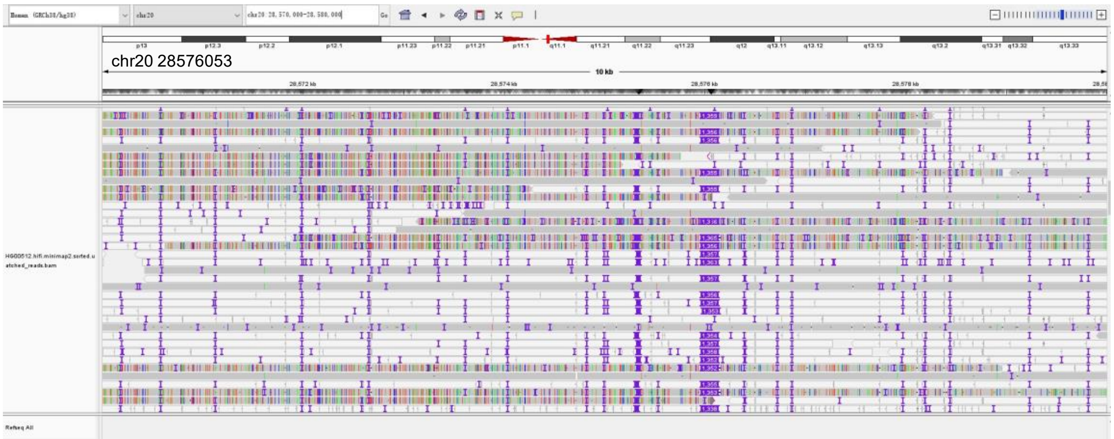


LINE1-HG00512-chr17:21,828,255

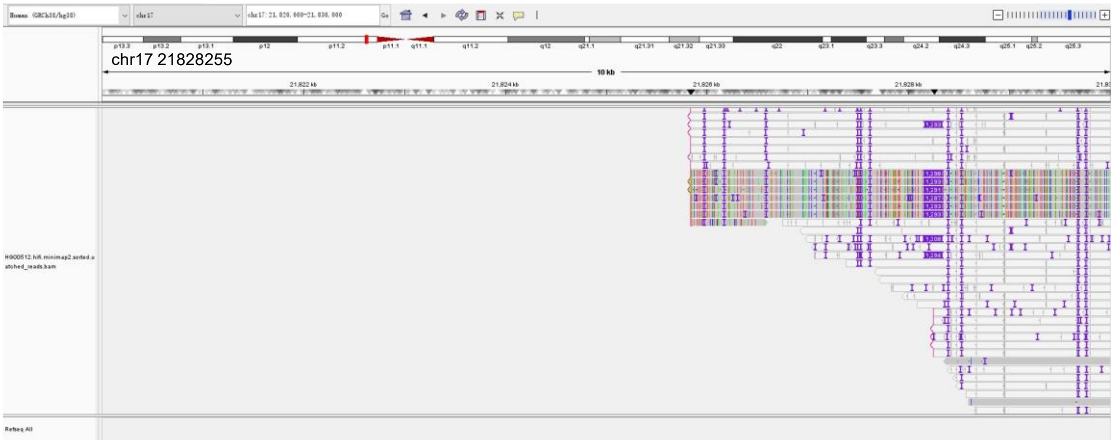


LINE1-HG00512-chr20:29,679,453

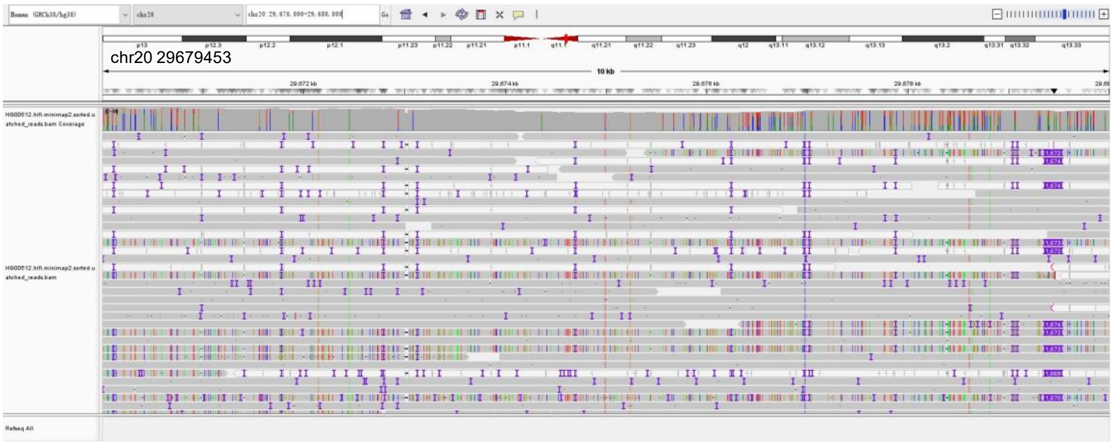

LINE1-HG00512-chr20:30,177,003

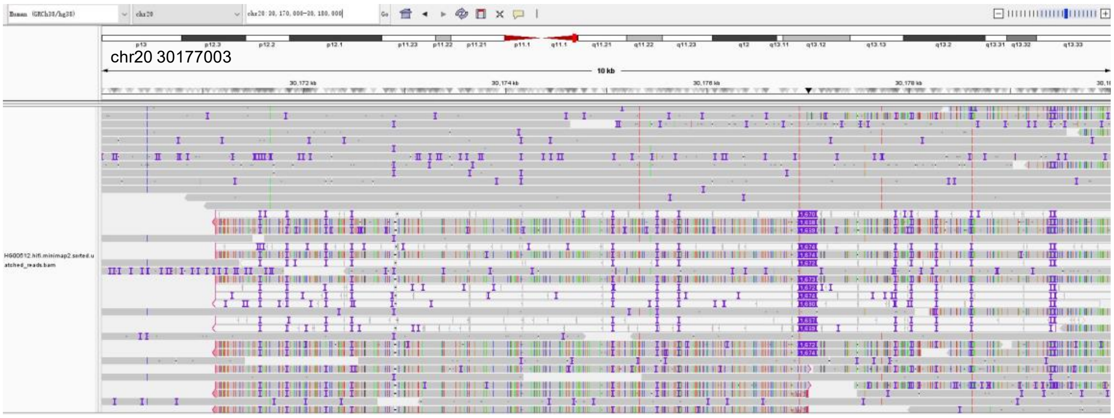


LINE1-HG00512-chr21:7,202,316

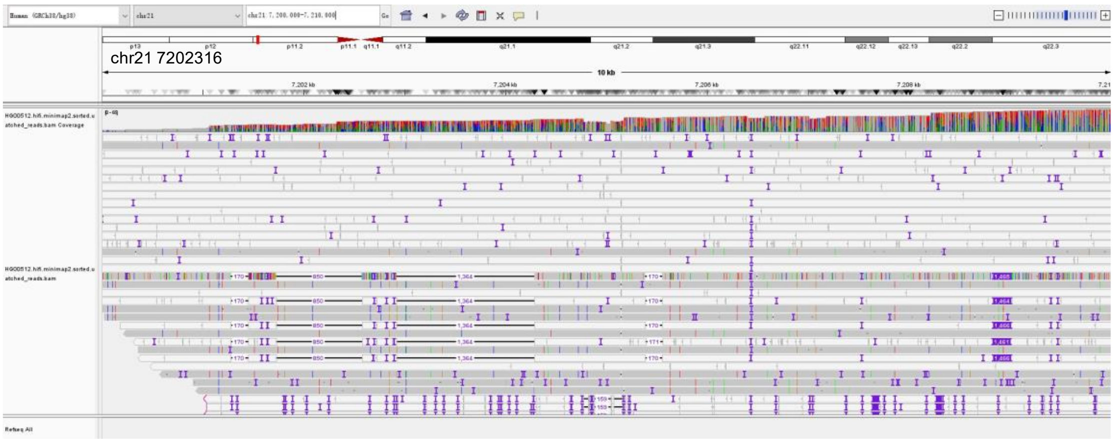


LINE1-HG00512-chr21:7,967,836

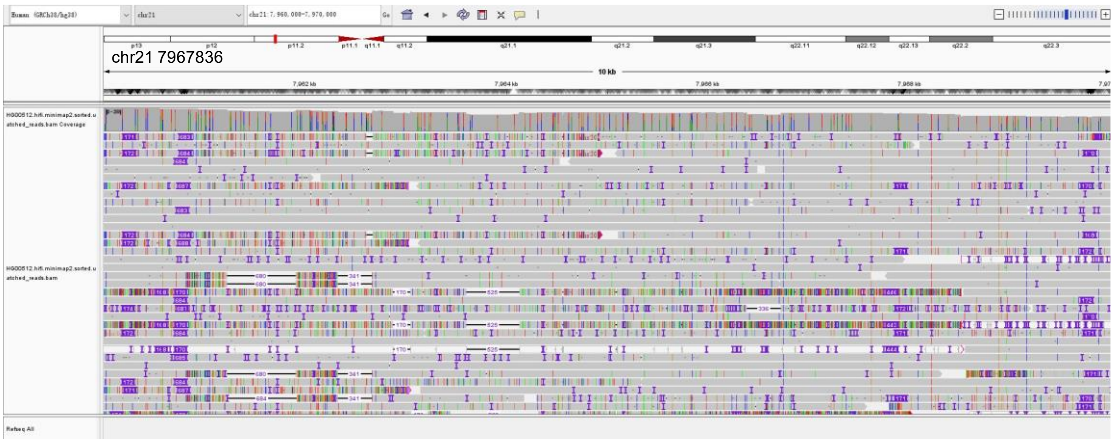

LINE1-HG00512-chr22:10,934,745

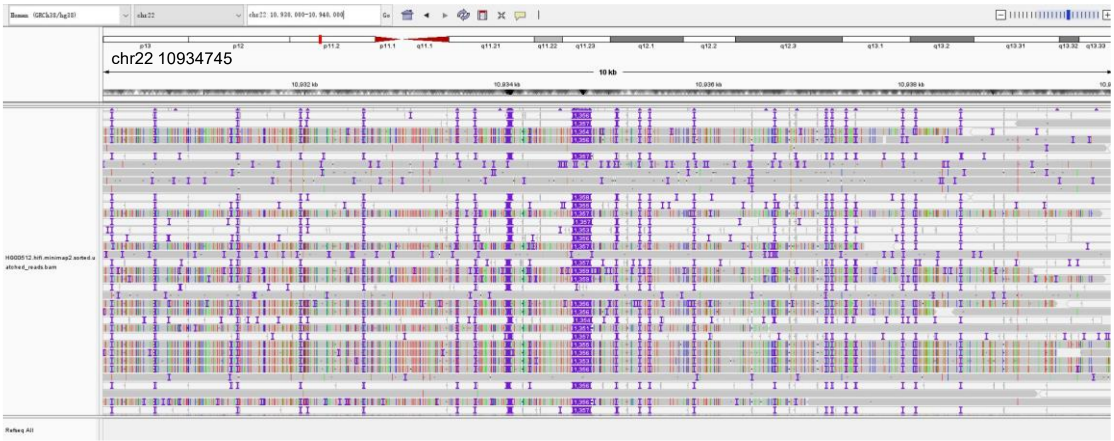


LINE1-HG00512-chr4:65,548,212

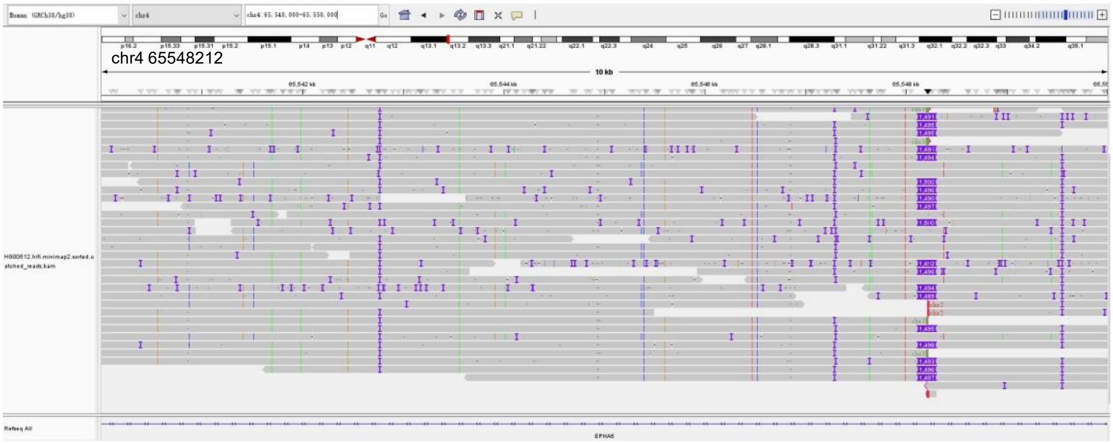


LINE1-HG00512-chr4:189,978,686

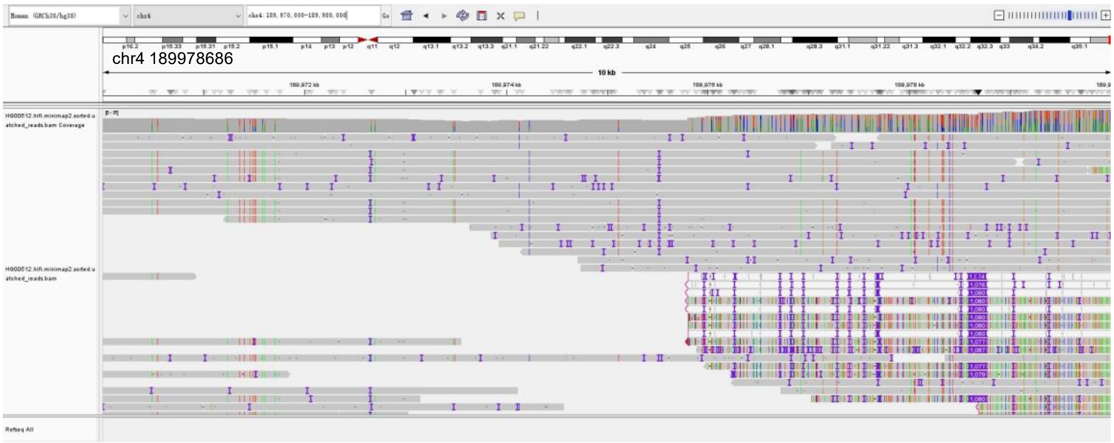

LINE1-HG00512-chr5:49,736,601

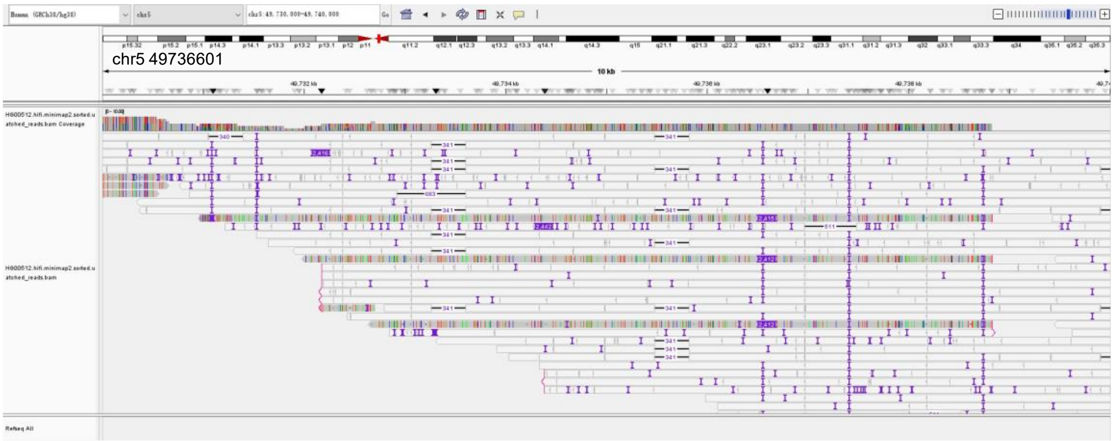


LINE1-HG00512-chr7:34,800,834

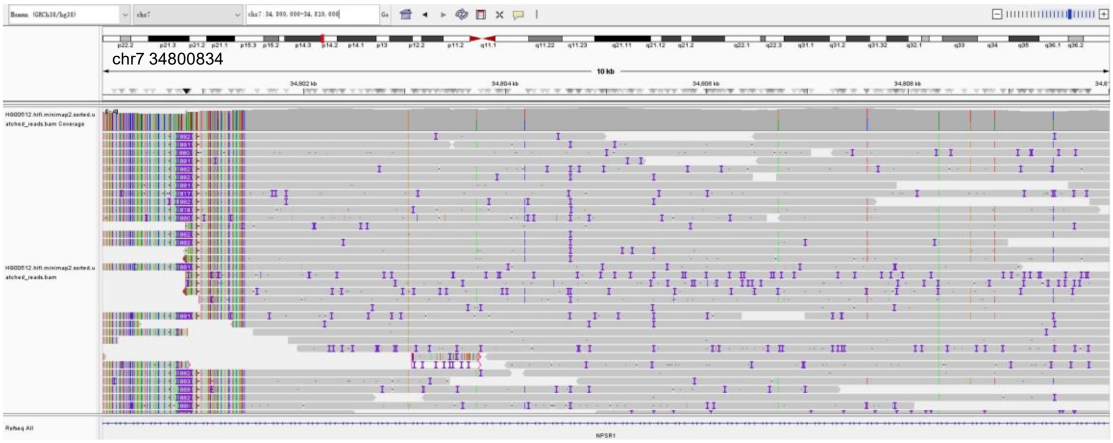


LINE1-HG00512-chr7:62,863,673

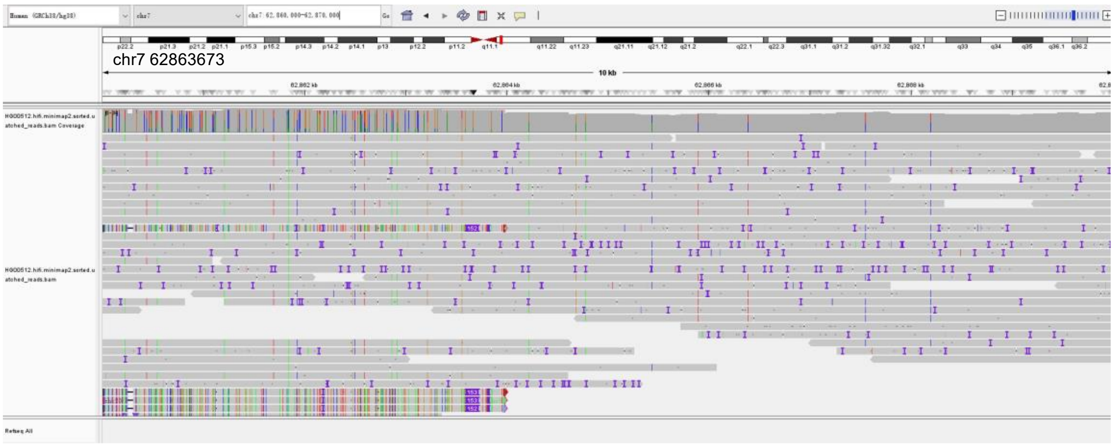

LINE1-HG00512-chr7:123,446,450

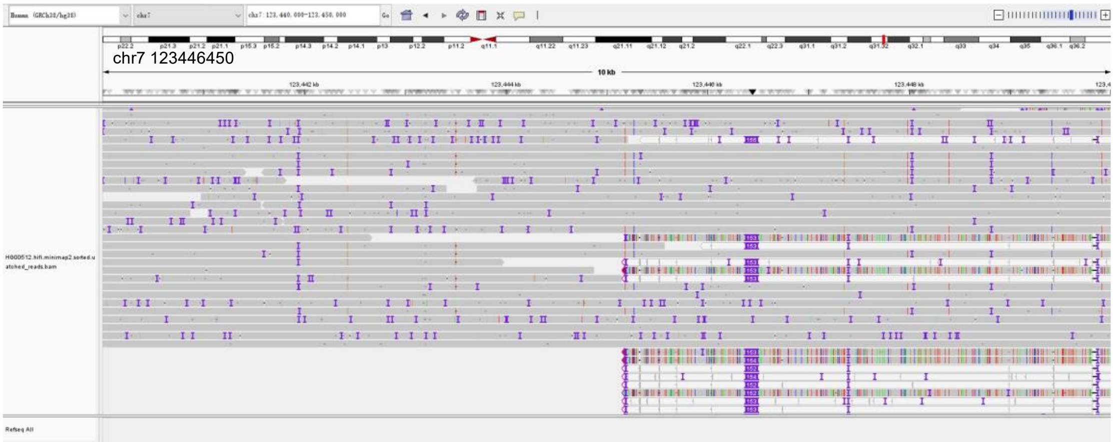


LINE1-HG00512-chr8:127,521,588

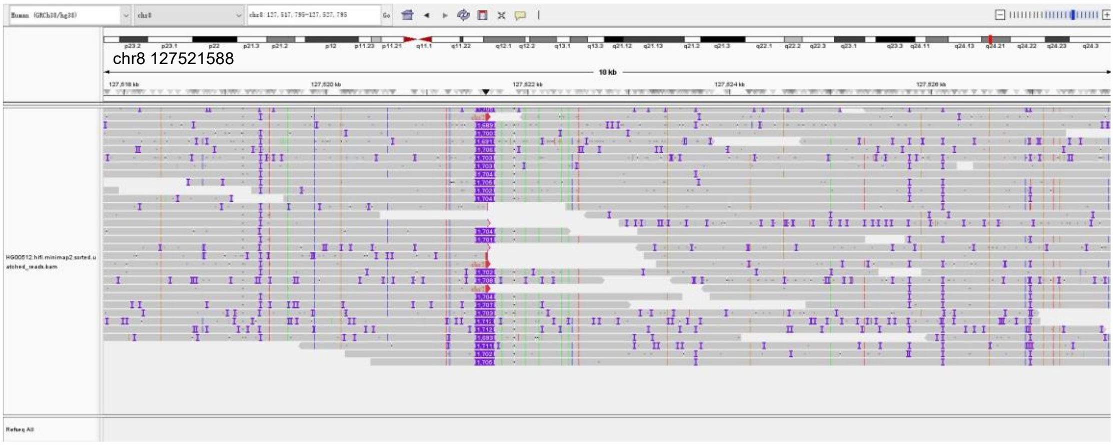


LINE1-HG00512-chrX:25,885,905

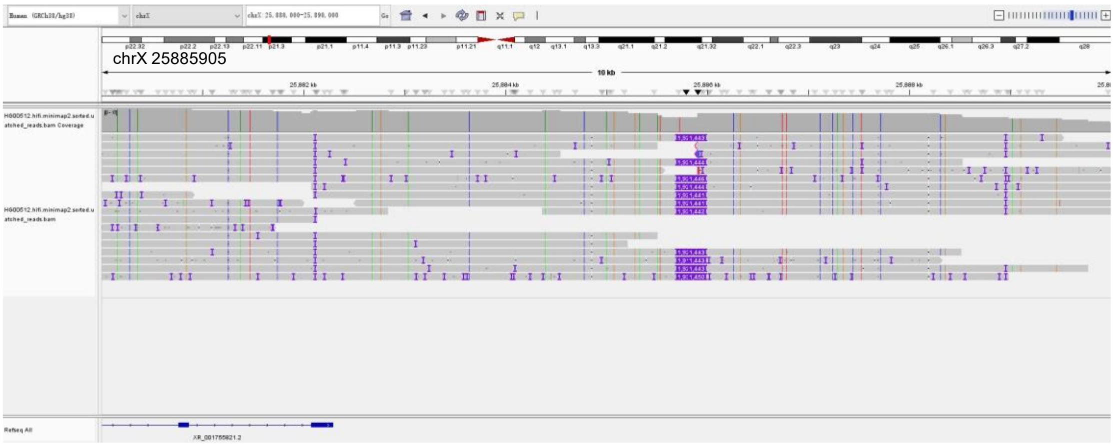


**Supplementary Fig. 4 | IGV validation of supplemented LINE1 insertions.**

IGV visualizations of long-read alignments supporting LINE1 insertions detected by MEIsensor but absent from the original benchmark dataset, consistent with bona fide insertion events missed during benchmark construction.

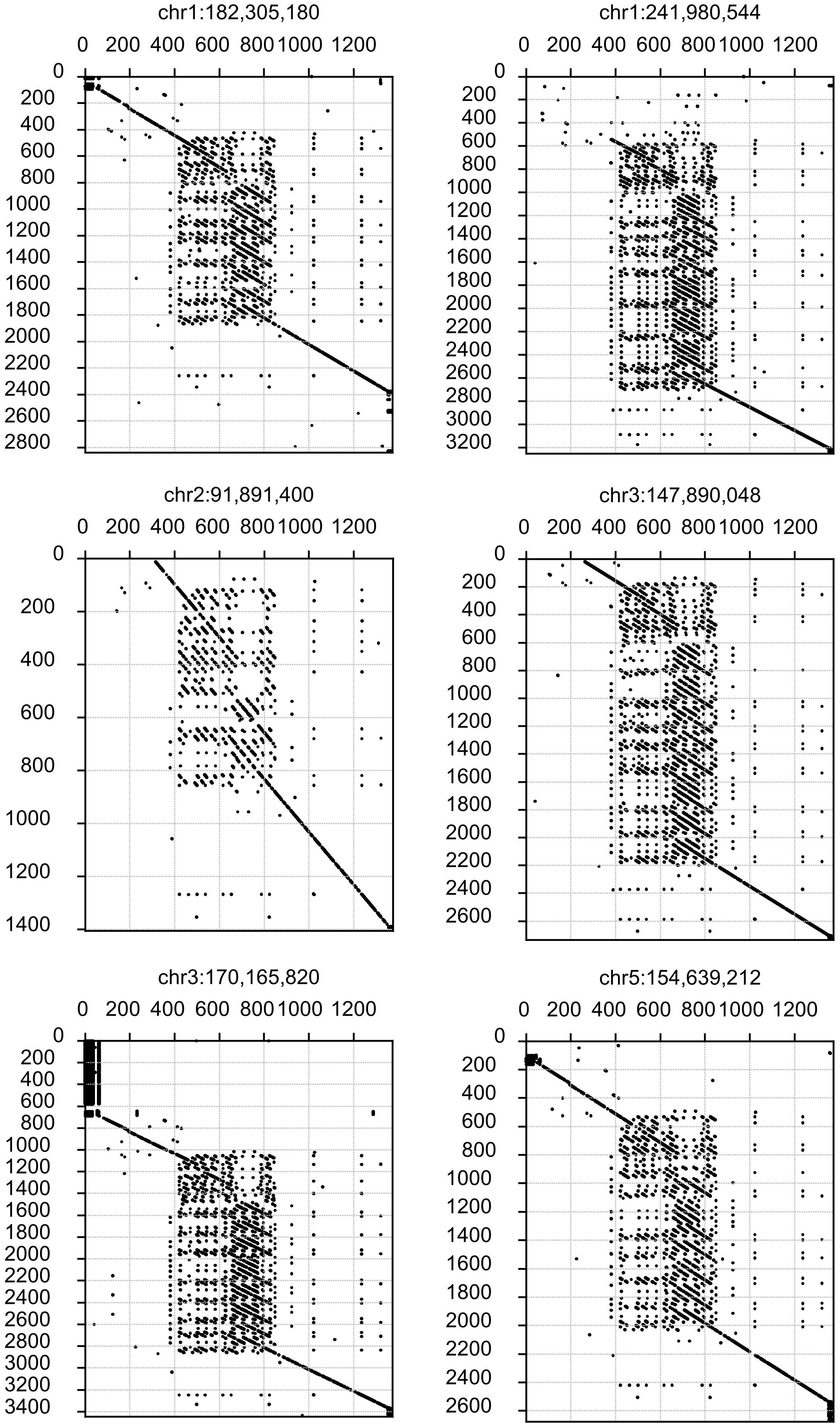
chr1:182,305,180
chr1:241,980,544
chr2:91,891,400
chr3:147,890,048
chr3:170,165,820
chr5:154,639,212

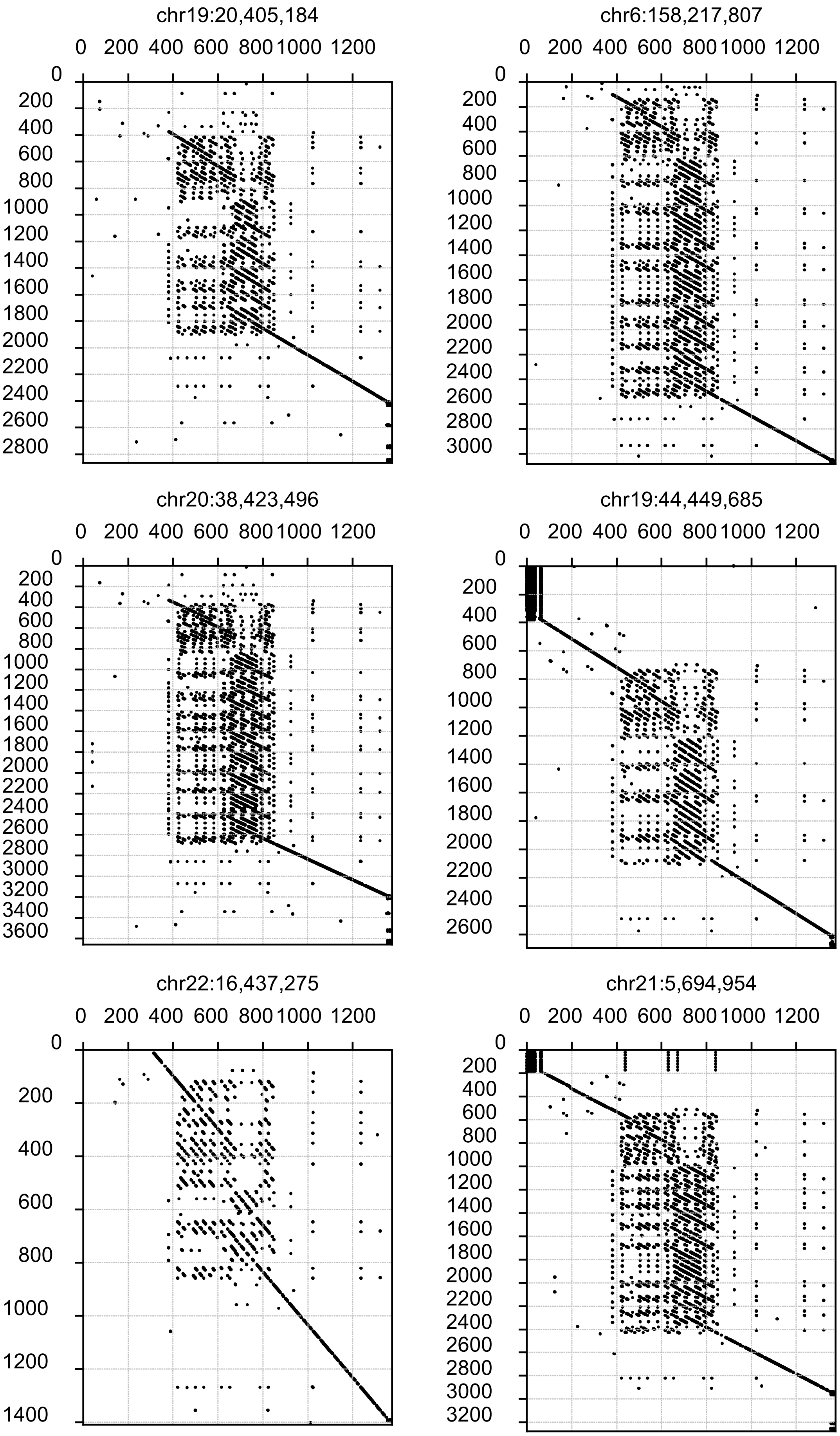
chr19:20,405,184
0 200 400 600 800 1000 1200
0 200 400 600 800 1000 1200 1400 1600 1800 2000 2200 2400 2600 2800
chr6:158,217,807
0 200 400 600 800 1000 1200
0 200 400 600 800 1000 1200 1400 1600 1800 2000 2200 2400 2600 2800 3000
chr20:38,423,496
0 200 400 600 800 1000 1200
0 200 400 600 800 1000 1200 1400 1600 1800 2000 2200 2400 2600 2800 3000 3200 3400 3600
chr19:44,449,685
0 200 400 600 800 1000 1200
0 200 400 600 800 1000 1200 1400 1600 1800 2000 2200 2400 2600
chr22:16,437,275
0 200 400 600 800 1000 1200
0 200 400 600 800 1000 1200 1400
chr21:5,694,954
0 200 400 600 800 1000 1200
0 200 400 600 800 1000 1200 1400 1600 1800 2000 2200 2400 2600 2800 3000 3200

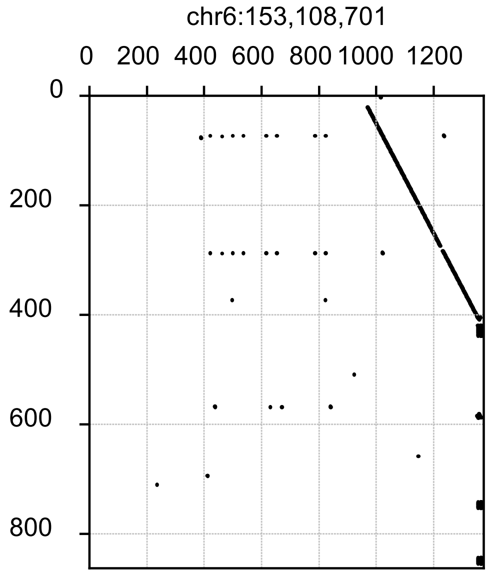


**Supplementary Fig. 5 | Dotplot of 12 complex SVA insertions and 1 simple SVA insertion uniquely detected by MEIsensor against the SVA consensus sequence.**

Dotplots of SVA insertions detected by MEIsensor but missed by xTea. The x-axis shows the SVA consensus sequence from Repbase, and the y-axis shows the inserted variant sequence. Upper panels represent structurally complex SVA insertions, while lower panels represent structurally simple SVA insertions. Shared *k*-mer matches reveal rearrangements and internal structural complexity, illustrating the ability of MEIsensor to recover complex SVA insertions.

ALU-HG00512-chr14:106,357,640

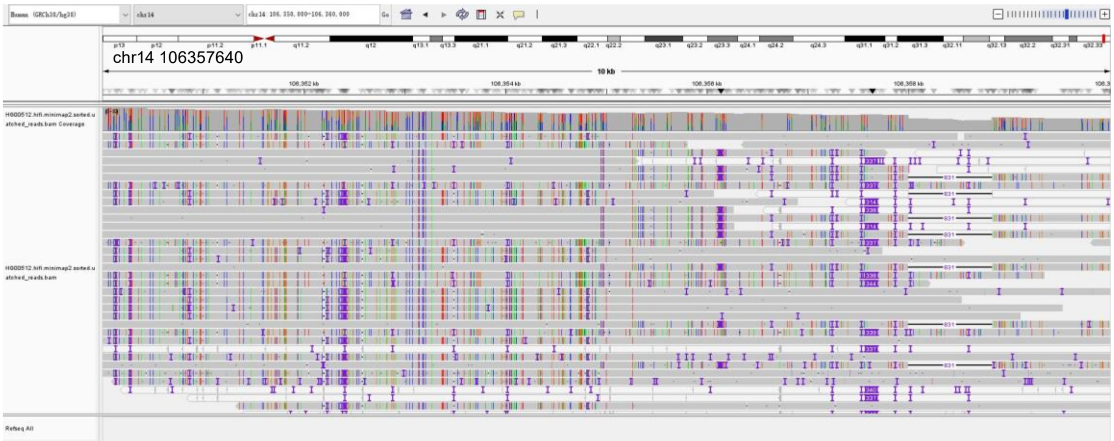


ALU-HG00512-chr16:21,850,494

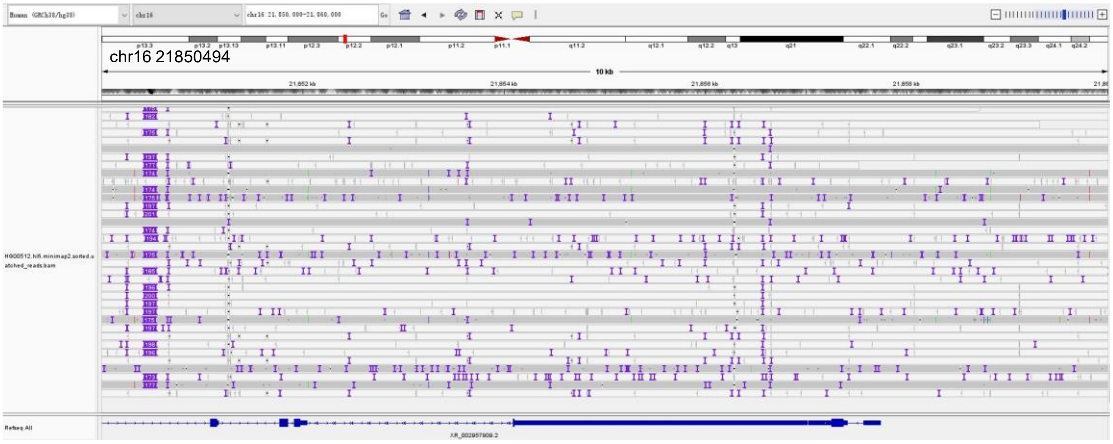


ALU-HG00512-chr10:80,636,473

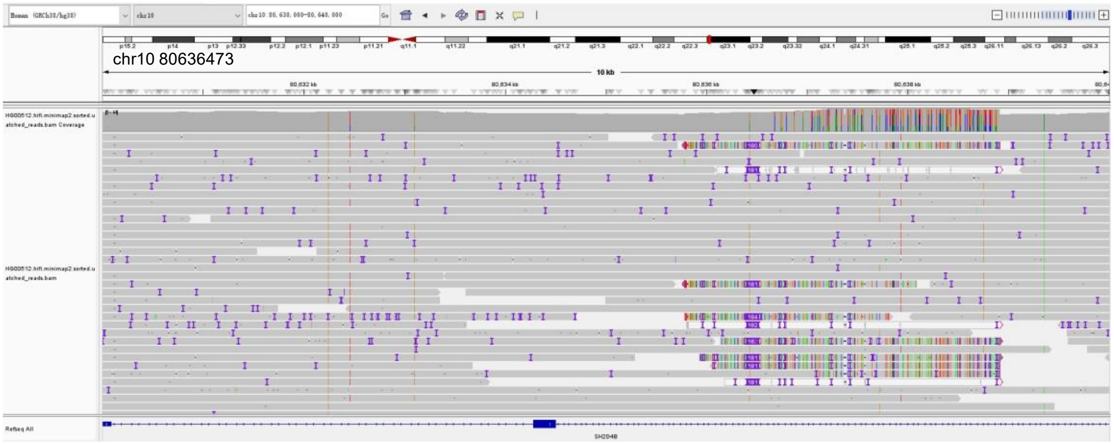

ALU-HG00512-chr15:20,238,878

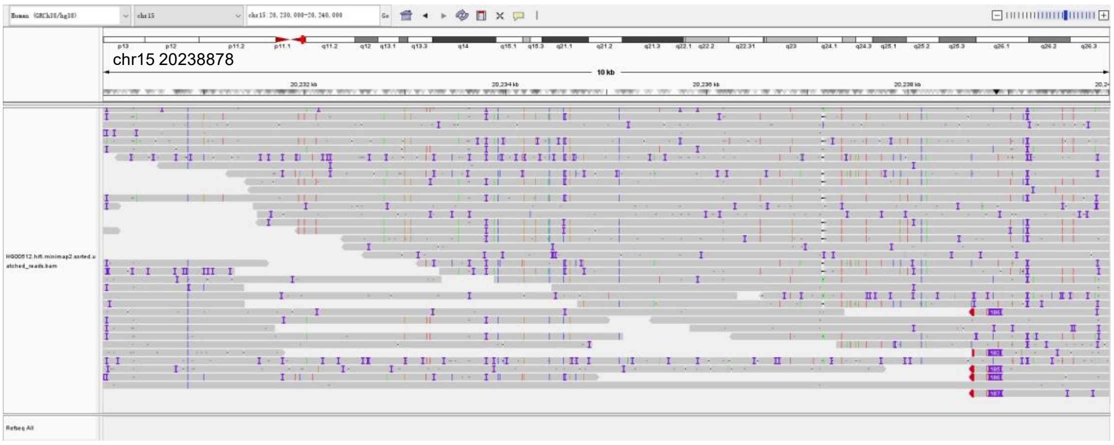


ALU-HG00512-chr17:38,076,286

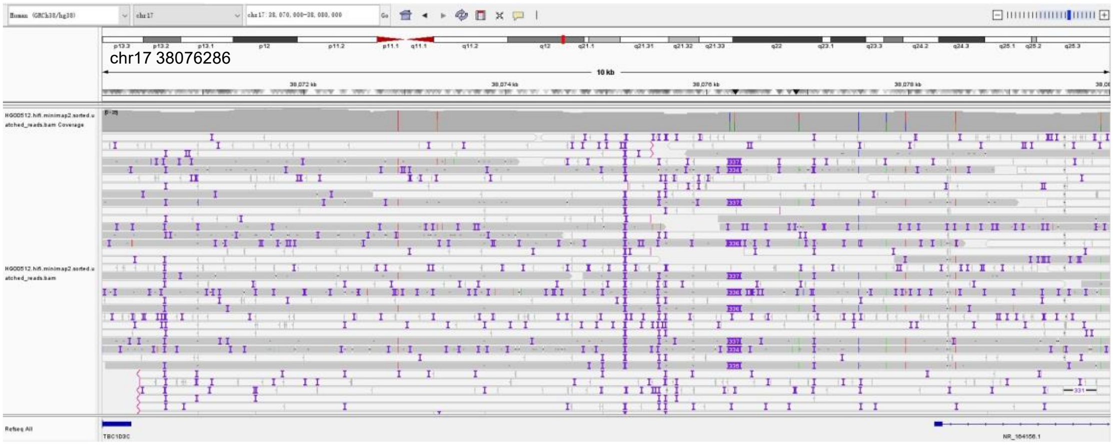


ALU-HG00512-chr20:29,885,837

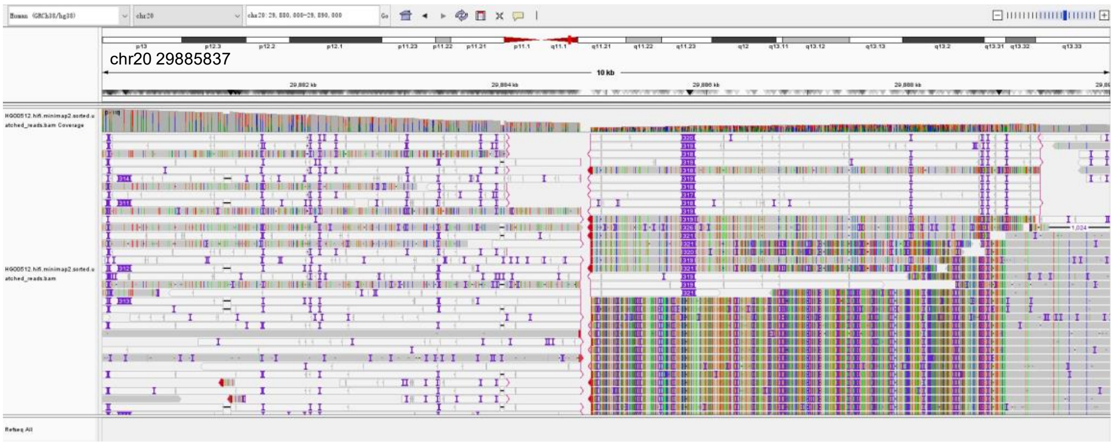

ALU-HG00512-chr21:5,312,285

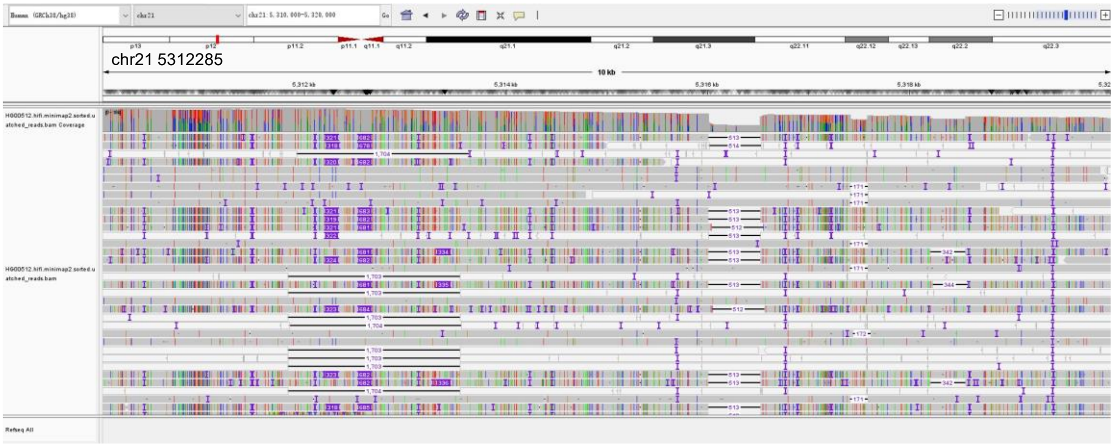


ALU-HG00512-chr21:20,075,431

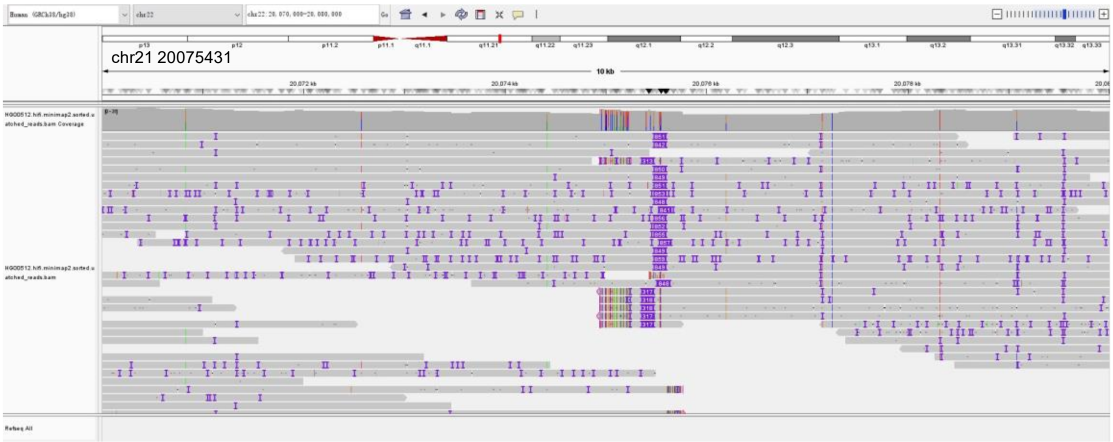


ALU-HG00512-chr21:130,090,881

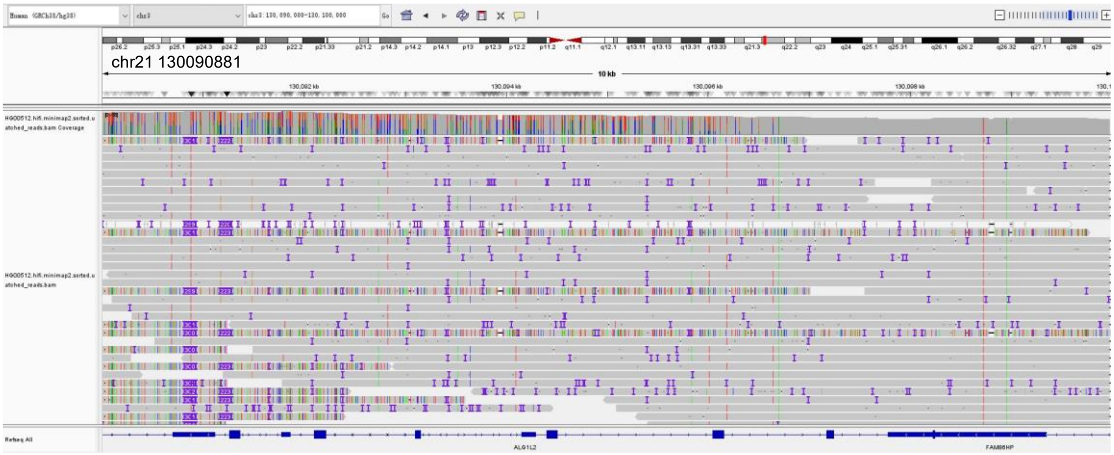

ALU-HG00512-chr4:29,731,596

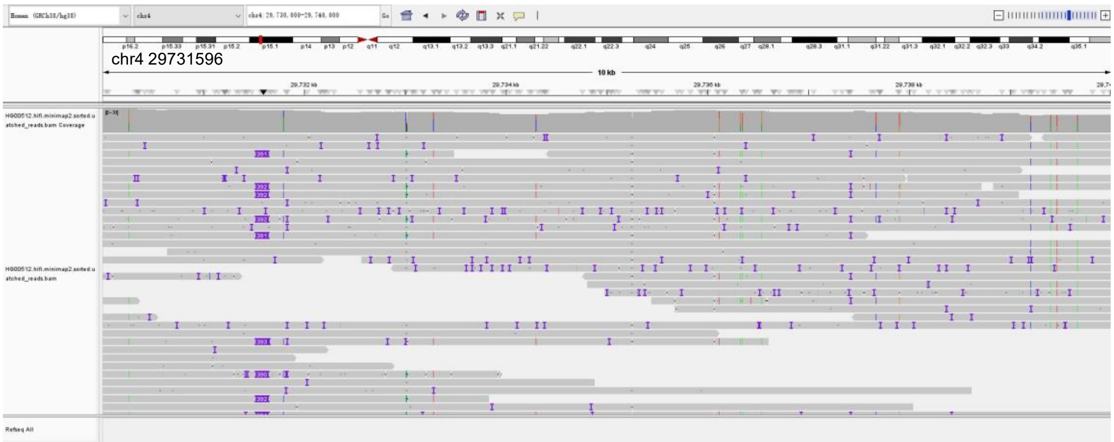


ALU-HG00512-chr5:70,017,992

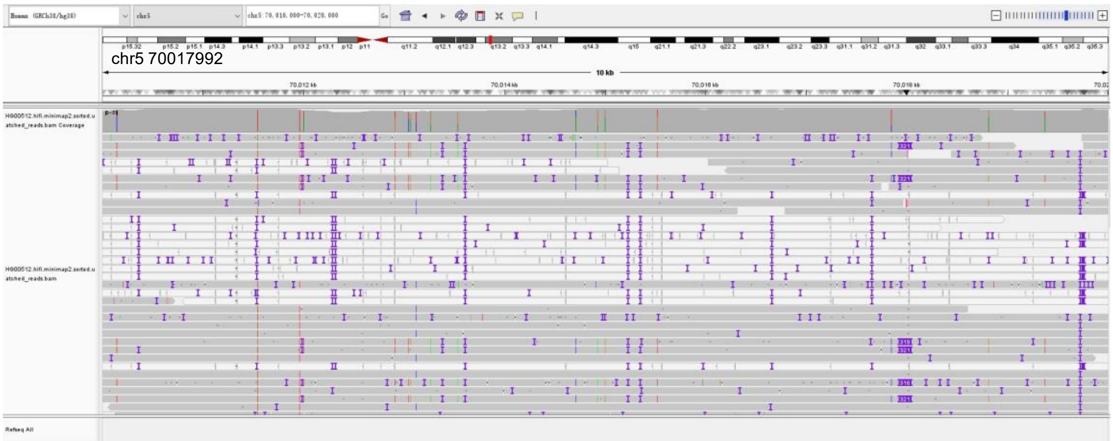


ALU-HG00512-chr5:70,327,057

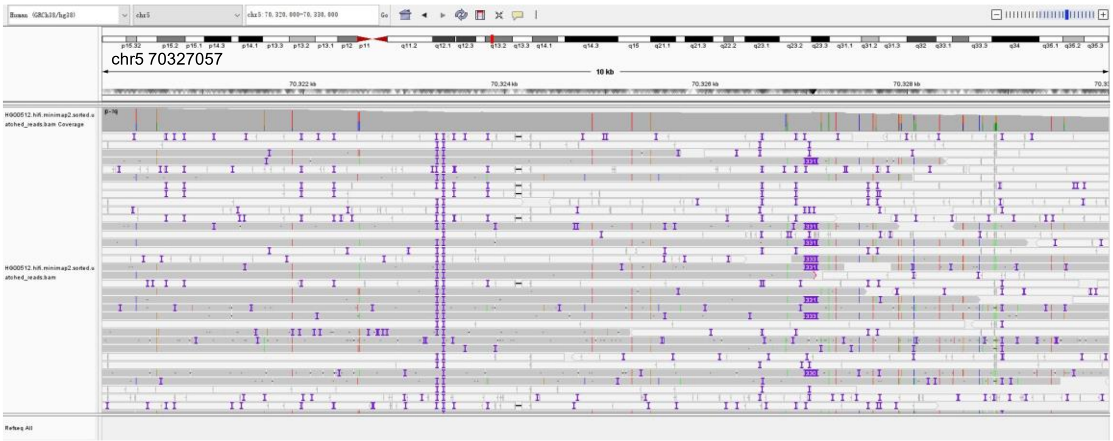

ALU-HG00512-chr9:34,115,382

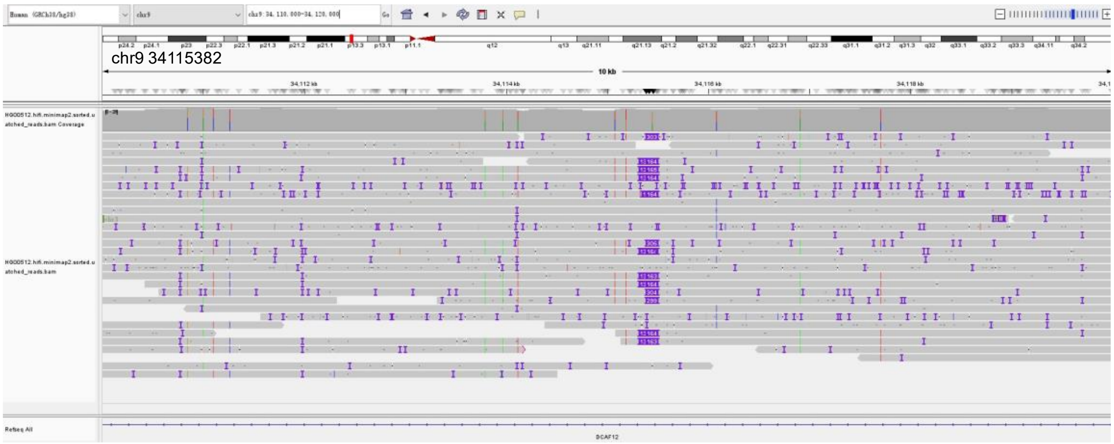


**Supplementary Fig. 6 | IGV validation of supplemented Alu insertions.**

IGV visualizations of long-read alignments supporting Alu insertions detected by MEIsensor but absent from the original benchmark dataset, consistent with bona fide insertion events missed during benchmark construction.

SVA-HG00512-chr19:18,724,785

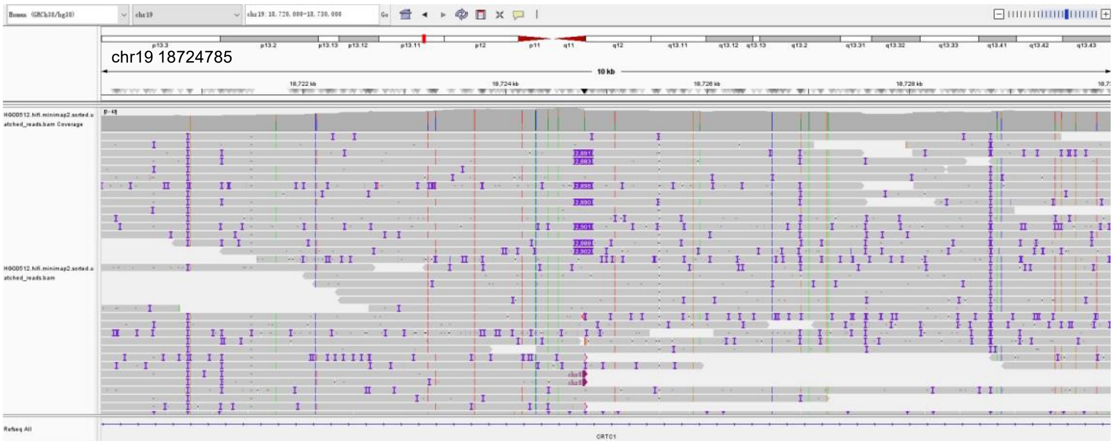


SVA-HG00512-chr5: 726,305

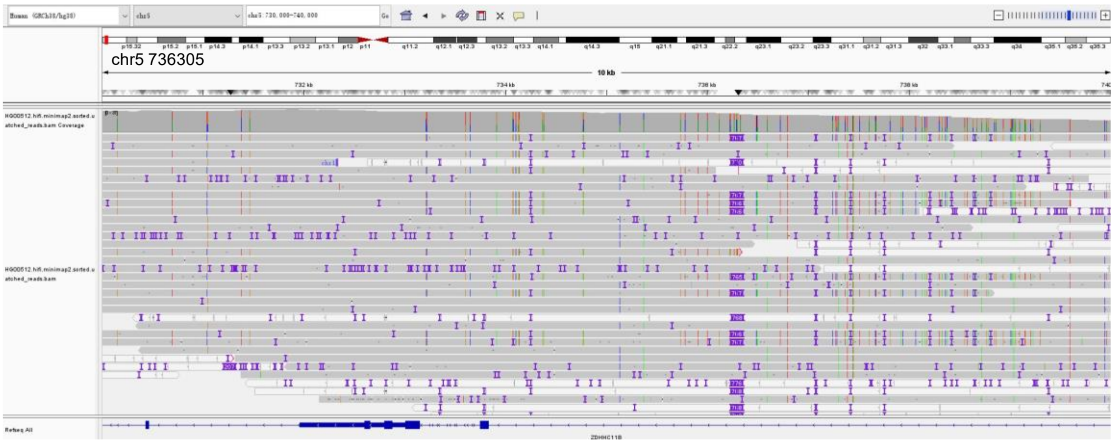


**Supplementary Fig. 7 | IGV validation of supplemented SVA insertions.**

IGV visualizations of long-read alignments supporting SVA insertions detected by MEIsensor but absent from the original benchmark dataset, consistent with bona fide insertion events missed during benchmark construction.

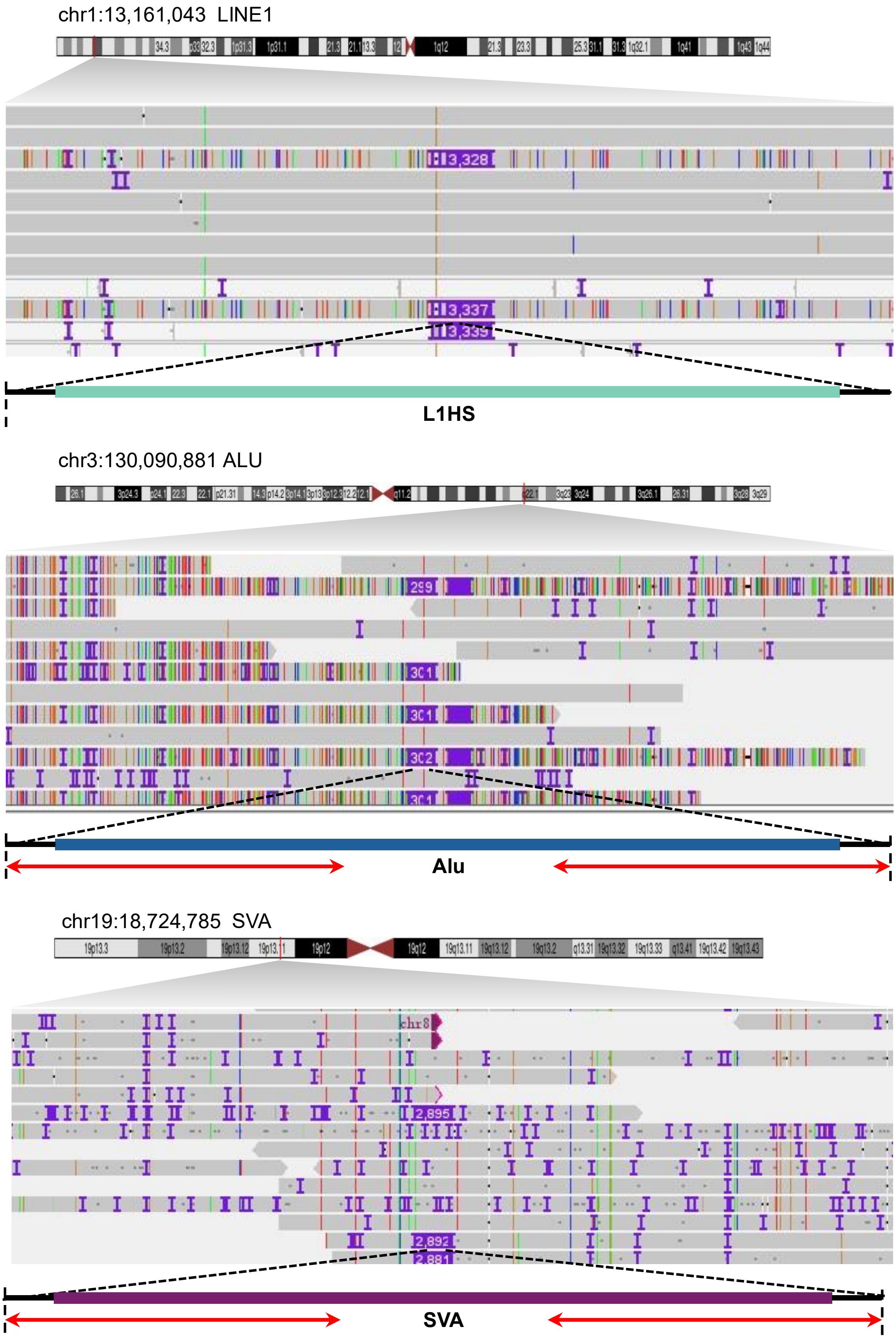


**Supplementary Fig. 8 | Representative benchmark-missing MEIs recovered by MEIsensor.** IGV visualizations show representative LINE1 (top), Alu (middle), and SVA (bottom) insertions that were detected by MEIsensor but absent from the original benchmark. Long-read alignment patterns support these events as bona fide insertions, illustrating how benchmark supplementation recovered previously unannotated MEIs and contributed to improved performance estimates.

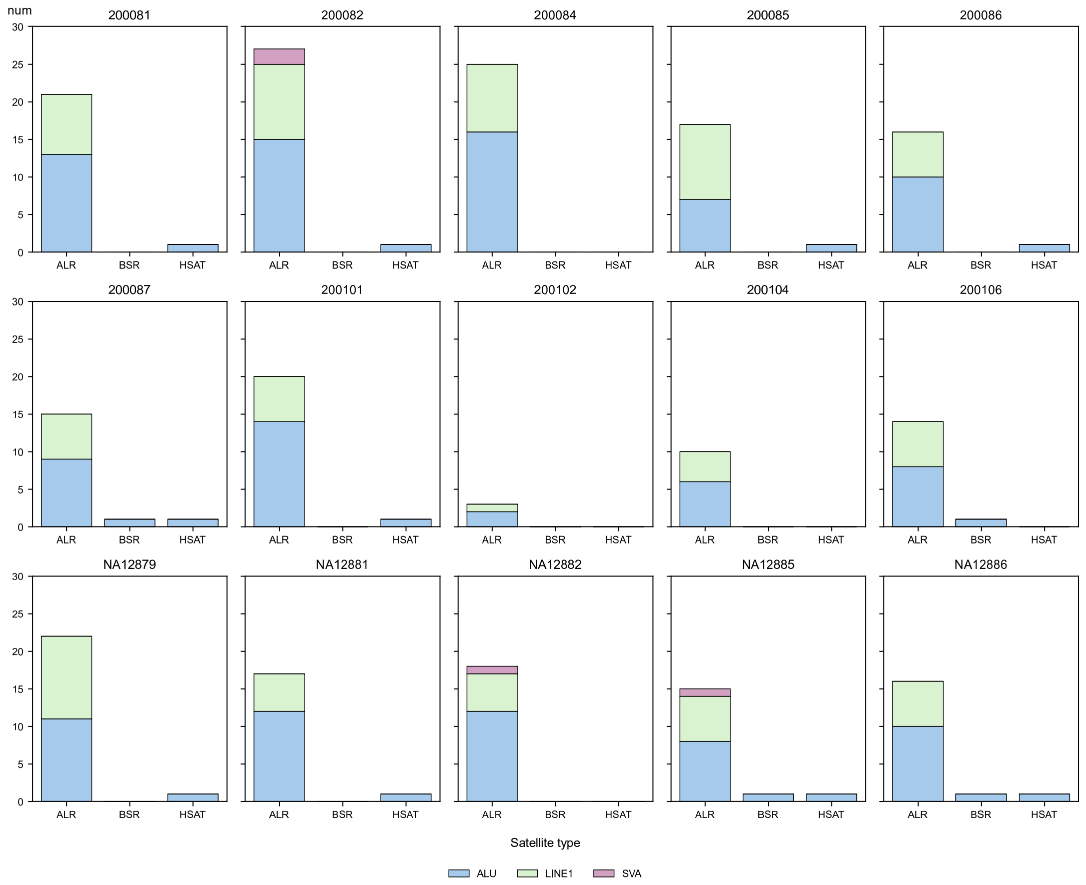


**Supplementary Fig. 9 | Centromeric transposable element insertions are predominantly enriched in α-satellite arrays.** Distribution of transposable element insertions within centromeric regions in selected individuals from the CEPH1463 pedigree. The majority of centromeric insertion variants were located within α-satellite (α-sat) arrays.
